# Chemical-guided SHAPE sequencing (cgSHAPE-seq) informs the binding site of RNA-degrading chimeras targeting SARS-CoV-2 5’ untranslated region

**DOI:** 10.1101/2023.04.03.535453

**Authors:** Zhichao Tang, Shalakha Hegde, Siyuan Hao, Manikandan Selvaraju, Jianming Qiu, Jingxin Wang

## Abstract

One of the hallmarks of RNA viruses is highly structured untranslated regions (UTRs) in their genomes. These conserved RNA structures are often essential for viral replication, transcription, or translation. In this report, we discovered and optimized a new type of coumarin derivatives, such as **C30** and **C34**, which bind to a four-way RNA helix called SL5 in the 5’ UTR of the SARS-CoV-2 RNA genome. To locate the binding site, we developed a novel sequencing-based method namely cgSHAPE-seq, in which the acylating chemical probe was directed to crosslink with the 2’-OH groups of ribose at the ligand binding site. This crosslinked RNA could then create read-through mutations during reverse transcription (i.e., primer extension) at single-nucleotide resolution to uncover the acylation locations. cgSHAPE-seq unambiguously determined that a bulged G in SL5 was the primary binding site of **C30** in the SARS-CoV-2 5’ UTR, which was validated through mutagenesis and in vitro binding experiments. **C30** was further used as a warhead in RNA-degrading chimeras to reduce viral RNA expression levels. We demonstrated that replacing the acylating moiety in the cgSHAPE probe with ribonuclease L recruiter (RLR) moieties yielded RNA degraders active in the in vitro RNase L degradation assay and SARS-CoV-2 5’ UTR expressing cells. We further explored another RLR conjugation site on the E ring of **C30/C34** and discovered improved RNA degradation activities in vitro and in cells. The optimized RNA-degrading chimera **C64** inhibited live virus replication in lung epithelial carcinoma cells.

---

RNA viruses usually have highly structured 5’ and 3’ UTRs in their RNA genome, which can potentially serve as therapeutic targets^1^. In this report, we used SARS-CoV-2 as a specific test-case example and explored to use RNA-degrading chimeras to inhibit virus replication. SARS-CoV-2 is an enveloped ssRNA(+) virus. The whole genome of SARS-CoV-2 (∼30,000 nucleotides) is encoded in a single RNA molecule^2^. The viral RNA in transmitted virions is 5’ capped and 3’ polyadenylated, and, therefore, it is first recognized and treated as mRNA^3^. In this step, the 5’ UTR is used to hijack the host ribosome to translate viral proteins^4^. Furthermore, the 5’ UTR plays an essential role in RNA transcription for each coronavirus structural protein, which is accomplished through a "discontinuous” transcription mechanism. Specifically, the replication transcription complex binds to the 5’ UTR leader transcriptional regulatory sequences (TRS-L), and then “hops” onto the body TRS (TRS-B) sequence located at the 5’-end of each structural gene^5,6^. That said, all SARS-CoV-2 transcripts share the same 5’ UTR leader sequence. In addition, the SARS-CoV-2 5’ UTR was reported essential for viral RNA packaging^7^.

Given the importance of the UTRs in SARS-CoV-2, we and others elucidated the RNA structures in SARS-CoV-2 UTRs^8–12^. The 5’ UTR RNA structures in cell-free buffers, virus-infected cells, and our reporter cell model are highly consistent ^8–13^, suggesting superior stability and suitability serving as drug targets. The 5’ UTR of SARS-CoV-2 contains five stem-loops, namely SL1–5. The start codon resides in SL5, a unique four-way helix^8–12^ (Fig. 1a). SL5 exists in all betacoronovirus species, including MERS and SARS-CoV, and the shapes of this RNA structure are similar^9^. We aligned the SARS-CoV-2 RefSeq and different lineages and demonstrated that the SL5 is highly conserved among all strains^1^. Although a predominant mutation was found in SL5B loop region from recent viral lineages (C241T), this mutation is unlikely to change the overall structure of SL5^1^. In SARS-CoV-2, several structures in the 5’ UTR, including SL4, SL5A, and SL6, were found binding to amilorides^14^ (Fig. 1a). Amilorides demonstrated antiviral activity in SARS-CoV-2 infected cells.

**Fig. 1.**
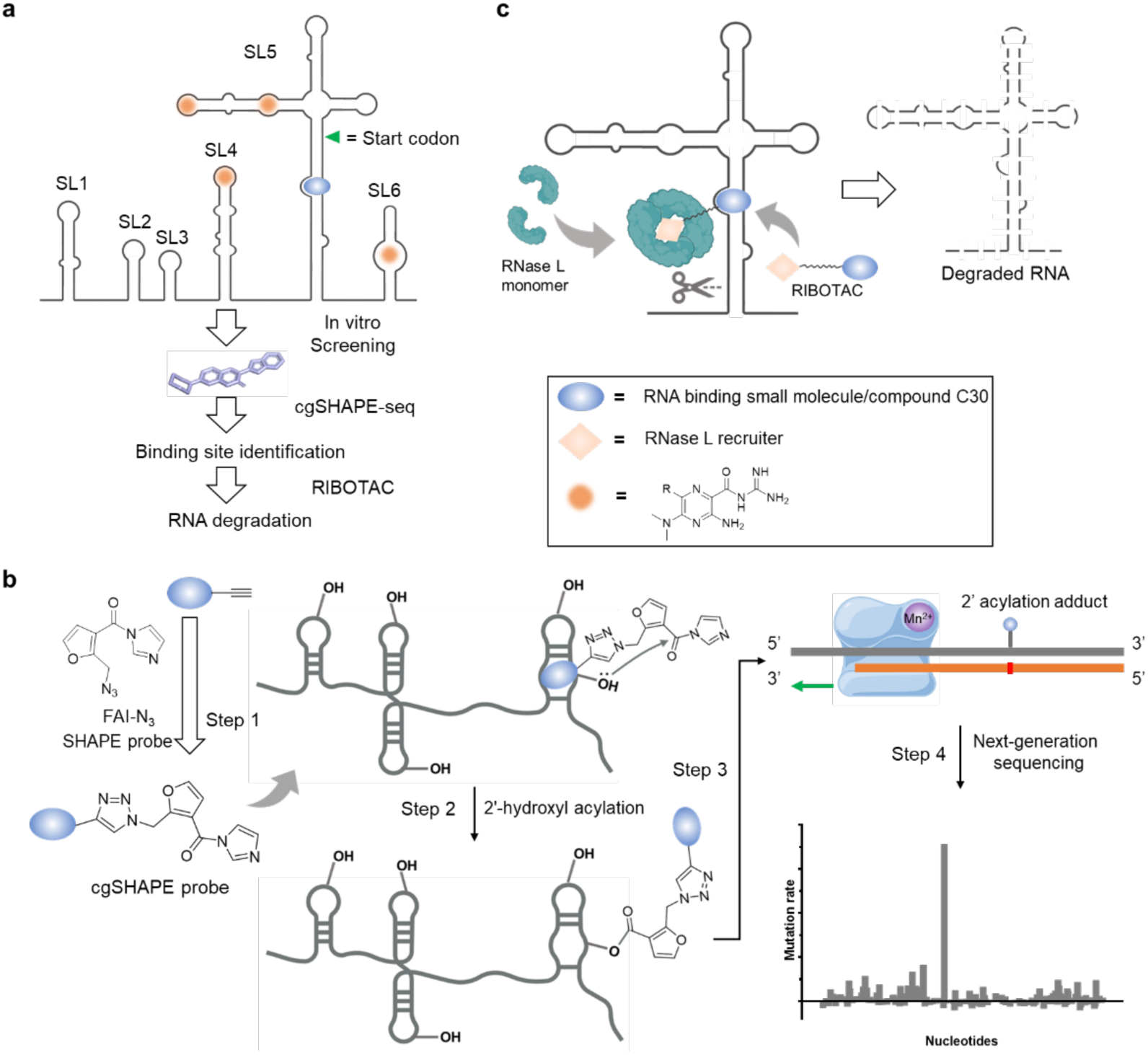
Development of cgSHAPE-seq and anti-viral RIBOTAC. **a**, RNA secondary structures in SARS-CoV-2 5’ UTR and the pipeline in identification of the ligand binding site and the development of RNA-degrading chimeras. **b**, Principle of cgSHAPE-seq for identifying small molecule binding sites in four steps. Step 1: Synthesis of FAI conjugated chemical probe. Step 2: Chemical-guided acylation at the 2’-OH of ribose at the binding site. Step 3: Reverse transcription in the presence of Mn^2+^ creates single-point mutations at the acylation site. Step 4: Mutational profiling and quantification identify the putative binding sites (figure adapted from Figdraw). **c**, RNA-degrading chimeras (RIBOTACs) recruit RNase L at the target RNA to degrade viral RNAs.

Here, we report a pipeline in antiviral discovery and optimization of RNA-degrading chimeras targeting SL5, highlighting a novel sequencing-based method namely chemical-guided (cg) selective 2′-hydroxyl acylation analyzed by primer extension (SHAPE) sequencing (seq), or cgSHAPE-seq, to rapidly locate the RNA ligand binding site (Fig. 1a). First, we screened a small coumarin derivative library in SL5 RNA binding assay and optimized the hit through structure-activity-relationship studies. To elucidate the RNA ligand binding site, we synthesized and applied a new type of chemical probe that can selectively acylate the 2’-OH on the ribose at the location of binding (Fig. 1b)^26,27^. The 2’-OH acylation locations were “recorded” onto RNA molecules by reverse transcriptase as single-point mutations at the modification sites during primer extension. The mutation sites were then captured and deconvoluted by next-generation sequencing^28,29^. Mutational profiling analysis in cgSHAPE-seq unambiguously identified a bulged G in SL5 as the primary binding site in the SARS-CoV-2 5’ UTR. In the literature, other sequencing-based methods were reported using affinity probes bearing nitrogen mustard or diazirine moiety (e.g., ChemCLIP-seq^15–19^ and PEARL-seq^20^). However, a major limitation of these methods is a strong labeling bias toward guanosines^21^. Similar to the SHAPE, cgSHAPE reacts with the 2’-OH group of the ribose in all nucleotides, A, U, G, or C. This can potentially increase the scope and accuracy of proximity-induced chemical reactions on RNAs for mapping purposes. While we were preparing for the manuscript, the Kool lab also reported a proximity-induced acylation approach and determined RNA-binding sites for several FDA-approved drugs^22^.

We further developed RNA-degrading chimeras by replacing the 2’-OH acylating moiety with RNase L recruiter (RLR) moieties on the cgSHAPE probe, as well as by conjugating the RLR moieties on other putative solvent-accessible sites on the RNA ligand. RNA-degrading chimera utilizing endogenous ribonuclease (RNase) L was first reported by the Silverman group in 1993^23^. Recently, the modality of RNA-degrading chimera was further developed by the Disney group and was demonstrated to be active using small-molecule RNA ligands^24–31^ (Fig. 1c). The Disney group also coined the name “ribonuclease targeting chimera or RIBOTAC for this type of RNA degraders. RIBOTACs were shown efficacious to degrade microRNA in cells^25,26^ and mouse models^27^. Importantly, a small-molecule RIBOTAC was recently used to degrade the SARS-CoV-2 RNA genome by targeting an RNA structure named attenuator hairpin near the programmed frameshift (PFS) regulatory element^24^. The viral RNA transcript level was shown to be reduced in a model cell system by ∼50% at 8 μM RIBOTAC^24^. Antisense-based RNA-degrading chimeras targeting spike-protein or envelope-protein encoding RNAs also demonstrated efficacy in SARS-CoV-2 infected cells^29^. Our optimized RIBOTAC robustly degraded SARS-CoV-2 RNA in cellular models at 1 μM and inhibited virus replication at 20 µM in lung epithelial cells. No significant toxicity was observed. Interestingly, we discovered the natural RNase L binding moiety is similar to or less active than the synthetic RLR in the RIBOTAC modality.

## Results

### Chemical optimization of the coumarin derivatives for SL5 RNA binding

We previously synthesized a collection of coumarin derivatives that are known to bind to RNAs^32^. Each of these coumarin derivatives is fluorescent (excitation/emission ∼400/480 nm), enabling us to use fluorescence polarization (FP) to determine the in vitro binding affinity with the RNA receptors. We in vitro transcribed SL5 RNA (144–303, RefSeq NC_045512) and screened the compound library using the FP assay (Extended Data Fig. 1). **C2** binds to SL5 at a dissociation constant (*K*_d_) of 1.45 μM (Table 1). Elimination of the ethyl substituent on the A ring (**C4**) did not improve the binding affinity to SL5 RNA (Table 1). In contrast, a bulky substituent on the A ring (**C6**) impeded the interaction with SL5 (Table 1). This indicated that a non-sterically hindered linker might be suitable for conjugation on ring A without affecting the ligand binding affinity to SL5 RNA. We found **C29** the best ligand in our compound collection, which binds to SL5 RNA at a *K*_d_ of 0.47 μM. Compared to **C4**, **C29** has a different set of substituents on ring E, and therefore, we further investigated ring E while keeping ring A as unsubstituted piperazine (Table 1). In vitro FP assay showed that a fluorinated analog, **C30**, further improves the binding affinity (*K*_d_ = 0.22 µM). Changing the F group into Cl (**C32**) or CF_3_ (**C36**) groups or alternating the fluorinated site (**C31**) all reduced the binding affinities (Table 1). In contrast, replacing the fluorine group in **C30** with methoxy group maintained the binding affinity (**C34**, *K*_d_ = 0.14 µM). To our knowledge, **C30 and C34** are the strongest small-molecule ligand to the SARS-CoV-2 SL5 RNA.

**Table 1.** Structure-activity-relationship study of coumarin derivatives for in vitro binding affinities with SL5 RNA.

| Name | R<sup>1</sup> | R<sup>2</sup> | R<sup>3</sup> | SL5 $K_d$ (μM) |
| --- | --- | --- | --- | --- |
| **C2** | CH<sub>2</sub>CH<sub>3</sub> | NA | NA | 1.45 ± 0.38 |
| **C4** | H | NA | NA | 1.24 ± 0.29 |
| **C6** | CH<sub>2</sub>CH<sub>2</sub>NHBoc | NA | NA | 5.67 ± 4.78 |
| **C29** | NA | H | H | 0.47 ± 0.09 |
| **C30** | NA | H | F | 0.22 ± 0.09 |
| **C31** | NA | F | H | 0.69 ± 0.07 |
| **C32** | NA | H | Cl | 0.42 ± 0.09 |
| **C36** | NA | H | CF<sub>3</sub> | 1.41 ± 0.25 |
| **C34** | NA | H | OMe | 0.14 ± 0.03 |

### cgSHAPE-seq uncovered the bulged G in SL5 as the C30 binding site

We were inspired by a commonly used method for RNA structure elucidation called SHAPE and sought to develop a new chemical-guided sequencing-based method to identify the binding site of **C30/C34** in SL5 RNA. We named the new chemical probing method chemical-guided SHAPE sequencing, or cgSHAPE-seq. Conventional SHAPE uses electrophilic reagents that can form ester adducts on the 2’-OH of the ribose. The unpaired nucleotides have higher accessibility for acylation reactions, which is the basis of structure-based differential acylation activity. Therefore, identification of the acylation site would provide information on RNA base-pairing in the conventional SHAPE. Importantly, the electrophile-ribose adduct can create a mutation during reverse transcription (i.e., primer extension). As a result, conventional SHAPE coupled with quantitative mutational profiling has become a gold standard method to explore RNA topology in vitro and in vivo^33–37^. The key advantage of SHAPE is that the electrophile can usually react with all four nucleotides (A, U, G, or C). We wondered if the ribose acylation could be repurposed for identification of small-molecule binding sites by covalently linking electrophile moieties to RNA-binding chemical ligands.

First, we selected furoyl acylimidazole (FAI) as the acylating electrophile for synthesizing the chemical probe. Compared to other electrophile moieties such as anhydride and acyl cyanide, FAI is more resistant to hydrolysis with a reported half-life of ∼73 min in aqueous solutions^38–40^. The hydrolysis-resistant probe design renders the synthesis and storage less demanding for anhydrous experimental facilities. An azide group on the furoyl moiety was used to provide a click-chemistry handle to conjugate with alkyne-modified **C30** (Fig. 2a). We confirmed that this alkyne group does not eliminate the binding of **C30** to SL5 RNA, even though the binding affinity reduced by 7-fold (Supplementary Information, Fig. S1). After the [3+2] cycloaddition reaction, the carboxylic acid was then converted into an acyl imidazole moiety under a mild condition, finalizing the synthesis of **C30-FAI** probe (Fig. 2a). The acyl imidazole probe was prepared freshly and used directly without further purification (see Method).

**Fig. 2.**
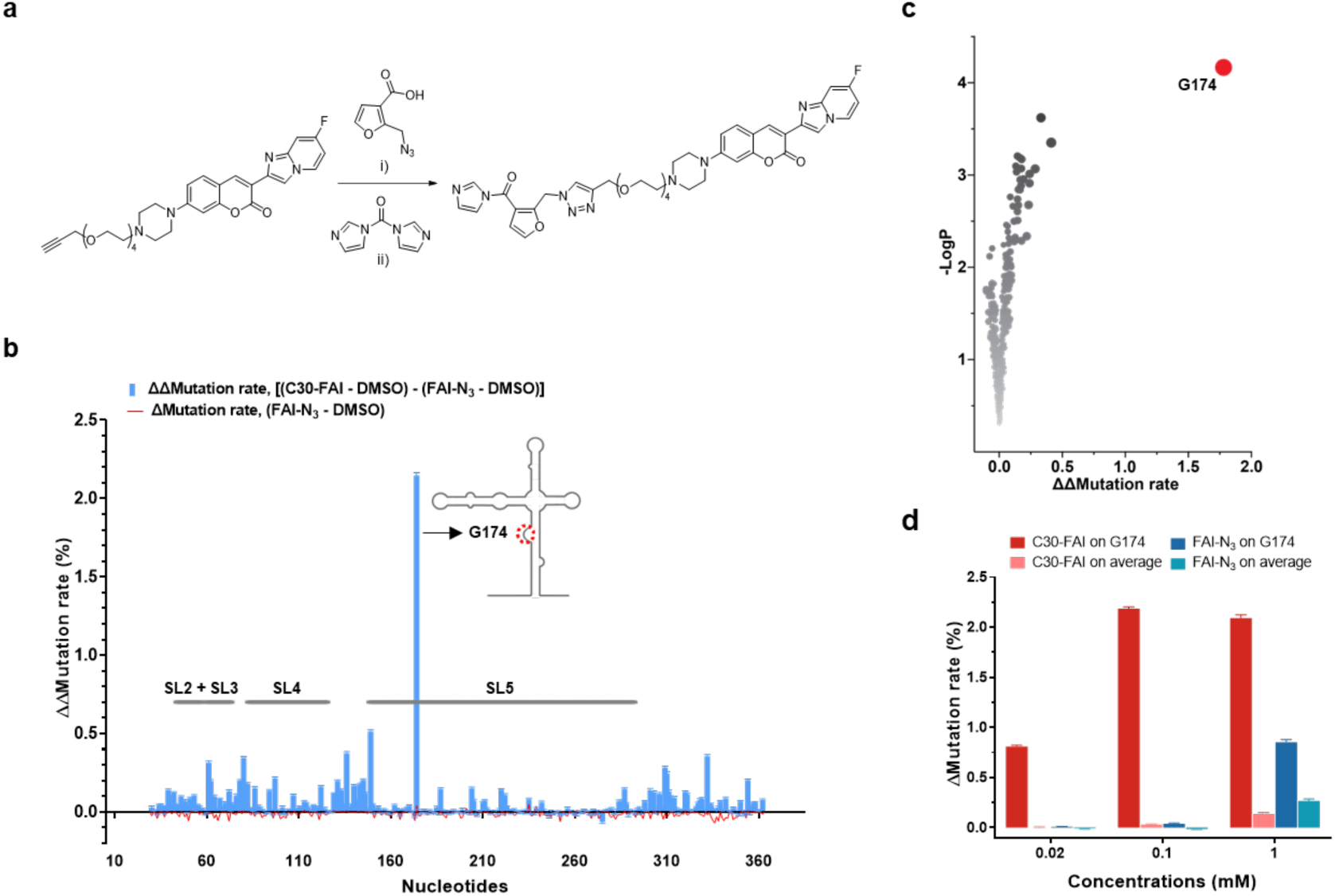
Identification of the binding site by cgSHAPE-seq. **a**, cgSHAPE-seq probe (**C30-FAI**) synthetic route. Reaction conditions: i) tris(hydroxypropyltriazolylmethyl)amine, CuSO_4_, sodium ascorbate, DMSO, room temperature; ii) anhydrous DMSO, room temperature. **b**, cgSHAPE-seq mutational profiling analysis of the SL5 sequence in total RNA extract treated with **C30-FAI** (0.1 mM). Δmutation rate (FAI-N_3_ – DMSO) indicates the background structure-based differential acylation. ΔΔmutation rate [(C30-FAI – DMSO) – (FAI-N_3_ – DMSO)] indicates the proximity-based differential acylation. The cgSHAPE-seq experiments were performed with three replicates (N = 3). **c**, Scatter plot of - LogP vs ΔΔmutation rate. **d**, Comparison of the Δmutation rates of G174 and on average in RNAs treated with different concentrations of **C30-FAI** or FAI-N_3_. Three data points (C75, G216, G285) were removed as outliers as they had an abnormally high mutation rate (*Z*-score > 4.0) in DMSO-treated samples.

Next, we applied **C30-FAI** in SARS-CoV-2 5’ UTR RNA to elucidate the binding site of **C30**. It is known that conventional SHAPE experiments usually require high concentrations (10–100 mM) of acyl imidazole (e.g., FAI) for ribose acylation. To avoid obtaining structure-based SHAPE activity caused by FAI moiety alone, we chose to use a much lower dose of the chemical probe for cgSHAPE-seq. We reasoned that at low concentrations of the probe (0.02–1 mM), the differential acylation activity would be predominantly caused by ligand binding (i.e., proximity-promoted acylation). Briefly, the total RNA was extracted from SARS-CoV-2 5’ UTR expressing cells (see Method) and refolded in buffer. **C30-FAI**, FAI-N_3_, or DMSO was individually reacted with the folded RNA for 15 min at 37 °C. After the reaction, we used ProtoScript II reverse transcriptase (New England Biolabs) in the presence of MnCl_2_ (3 mM) for primer extension using a protocol modified from the literature report (see Method)^33^. In this step, we screened several commercially available reverse transcriptases and found ProtoScript II one of the best enzymes that can tolerate Mn^2+^ in the reaction (Supplementary Information, Fig. S2). The cDNA was then amplified by PCR in the SARS-CoV-2 5’ UTR region and the resulting amplicon was subsequently sequenced. We applied an existing software package ShapeMapper2 developed by the Weeks group for mutational profiling analysis^33,39^. We calculated the background Δmutation rate (FAI-N_3_ – DMSO) for each nucleotide and pleasingly observed a very low background signal at 0.02 and 0.1 mM of FAI-N_3_, indicating that the structure-based differential acylation activity is negligible in cgSHAPE-seq at these concentrations (Extended Data Fig. 2). In contrast, FAI-N_3_ at 1 mM significantly increased the Δmutation rate, implying the contribution of structure-based SHAPE activities started to emerge at high concentrations, confirming the suitable probe concentration for cgSHAPE is in the range 0.02−0.1 mM (Fig. 2d). We then calculated the RNA ligand-induced ΔΔmutation rate [(C30-FAI – DMSO) – (FAI-N_3_ – DMSO)] for each nucleotide and identified G174 as the only significantly mutated nucleotides in 0.02–0.1 mM probe-treated samples (Fig. 2b and 2c, Extended Data Fig. 3). In 1 mM probe-treated samples, the cgSHAPE signal of G174 is comparable with 0.1 mM probe-treated ones, but a high signal-to-noise ratio was observed due to the structure induced SHAPE background (Fig. 2d). Mapping the nucleotide with previously identified secondary structures uncovered that G174 is a single-nucleotide bulge in the SL5 stem region^8–12^. It is worth noting that a previous study showed that G174 was highly reactive in a regular SHAPE experiment using another acylation agent, NAI, at 100 mM^11^. This concentration is at least 1,000 times higher than the optimal cgSHAPE probe concentration determined in our study (0.02−0.1 mM), and therefore, the cgSHAPE signal is unlikely attributed to the structure-induced SHAPE background at this position (Fig. 2d).

We then validated the **C30** binding site in SL5 by testing individual substructures of the SL5 RNA. The loop region of SL5A, SL5B, and a minimized four-helix junction, named SL5^M^ (containing shorter stems) were synthesized chemically or enzymatically (Fig. 3a). The in vitro binding results demonstrated that only SL5^M^ retained similar binding affinity to **C30** (Fig. 3a, Extended Data Fig. 4). To further validate the putative binding site G174 in SL5^M^, we designed and synthesized SL5^M^ RNAs with different mutations that disrupt the bulged G or other RNA structures (Fig. 3b). As expected, deletion of G174 or base-pairing G174 with an additional C both resulted in a 7-fold decrease in binding affinity to **C30**. Replacement of G174 with A, C, or U also significantly reduced **C30** binding. We also expanded the bulged G by inserting different nucleotides between C173 and U175 or between A270 and G271, all these mutated RNAs demonstrated 5–6-fold reduced binding affinity to **C30**. These results suggested the importance of a single bulged G in accommodating C30’s binding. Changing the closing U-A base pair into C-G (3’-end of G174) or C-G base pair into U-A (5’-end of G174) also resulted in a 4-fold decreased binding. Mutations on other parts of the RNA have less impact (i.e., within 2-fold) in changing the binding affinity to **C30** (Fig. 3b). Altogether, these observations validated that the bulged G region is the primary binding site in SL5 RNA for **C30**. We concluded that cgSHAPE-seq is a validated method for identifying the binding site of RNA-binding small molecules.

**Fig. 3.**
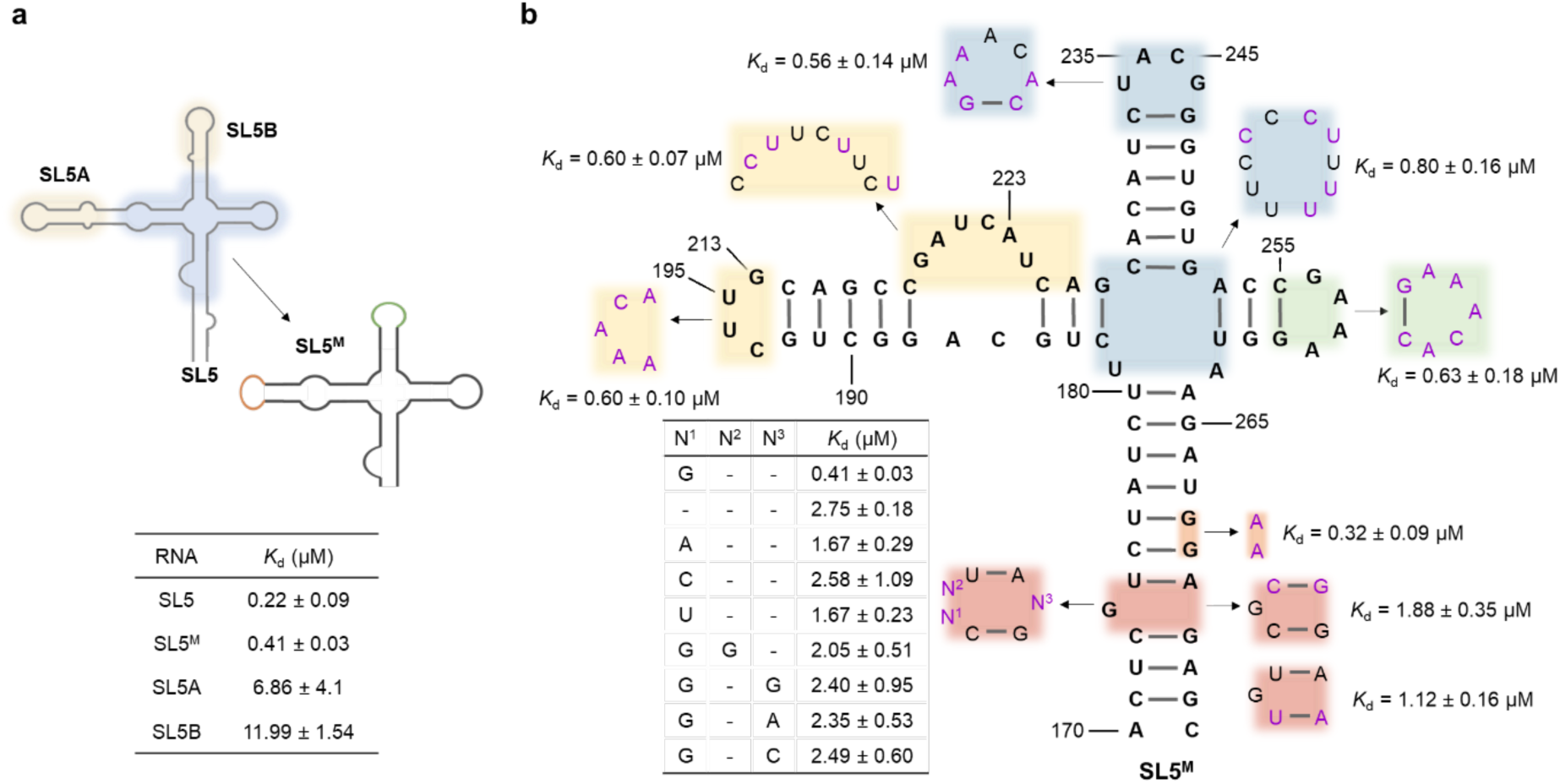
Validation of the binding site. **a**, Structural fragments of SL5 and their binding affinities to **C30**. **b**, SL5^M^ mutants and their binding affinities to **C30**.

### Comparison of cgSHAPE-seq and other sequencing-based RNA ligand localization methods

We compared cgSHAPE-seq with two existing sequencing-based methods for determining RNA ligand binding sites, PEARL-seq^20^ and Chem-CLIP^15–19^. PEARL-seq used photocrosslinking moieties such as diazirine to covalently link the bound RNA nucleotides. On the other hand, Chem-CLIP uses a nitrogen mustard moiety to alkylate nucleobases by nucleophilic reactions. To compare these two approaches with cgSHAPE-seq, we synthesized three new C30 probes: both C30-D and C30-BD contain a diazirine group for photocrosslinking, and C30-NM contains a nitrogen mustard group for alkylation. We also confirmed that these probes have similar binding affinity to SL5 RNA compared to the cgSHAPE probe (Supplementary Information, Fig. S1 and S3). These three chemical probes were used to treat the total RNA extracted from SARS-CoV-2 5’ UTR expressing cells at 0.1 mM under appropriate crosslinking conditions (see Supplementary Information). We used the same reverse transcription and analysis pipeline as those used in our cgSHAPE-seq experiments. We observed that C30-D in PEARL-seq can also identify G174 as the binding site albeit with a higher background than cgSHAPE-seq, while the other two probes didn’t significantly differentiate G174 from other modified nucleotides (Extended Data Fig. 5). Taken together, we concluded that the FAI-based acylation probe is a more sensitive chemical probe to identify RNA binding sites of small molecules.

### Comparison of two RNase L recruiting moieties in RIBOTACs

We then conjugated **C30/C34** with RLRs to synthesize SL5-targeting RNA degraders (RIBOTACs). For RLR conjugation on the A ring, the acylating moiety in the above cgSHAPE probe (**C30-FAI**) was replaced with RLR moieties. We noticed that in most reported co-NMR structures of RNA bulges and small molecules that are similar to **C30/C34**, both termini of the small molecules are solvent accessible^41–43^, and, therefore, ring E was also explored for conjugation to RLRs, where the methoxy group of **C34** was substituted for RLR attachment (Fig. 4a).

**Fig. 4.**
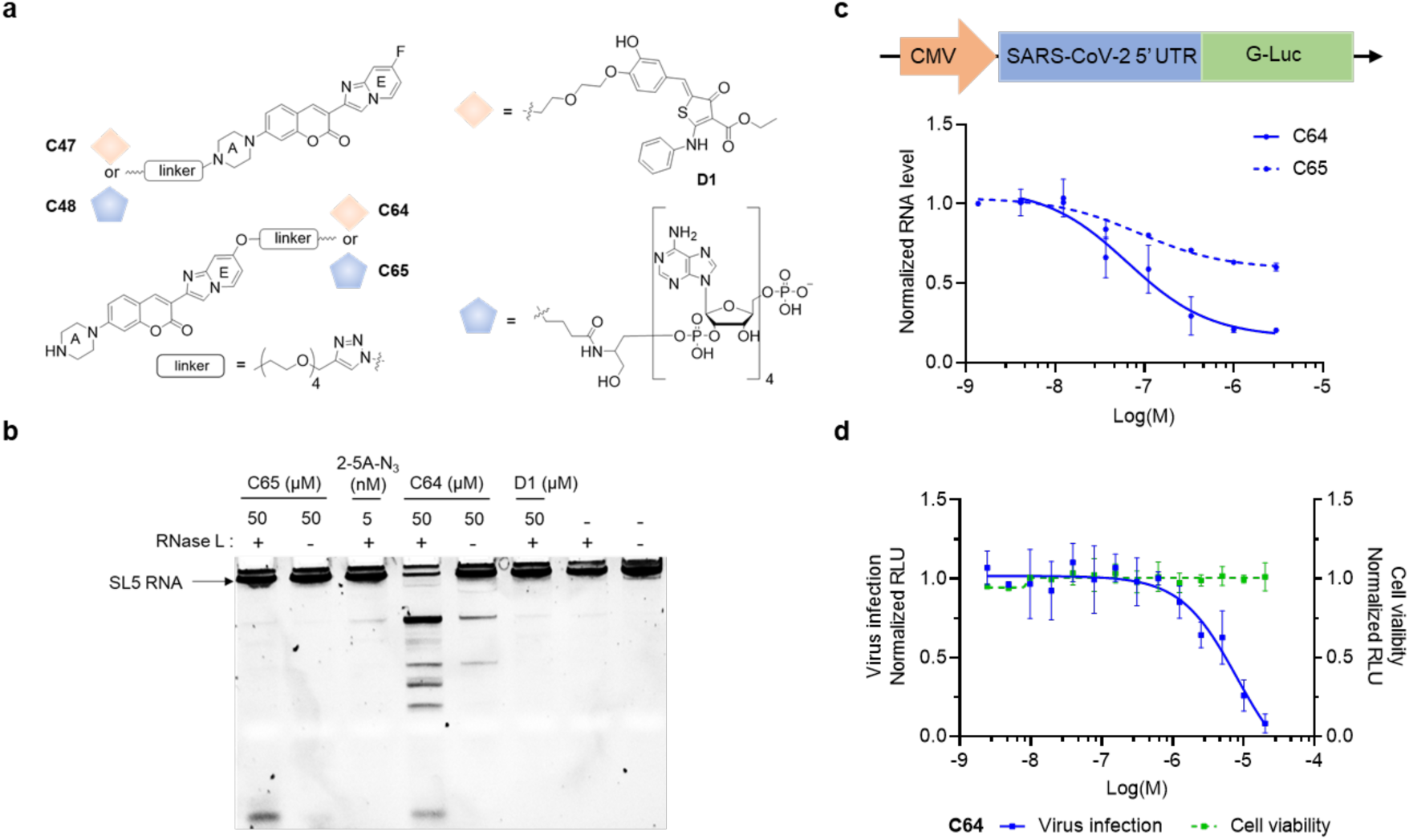
RNA degrading activity and anti-viral activity of C30-based RIBOTACs. **a**, Synthesis of **C30**-based RIBOTACs using conjugation sites on rings A or E of **C30**. **b**, Comparison of two RLR moieties in the RIBOTAC modality using the in vitro RNase L degradation assay with purified SL5 RNA. **c**, Cellular activity of RIBOTACs in SARS-CoV-2 5’ UTR expressing cells. **d**, Inhibitory effect of RIBOTAC **C64** in SARS-CoV-2 infected A549 cells. The cytotoxicity of the compound was also evaluated. The dose-response curves are representative of three independent measurements (N = 3).

For RLR moieties, the natural RNase L ligand 2’-5’-lined oligoadenylate (2-5A) and its synthetic mimic **D1** were both previously reported to be used in RNA-degrading chimeras (Extended Data Fig. 6)^24,25^. Combining the two conjugation sites and two RLR structures, we obtained four RIBOTAC candidates, **C47**, **C48**, **C64**, and **C65**, for SL5 RNA degradation (Fig. 4a). We validated that the polyethylene glycol (PEG) linker on 2-5A does not affect the activity in a reported RNase L degradation assay with a 5’ 6-fluorescein-tagged model RNA containing multiple RNase L cleavage sites^44,45^ (Extended Data Fig. 6a). It was demonstrated that the binding affinity between RNase L and **D1** (*K*_d_ ≈ 18 μM) is 80,000-fold weaker than that observed for 2-5A^45^. Consistent with this reported in vitro binding data, the synthetic **D1** alone is > 10,000 times weaker than 2-5A in the in vitro RNase L degradation assay (Extended Data Fig. 6b and 6c).

Next, we tested the four RIBOTACs in the in vitro RNase L degradation assay with purified SL5 RNA and observed their activities in order: **C64** > **C47** ≈ **C48** > **C65** (Fig. 4b, Extended Data Fig. 7a). To our surprise, the RIBOTAC **C64** with **D1** as the RLR moiety is much stronger than **C65** with 2-5A at 50 μM (Fig. 4b). This result is contrary to what we would have predicted based on the activities of the RLR moieties per se. We validated these in vitro findings in SARS-CoV-2 5’ UTR expressing 293T cells. In this cell model, the SARS-CoV-2 5’ UTR sequence was fused to a CMV promoter-controlled Gaussia luciferase expression cassette (Fig. 4c; for sequences, see Method). Consistent with the RNase L degradation assay result, the maximum potency of **C64** (i.e., RNA reduction level) was significantly better than **C65** (Fig. 4c). The activities of **C47** and **C48** in this cell model are similar to those of **C64** and **C65** (Extended Data Fig. 7b).

### Efficacy of RIBOTAC in live virus infection assay

Finally, we tested the activity of **C64** in SARS-CoV-2 infected cells. The SARS-CoV-2 virus was engineered to include a Nano Luciferase (NLuc) reporter by fusing NLuc onto ORF7 of the SARS-CoV-2 genome^46^. In this way, the NLuc signal is proportional to the viral protein copy number in cells. We applied a human lung epithelial carcinoma cell line A549 that expresses high level of ACE2 as the host cell^46^. The cells were infected with the SARS-CoV-2-NLuc virus at a multiplicity of infection (MOI) of 2.0 at 1 h before the treatment with RIBOTACs **C64** for 3 d. To our satisfaction, **C64** showed > 95% inhibition at 20 μM (Fig. 4d). At the same concentration, no major toxicity is observed in A549 cells (Fig. 4d).

## Discussion

In cgSHAPE-seq, it is critical to employ a concentration at which the predominant source of SHAPE activity is the proximity-enhanced activity. Specifically, a standard SHAPE probing should be performed parallelly with an identical probe concentration as that of the cgSHAPE probe to determine the full set of SHAPE reactivity in the folded RNA (Fig. 2d). At the optimal cgSHAPE probe concentration (i.e., 0.02−0.1 mM in this study), the standard SHAPE activity should be minimal (typically < 0.1% as demonstrated in Fig. 2d). In addition, the putative binding sites determined by cgSHAPE-seq require extensive validation. In the current study, we used mutagenesis and in vitro binding assay to validate the binding site (Fig. 3). We envision that reverse genetics with a mutated binding site coupled with relevant functional assays can also be used for validation purposes if the RNA binder has well-defined biological consequences.

Multiple crucial factors merit attention when designing cgSHAPE probes: (1) the cgSHAPE probe should be considerably stable in water solution to allow sufficient target engagement and to avoid hydrolysis by moisture in the air during the experiment preparation steps. In this current study, we chose FAI moiety with a reported half-life of ∼73 min. We attempted to use C30-NAI conjugate but observed that the probe is considerably less stable than **C30-FAI** (data not shown). (2) The conjugation site of acylation moiety should be solvent accessible and not impede ligand binding. In this current study, the **C30-FAI** precursor for cgSHAPE and **C30-D** probe for PEARL-seq both showed significant binding to the RNA target, albeit 3−7 fold less binding affinity (Supplementary Information, Fig. S1 and S3). (3) The functional group should be compatible with acylating moiety. For example, the FAI-based probes used in this report would not be compatible with nucleophilic RNA ligands^14^ due to self-reaction. cgSHAPE-seq potentially has a limitation with bias towards certain types of small molecule-RNA interactions. As shown in conventional SHAPE, FAI moiety has a higher reactivity towards unpaired RNA nucleotides. Although most of the reported RNA ligands target the unpaired region^47,48^, cgSHAPE-seq may be less reactive for ligands that bind to the double-stranded RNA grooves.

In our cgSHAPE-seq result, apart from G174, we also observed a cluster of nucleotides from A131 to G149 showing slightly higher mutation rates than others (Fig. 2b and 2c). This can be potentially caused by the nonspecific binding of **C30** with flexible sequence^32^. RNA targeting strategies are known to have off-target effects due to shallow binding sites on RNAs and relatively weak binding affinity for small molecules. **C64** at 3 μM can cause >100 genes up- or down-regulated (|Log2FoldChange| > 2) in the transcriptome (Extended Data Fig. 8, Supplementary Table S1). The activity of the RIBOTAC might be improved if a more potent and selective RNase L recruiter is used^49^. Specifically, we showed that the natural RNase L recruiter/activator 2-5A, which is negatively charged, is sometimes not compatible with the positively charged RNA binder **C30** (Fig. 4b). For this reason, new synthetic RNase L recruiter should probably be considered to be neutral or positively charged as most of the reported RNA ligands are also positively charged^50–52^.

In summary, we developed a new generalizable chemical probing method called cgSHAPE-seq for quickly identifying small molecule-RNA binding sites by sequencing. cgSHAPE probes react with the 2’-OH groups on the ribose close to the binding sites with a mitigated dependency of the nucleobase identity observed in other reported methods. We used cgSHAPE-seq to identify a bulged G on SL5 as the primary binding site on the SARS-CoV-2 5’ UTR targeted by the newly discovered coumarin derivatives, such as **C30** and **C34**. Finally, we developed a novel **C30/C34**-based RNA degrader (RIBOTAC) capable of degrading viral RNA transcripts in cells and inhibiting virus replication in SARS-CoV-2 infected cells, offering crucial insights into RNA degrading chimeras’ design.

## Methods

### Synthesis of C30-FAI (cgSHAPE probe)

Compound **C30-alkyne** (50 mg, 0.08 mmol) in DMSO (1 mL) was added 2-(azidomethyl)furan-3-carboxylic acid (15 mg, 0.09 mmol), THPTA (9 mg, 0.02 mmol), sodium ascorbate (8 mg, 0.04 mmol) and CuSO_4_ (3 mg, 0.02 mmol). The reaction vial was sealed, evacuated, and refilled with N_2_ three times and stirred at room temperature overnight. DMSO was removed under vacuum and the residue was purified by silica gel column chromatography (0 – 10% CH_3_OH in CH_2_Cl_2_) to afford **C30-FCA** as a yellow solid (45 mg, 70%). MS-ESI (*m*/*z*) [M+1]^+^ 746.28.

^1^H NMR (500 MHz, DMSO-*d*_6_) δ 8.71 – 8.68 (m, 2H), 8.52 (s, 1H), 8.09 (s, 1H), 7.71 – 7.68 (m, 2H), 7.38 (dd, *J* = 10.1, 2.6 Hz, 1H), 7.02 (dd, *J* = 9.0, 2.3 Hz, 1H), 6.97 (td, *J* = 7.6, 2.6 Hz, 1H), 6.88 (d, *J* = 2.4 Hz, 1H), 6.72 (d, *J* = 1.9 Hz, 1H), 5.92 (s, 2H), 4.52 (s, 2H), 3.58 – 3.51 (m, 14H), 3.38 (t, *J* = 5.1 Hz, 4H), 2.59 (t, *J* = 5.1 Hz, 4H), 2.56 (t, *J* = 5.8 Hz, 2H).

^13^C NMR (126 MHz, DMSO-*d*_6_) δ 160.2 (d, *J* = 252 Hz), 159.5, 154.9, 153.3, 152.2, 144.4, 144.3, 143.4, 139.4, 138.6, 129.6, 129.2 (d, *J* = 11.8 Hz), 124.2, 114.5, 112.1, 111.7, 111.4, 110.2, 104.2 (d, *J* = 29.8 Hz), 99.6, 99.4, 69.8, 69.7, 69.1, 63.4, 57.0, 54.9, 52.6, 46.7, 44.8.

Compound **C30-FCA** (45 mg, 0.06 mmol) in anhydrous DMSO (0.6 mL) was added carbonyldiimidazole (CDI, 10 mg, 0.06 mmol) and the reaction mixture was stirred at room temperature for 1 h. The reaction mixture contains ∼75% **C30-FAI** and ∼25% unreacted **C30-FCA** (see Supplementary Information) and was used directly in RNA modification. The stock solution was used as **75 mM** and can be stored at –80 °C for long-term storage.

### cgSHAPE-seq using Total RNA Extract from Cells

SARS-CoV-2 5’ UTR expressing cells were harvested and pelleted. Total RNA was extracted using TRIzol Reagent (Invitrogen) per the user’s manual. An on-column DNA digestion was performed to remove the residual genomic DNA in total RNA using DNase I (10 U/μL, Roche) and RDD buffer (Qiagen). Purified total RNA was dissolved in water and stored at –80 °C before use. For RNA modification, 5 μg total RNA was used for each reaction. **C30-FAI** and FAI-N_3_ were prepared at 20 mM, 2 mM, and 0.4 mM in DMSO as 20× working solution. Briefly, total RNA was added water and 5× folding/reaction buffer (500 mM HEPES pH 7.4, 500 mM KCl, 30 mM MgCl_2_) to make a 47.5 μL solution. The solution was incubated at 37 °C for 30 min to refold. 2.5 μL **C30-FAI** (cgSHAPE probe), FAI-N_3_ (background control) or DMSO was added to the total RNA and mixed well by pipetting. The mixture was incubated at 37 °C for 15 min and then quenched by adding RLT buffer (Qiagen). The RNA was then extracted using RNeasy kit (Qiagen). 500 ng total RNA was used for reverse transcription and then PCR as described below. All reactions were performed in triplicates.

For reverse transcription (10 μL reaction), probe or DMSO treated RNA and reverse transcription primer (0.5 μM in final reaction buffer) were heated at 70 °C for 5 min and snap-cooled on ice for 1min. 5× reaction buffer (375 mM Tris-HCl, 500 mM KCl, 15 mM MnCl_2_, pH 7.4, 2 μL), DTT (100 mM, 1 μL), dNTP (10 mM, 0.5 μL), ProtoScript II (0.5 μL, New England Biolabs, M0368L) and RNase inhibitor (0.2 μL, ApexBio, K1046) were added. The reaction was incubated at 42 °C for 1 h and deactivated at 70 °C for 15 min. In each PCR reaction (50 μL), cDNA (2.5 μL) was mixed with Phire Hot Start II DNA Polymerase (Thermo Fisher, 1 μL), dNTP (10 mM, 1 μL), 5× Phire Green reaction buffer (10 μL), primers (0.5 μM in final reaction buffer) and water (35.5 μL). After reaction, the amplicon was purified using a DNA Clean & Concentrator kit (Zymo Research) following the user’s manual. The purified DNA was submitted for next-generation sequencing (Amplicon-EZ, Azenta Life Sciences).

An integrated software package developed by Busan and Weeks, ShapeMapper2 was used to analyze the fastq files for mutational profiling and the rresult was used to generate Figures 2b, 2c, and Extended Data Fig. 339. The reference sequence (SARS-CoV-2_5_UTR.fa) required for ShapeMapper2 is listed below.

>SARS-CoV-2_5_UTR

aggtttataccttcccaggtaacAAACCAACCAACTTTCGATCTCTTGTAGATCTGTTCTCTAAACGAACTTTAAAATCTGTGTGGCTGTCACTCGGCTGCGTGCTTAGTGCACTCACGCAGTATAATTAATAACTAATTACTGTCGTTGACAGGACACGAGTAACTCGTCTATCTTCTGCAGGCTGCTTACGGTTTCGTCCGTGTTGCAGCCGATCATCAGCACATCTAGGTTTCGTCCGGGTGTGACCGAAAGGTAAGATGGAGAGCCTTGTCCCTGGTTTCAACGAGGGAGTCAAAGTTCTGTTTGCCCTGATCTGCATCGCTGTGGCCGAGGCCAAGCCCACCGAGAACAACGAagacttcaacatcgtggccg. (lowercase = primer binding sequences).

### *In Vitro* RNase L Degradation Assay

Purified recombinant GST-tagged RNase L was purchased from MyBioSource (MBS1041064). The buffer of RNase L was exchanged into a buffer containing 50 mM Tris-HCl (pH 7.4) and 100 mM NaCl using Zeba Desalting Column (Thermo Fisher, 8766) using the manufacturer’s protocol. For RNase L degradation of SL5 RNA, T7 transcribed SL5 RNA was first purified by polyacrylamide gel electrophoresis (PAGE) and recovered using small-RNA PAGE Recovery Kit (Zymo Research, R1070). RNase L (1.3 µg in 5 µL) was incubated in the presence of **C47**, **C48**, **C64**, **C65**, or DMSO control in the cleavage buffer (final reaction volume is 8 µL) at 4 °C for 12 h. The 1X cleavage buffer contains 25 mM Tris-HCl (pH 7.4), 10 mM MgCl_2_, 100 mM KCl, 50 µM ATP, and 7 mM β-mercaptoethanol. The SL5 RNA (120 ng in 2 µl H_2_O) was then added into the reaction mixture and incubated for another 2 h at 22 °C. The reaction was stopped by adding RNA Gel Loading Dye (Thermo Fisher, R0641) at 1:1 ratio. The samples (4 µl) were then loaded on TBE-urea polyacrylamide gel (20%) for electrophoresis (180 V for 85 min). The gel was stained with SYBR safe (1/50,000, ApexBio, A8743) in TBE buffer for 1 min and visualized on gel imager (Thermo Fisher, iBright FL1500).

The RNA sequence of SL5 used in this assay is: 5’- UCGUUGACAGGACACGAGUAACUCGUCU AUCUUCUGCAGGCUGCUUACGGUUUCGUCCGUGUUGCAGCCGAUCAUCAGCACAUCUA GGUUUCGUCCGGGUGUGACCGAAAGGUAAGAUGGAGAGCCUUGUCCCUGGUUUCAACG A.

For RNase L degradation of a model 6-FAM-tagged RNA, RIBOTACs in the above protocol were replaced with **D1** (0.37 µg in 0.75 µl DMSO) or **2-5A-N_3_** (0.12 ng in 0.75 µl H_2_O)^44,45^. After electrophoresis, the gel was not stained and was directly visualized on the gel imager at the 6-FAM fluorescence channel. The 6-FAM RNA (5’-6-FAM-UUAUCAAAUUCUUAUUUGCCCCAUU UUUUUGGUUUA-BHQ) was purchased from IDT.

### SARS-CoV-2 5’ UTR Expressing Stable Cell Line

293T cells (Thermo Fisher, R70007) were cultured in DMEM growth medium (Gibco, 11995040) supplemented with 10% FBS (Cytiva, SH30910.03) and 1% Antibiotic-Antimycotic (Gibco, 15240062) at 37 °C in 5% CO_2_ atmosphere. For producing the lentivirus, 293T cells were seeded in a 6-well plate (Fisher, FBO12927) at 3 x 10^5^ cells per well and transfected with 1 µg of SARS-CoV-2 5’ UTR expressing lentivirus vector (pLV-SARS-CoV-2-5’UTR-GLuc) along with the packaging plasmids pMD2.G (0.4 µg) and psPAX2 (0.6 µg) using Lipofectamine 2000 (Invitrogen, 11668019). At 24 h post-transfection, the cell medium was replaced with fresh growth medium. 48 h after the change of media, the supernatant containing the lentivirus particles was siphoned and centrifuged at 500 g for 10 min at 4 °C to remove the cell debris. The virus particles were further concentrated at 10X in volume using Lenti-X-concentrator (Clontech, PT4421-2) according to the manufacturer’s protocol. The lentivirus can be quantified using literature method^53^. Usually, 10^7^–10^8^ plaque forming units (pfu)/mL lentivirus was obtained after the concentrator treatment. For lentiviral transduction, 293T cells were inoculated with the concentrated viral suspension (multiplicity of infection ∼10) using polybrene (Sigma-Aldrich, TR-1003-G) at a final concentration of 8 μg/mL. At 24 h post-transduction, the culture medium was replaced with fresh growth media. After recovery for 24 h, the transduced cells were then selected in blasticidin (10 μg/mL, Invivogen, ant-bl) for 2 weeks. For stable single clone selection, the cells were diluted in the growth medium containing blasticidin (10 μg/mL) to a final density of 1 cell per 100 µL. The diluted cell suspension was then dispensed to a 96-well plate (100 µL per well). The plate was incubated at 37 °C for 4 weeks. A single cell colony from one of the wells was then selected for experiments.

pLV-SARS-CoV-2-5UTR-Luc was constructed by inserting the SARS-CoV-2 5’ UTR and Gaussian luciferase into the pLV vector, under the control of the CMV promoter. The insert sequence is as follows:

attaaaggtttataccttcccaggtaacaaaccaaccaactttcgatctcttgtagatctgttctctaaacgaactttaaaatctgtgtggctgtcactcggctgcgtgcttagtgcactcacgcagtataattaataactaattactgtcgttgacaggacacgagtaactcgtctatcttctgcaggctgcttacggtttcgtccgtgttgcagccgatcatcagcacatctaggtttcgtccgggtgtgaccgaaaggtaagatggagagccttgtccctggtttcaacgagggagtcaaagttctgtttgccctgatctgcatcgctgtggccgaggccaagcccaccgagaacaacgaagacttcaacatcgtggccgtggccagcaacttcgcgaccacggatctcgatgctgaccgcgggaagttgcccggcaagaagctgccgctggaggtgctcaaagagatggaagccaatgcccggaaagctggctgcaccaggggctgtctgatctgcctgtcccacatcaagtgcacgcccaagatgaagaagttcatcccaggacgctgccacacctacgaaggcgacaaagagtccgcacagggcggcataggcgaggcgatcgtcgacattcctgagattcctgggttcaaggacttggagcccatggagcagttcatcgcacaggtcgatctgtgtgtggactgcacaactggctgcctcaaagggcttgccaacgtgcagtgttctgacctgctcaagaagtggctgccgcaacgctgtgcgacctttgccagcaagatccagggccaggtggacaagatcaagggggccggtggtgactaa (lowercase = SARS-CoV-2 5’ UTR; uppercase = Gaussia luciferase; underline = start codon).

### Quantitative Reverse Transcription PCR (RT-qPCR) Assay

The SARS-CoV-2 5’ UTR expressing cells were seeded at 3 x 10^5^ cells per well in 12-well plates in 1 mL growth medium at 37 °C for 3 h. The cells were then treated with the compounds (**C47**, **C48**, **C64**, or **C65**) at various concentrations (1.3 nM–3 µM) for 48 h. After treatment, the supernatant was aspirated from each well and the total RNA was then extracted from the cells using RNeasy mini kit (Qiagen, 74104). The total RNAs were quantified by ultraviolet absorption at 260 nm (Thermo Fisher, NanoDrop 1000). Usually 10–20 μg total RNA was obtained from each well. cDNAs were synthesized from 500 ng of total RNA for each sample using M-MLV reverse transcriptase (Promega, M1701) and (dT)_25_ according to the manufacturer’s protocol. 1 µl of cDNA mixture was used in a 15 µl RT-qPCR reaction (Apex-Bio, K1070). The human GAPDH RNA level was used as the reference for normalization. The RT-qPCR primer sequences used for the PCR are shown below:

SL5-SYBR-FW: 5’-CGTTGACAGGACACGAGTAA

SL5-SYBR-RV: 5’-TTGAAACCAGGGACAAGGCTC

GAPDH-FW: 5’-GACAAGGCTGGGGCTCATTT

GAPDH-RV: 5’-CAGGACGCATTGCTGATGAT

### SARS-CoV-2 Inhibition Assay

Vero-E6 cells (ATCC® CRL-1586™) and A549 cells (ATCC® CCL-185) were cultured in Dulbecco’s modified Eagle’s medium (DMEM, Cytiva Life Science, SH30022) with addition of 10% fetal bovine serum (FBS, Millipore Sigma, F0926) at 37°C under 5% CO_2_ atmosphere. A549 cells were transduced with a human ACE2-expressing lentivirus vector, and the transduced were cultured in the DMEM plus 2 µg/µL puromycin^46^.

#### Virus and titration

SARS-CoV-2-Nluc was created by engineering the nanoluciferase (Nluc) gene into the OFR7 of the SARS-CoV-2 genome. The insertion site of Nluc at ORF7 was based on previous mNeonGreen reporter SARS-CoV-2^54^. The virus was propagated in Vero-E6 cells once, aliquoted in DMEM, and stored at –80°C. A biosafety protocol to work on SARS-CoV-2 in the BSL3 Lab was approved by the Institutional Biosafety Committee of the University of Kansas Medical Center.

#### Plaque assay^55,56^

Vero-E6 cells were seeded in 24-well plates at a density of 0.5 ×10^6^ cells per well. A virus stock was serially diluted at 10-fold in Dulbecco’s phosphate-buffered saline, pH7.4 (DPBS). 200 µL of the diluent were added to each well and incubated for 1 h on a rocking rotator. After removing the virus diluent, 0.5 mL of overlay media (1% methylcellulose in DMEM with 5% FBS) were added to each well. The plates were incubated at 37 °C under 5% CO_2_ for 4 days. The methylcellulose overlays were aspirated, and the cells were fixed with 10% formaldehyde solution for 30 min and stained with 1% crystal violet solution followed by extensive washing. Plaques in each well were counted and multiplied by the dilution factor to determine the virus titer at pfu/mL.

#### Determination of half-maximal inhibitory concentration (IC_50_)^46,57^

ACE2-A549 cells were seeded into 96 well plates. When the cells were confluent, SARS-CoV-2-NLuc viruses were diluted with cold PBS and added into each well at a multiplicity of infection (MOI) of 2 (2 pfu/cell). The plates were kept in the CO_2_ incubator for 1 hour. Compound **C64** was diluted at 2 x serials from 20 µM to 0.002 µM. The virus-PBS solution was aspirated. Each well was washed with cold PBS three times, and was loaded with the diluted compounds. Each concentration was loaded in triple wells in the plates, and the total volume of each well was 0.2 mL. The plates were kept in the incubator. After 3 days post-infection, the culture media were aspirated from each well and the wells were washed with PBS three times. The nano-luciferase activity assay (Promega, N1110) was carried out following the manufacturer’s instructions. Briefly, 100 ul of cell lysis buffer were added to each well for 10 minutes to completely lyse the cells. Then 100 ul of nano-luciferase reaction reagent were added to each well and the luminescent signal was determined at A490 absorbance on a plate reader (Bio-Tek, Synergy). The IC_50_ was calculated using GraphPad Prism 8.0 software.

### Statistical Analysis

All data shown as means ± s.d. with sample size (N) listed for each experiment. Statistical analysis was carried out with Prism GraphPad 8.0. Unpaired two-sample t-tests were used to analyze significant differences between the group means. The P values were calculated by Prism GraphPad 8.0 or R. For data generated from ShapeMapper2, the standard error (stderr) associated with the mutation rate at a given nucleotide in the S (probe treated) or U (DMSO treated) samples was calculated as: stderr 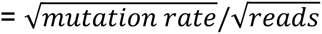. The standard error of the Δmutation rate at a given nucleotide is: 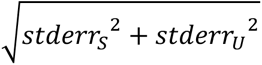.

## Data Availability

The cgSHAPE-seq data for **C30-FAI**, FAI-N_3_ or DMSO-treated total RNAs, and RNA-seq data for **C64**-treated cells were deposited in NCBI SRA with accession numbers PRJNA1029650 and PRJNA947619, respectively.

## Acknowledgments

Research reported in this article was supported by the National Institute of General Medical Sciences (NIGMS) of the National Institutes of Health (NIH) under award numbers R35GM147498 and P20GM113117, W. M. Keck Foundation, and the University of Kansas General Research Funds. We also thank Dr. Robert Silverman at Cleveland Clinic for providing the authentic 2-5A samples for comparison. We have obtained SARS-CoV-2-Nluc from Drs. Shi and Menachery through The University of Texas Medical Branch (UTMB)’s World Reference Center for Emerging Viruses and Arboviruses. We thank the University of Kansas Medical Center Genomics Core for doing RNA-seq experiment. (The KUMC Genomics Core was supported by Kansas Intellectual and Developmental Disabilities Research Center (NIH U54 HD 090216), the Molecular Regulation of Cell Development and Differentiation COBRE (P30 GM122731-03), the NIH S10 High-End Instrumentation Grant (NIH S10OD021743) and the Frontiers CTSA grant (UL1TR002366)).

## Author contributions

J.W. conceived the work. J.W., Z.T., S. Hegde and J.Q. wrote the paper. Z.T and M.S. performed chemical synthesis. Z.T. and J.W. conducted the RNA sequencing and analysis. S. Hegde collected RNA degradation data in vitro and in cells. S. Hao and J.Q. performed the live virus assay.

---

**Extended Data Fig. 1.**
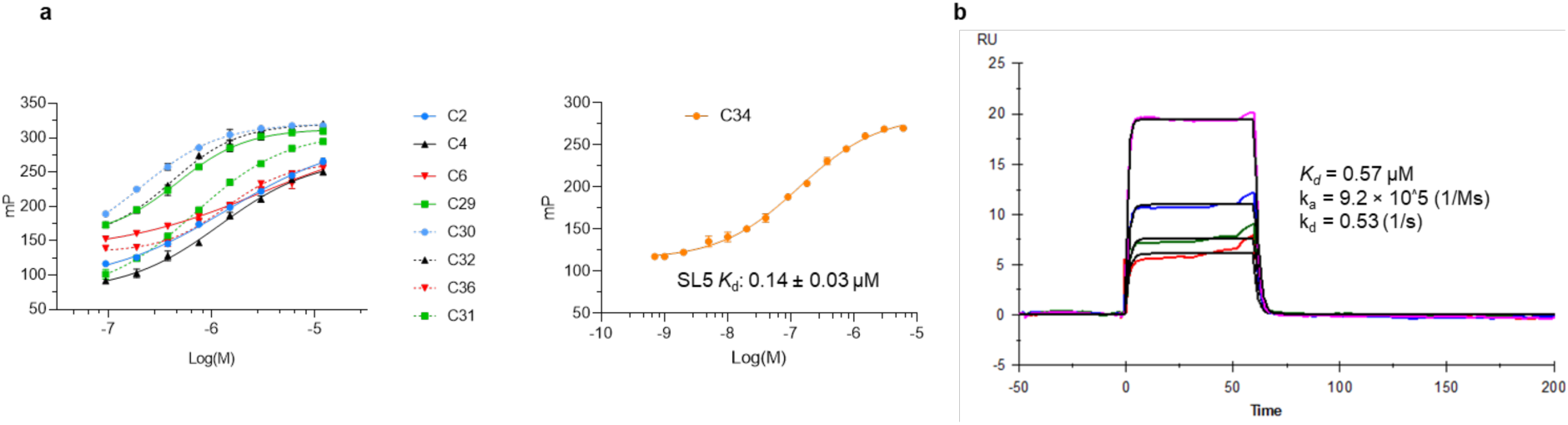
Binding affinity of coumarin derivatives to SL5 RNA. **a**, Dose-response curves of coumarin derivatives in fluorescence polarization assay with in vitro transcribed SL5 RNA. All compounds were used at a concentration of 80 nM. Each data point represents the mean fluorescence polarization value of two independent replicates (N = 2). **b**, Surface plasmon resonance (SPR) binding analysis of **C30** with SL5 RNA. (Curve fitting is shown as black line. [SL5 RNA] = 0.075, 0.15, 0.3, and 0.6 μM).

**Extended Data Fig. 2.**
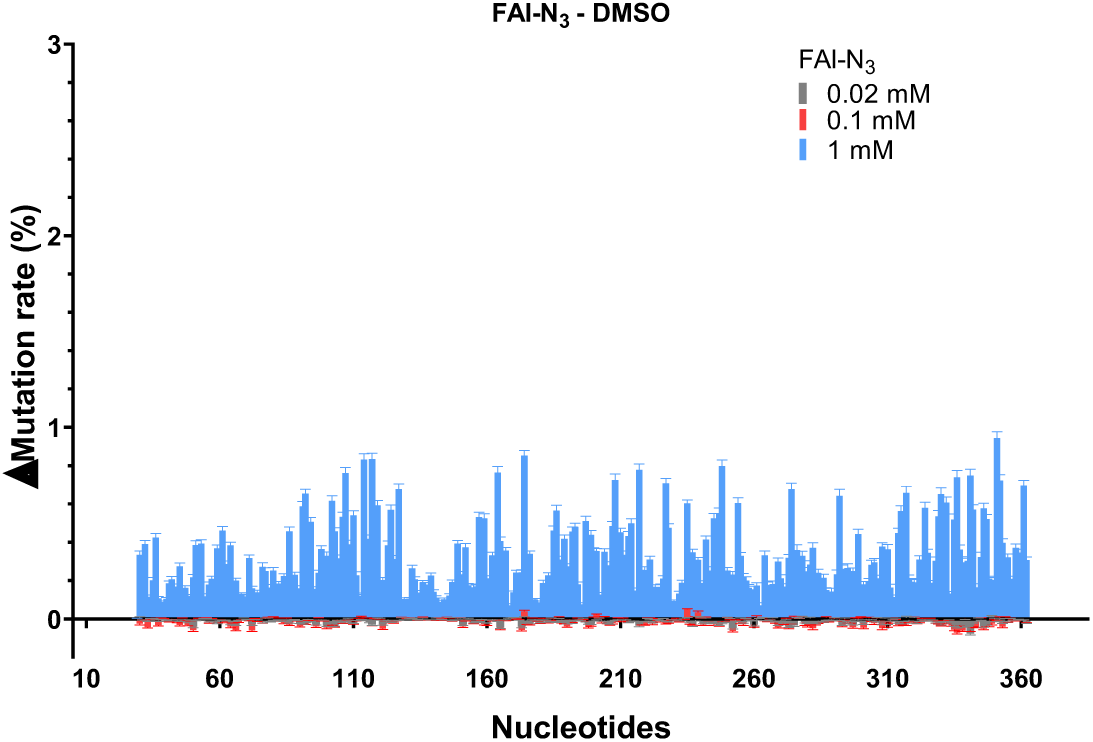
Background Δmutation rate (FAI-N_3_ – DMSO) of the SL5 sequence in total RNA extract treated with different concentrations of FAI-N_3_. Three data points (C75, G216, G285) were removed as outliers as they had an abnormally high mutation rate (*Z*-score > 4.0) in DMSO-treated samples.

**Extended Data Fig. 3.**
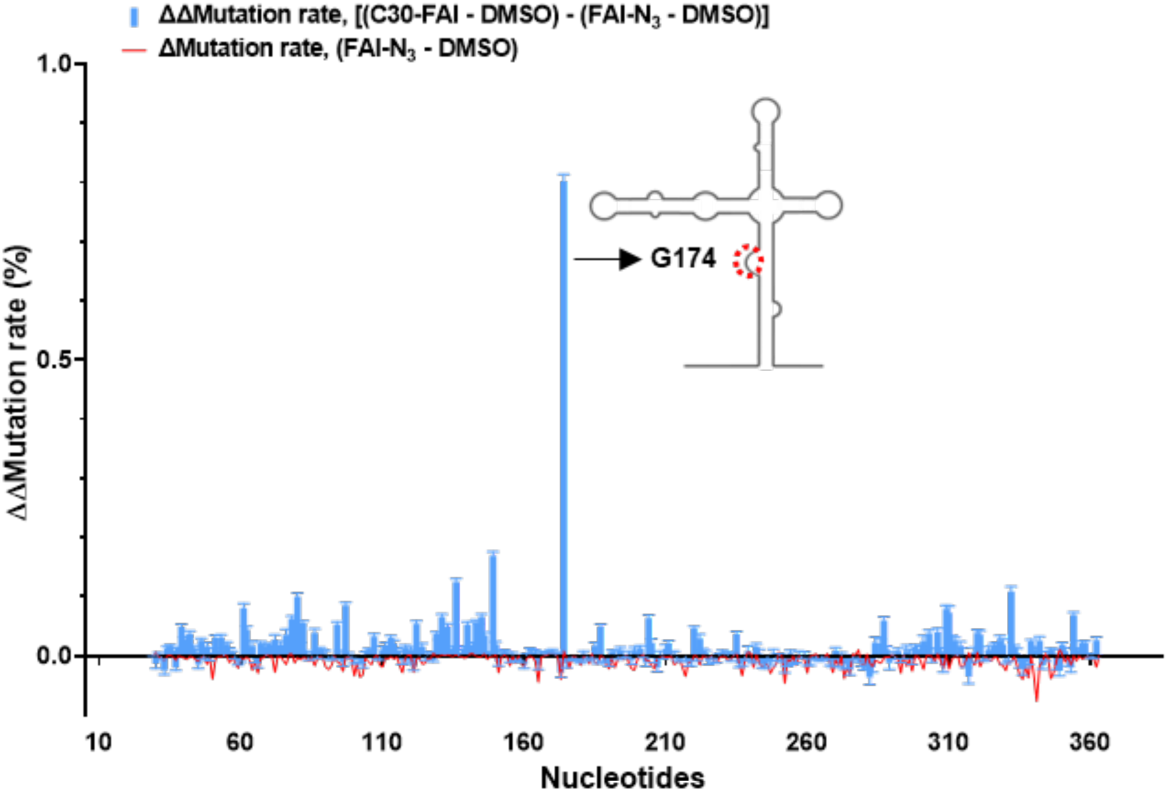
cgSHAPE-seq mutational profiling analysis using C30-FAI (0.02mM). Δmutation rate (FAI-N_3_ – DMSO) indicates the background structure-based differential acylation. ΔΔmutation rate [(C30-FAI – DMSO) – (FAI-N_3_ – DMSO)] indicates the proximity-based differential acylation. The cgSHAPE-seq experiments were performed with three replicates (N=3). Three data points (C75, G216, G285) were removed as outliers as they had an abnormally high mutation rate (*Z*-score > 4.0) in DMSO-treated samples.

**Extended Data Fig. 4.**
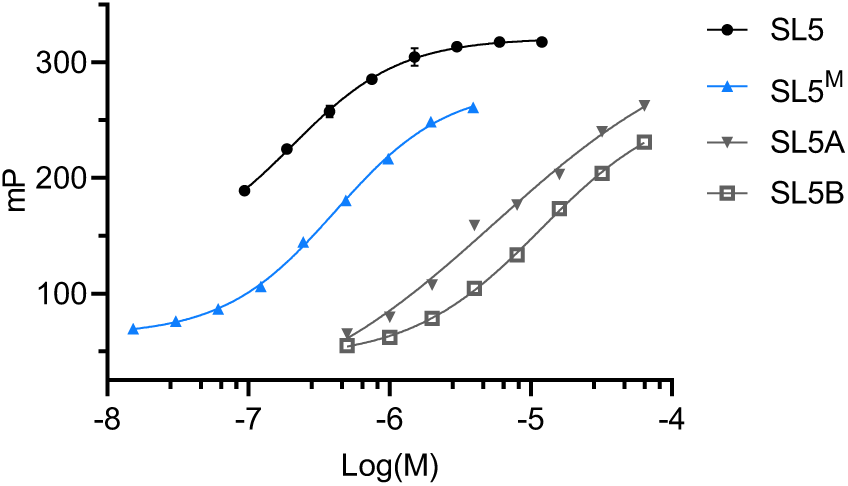
Dose-response curves of **C30** (80 nM) in fluorescence polarization assay with SL5 RNA and its substructures. Each data point represents the mean fluorescence polarization value of two independent replicates (N = 2).

**Extended Data Fig. 5.**
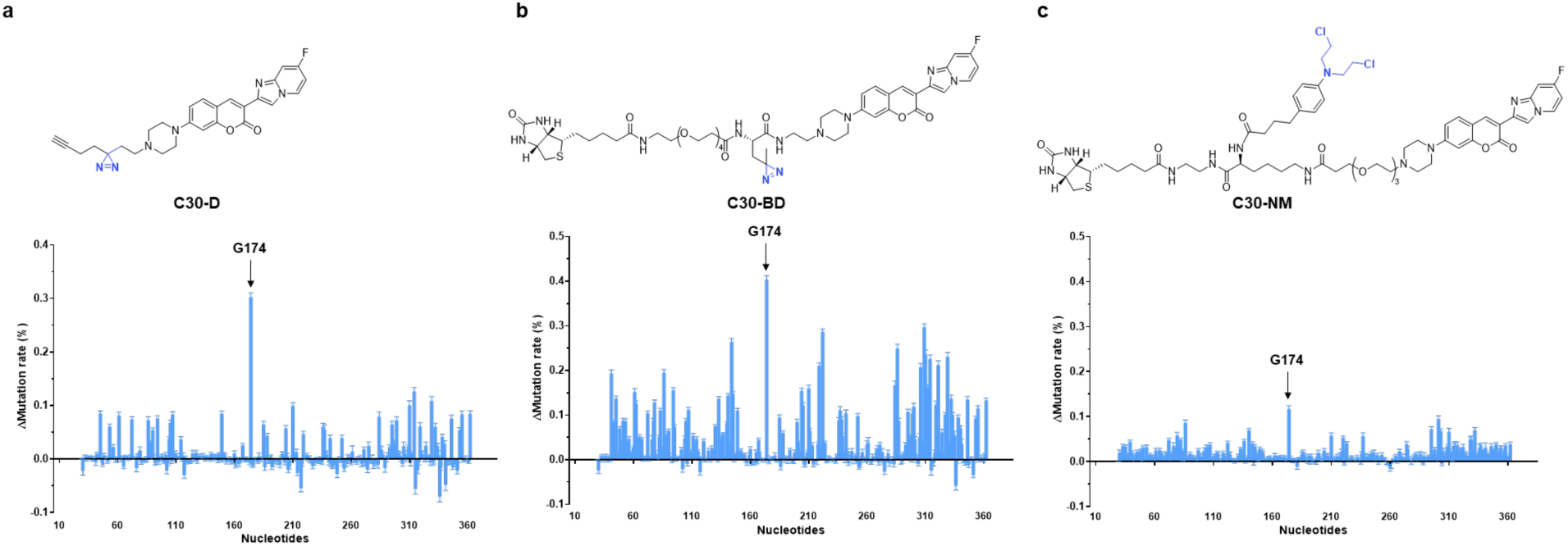
Mutational profiling analysis of crosslinking probes. **a,** Structure of C30-D and mutational profiling analysis. **b,** Structure of C30-BD and mutational profiling analysis. **c,** Structure of C30-NM and mutational profiling analysis. All experiments were performed with three replicates (N = 3). Three data points (C75, G216, G285) were removed as outliers as they had an abnormally high mutation rate (*Z*-score > 4.0) in DMSO-treated samples.

**Extended Data Fig. 6.**
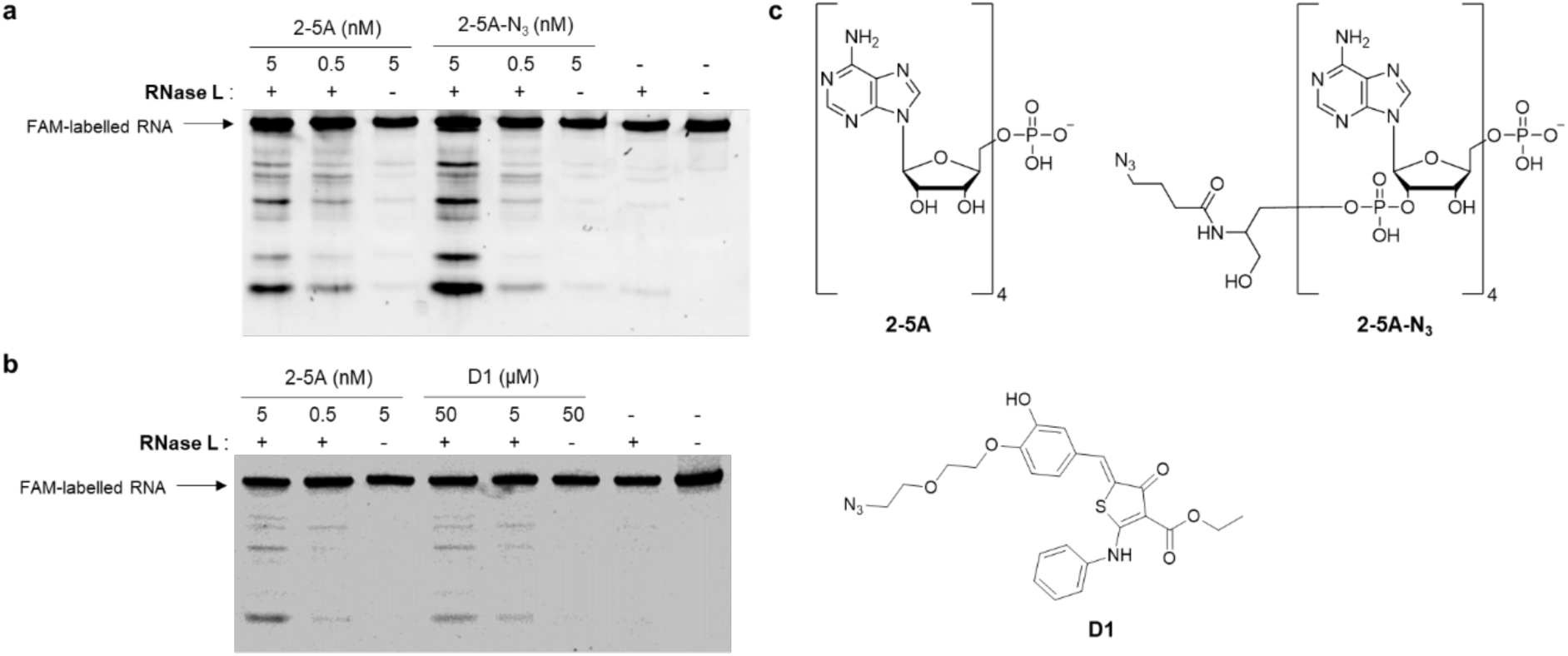
In vitro RNase L degradation assay with a 5’ 6-fluorescein-tagged model RNA containing multiple RNase L cleavage sites. **a**, Comparison of RNA degradation activity of RNase L in the presence of **2-5A** and **2-5A-N_3_** with no significant differential activity observed. **b**, Comparison of synthetic RNase L recruiter (D1) and 2-5A. The activity of **D1** in RNase L activation is ∼ 10,000 times weaker than **2-5A**. **c**, Chemical structures of **2-5A**, **2-5A-N_3_**, and **D1**.

**Extended Data Fig. 7.**
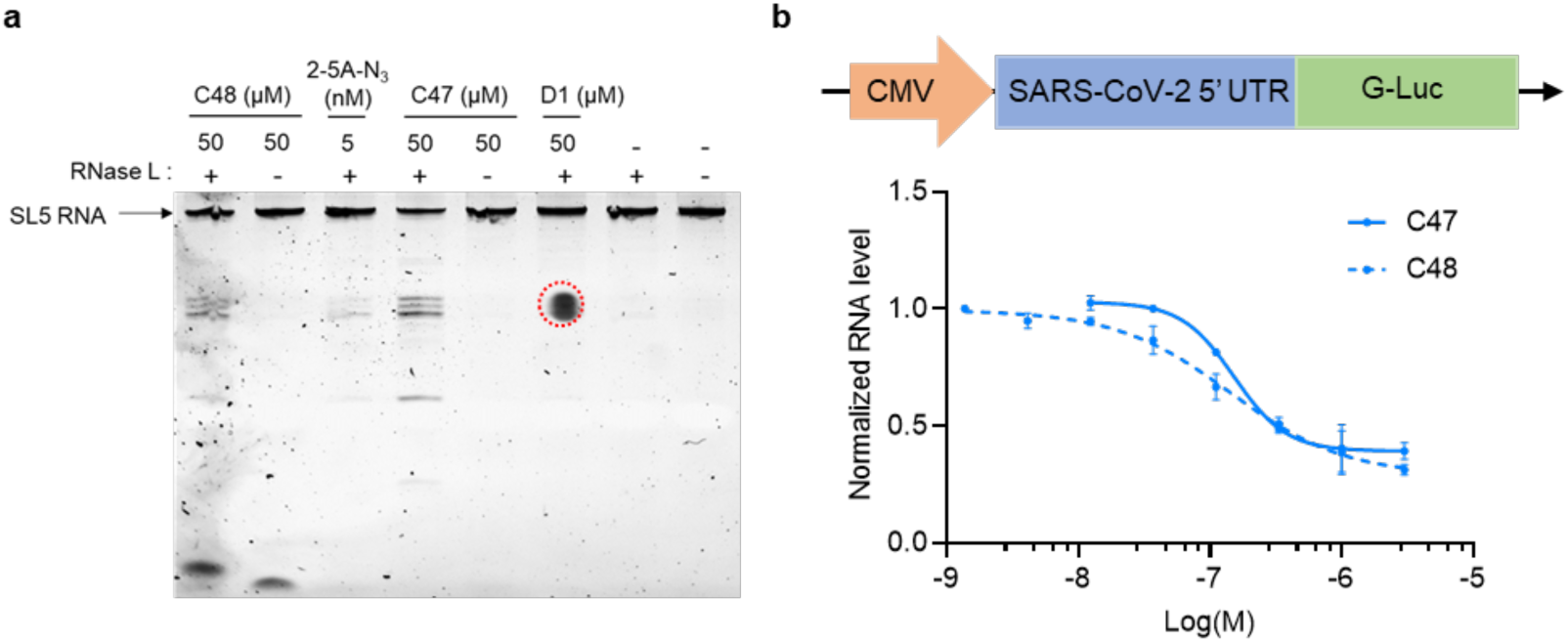
RNA degrading activity of C47 and C48. **a**, Comparison of two RLR moieties in the RIBOTAC modality using the in vitro RNase L degradation assay with purified SL5 RNA (red circle is a staining artifact). **b**, Cellular activity of **C47** and **C48** in SARS-CoV-2 5’ UTR expressing cells.

**Extended Data Fig. 8.**
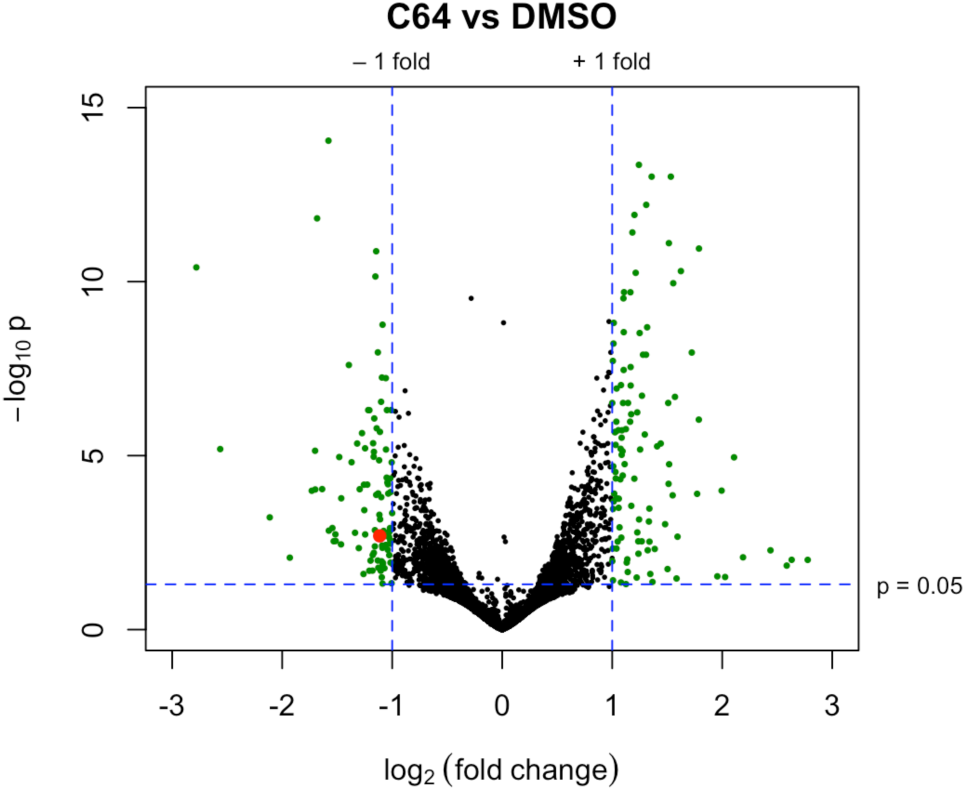
Volcano plot of differential gene expression in SARS-CoV-2 5’ UTR expressing cells treated with C64 (3 µM). DMSO-treated cells were used as a control. Red spot = SARS-CoV-2 5’ UTR transcript. The RNA-seq analysis was performed with three biological replicates (N = 3).

