## Supplementary Information for "Chemical-guided SHAPE sequencing (cgSHAPE-seq) informs the binding site of RNA-degrading chimeras targeting SARS-CoV-2 5’ untranslated region"

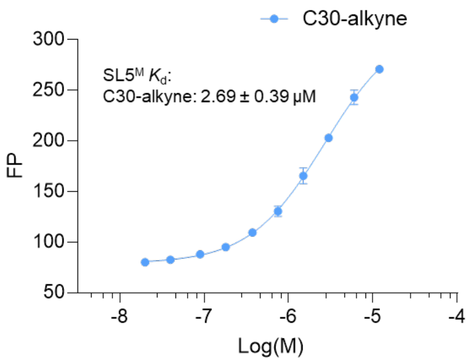


**Fig. S1 | Binding affinity of C30-alkyne to SL5$^M$ RNA.** C30-alkyne was used at a concentration of 80 nM. Each data point represents the mean fluorescence polarization value of two independent replicates (N = 2).

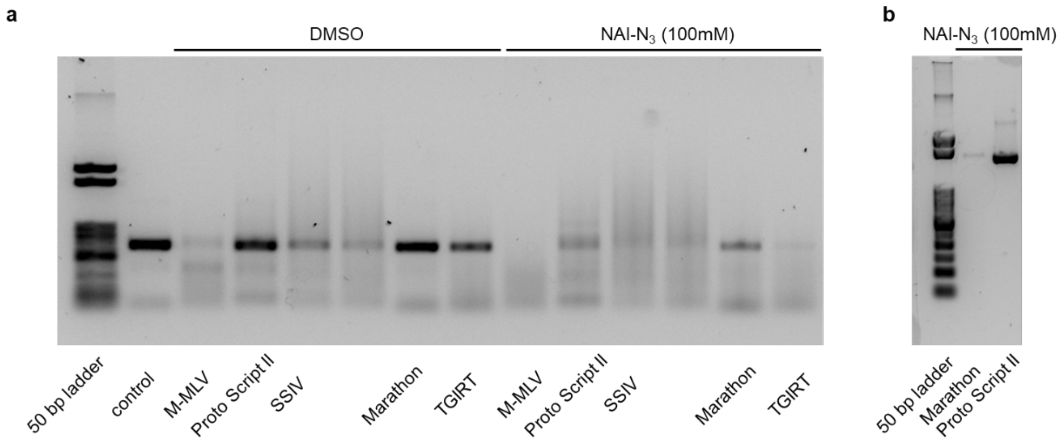


**Fig. S2 | Comparison of reverse transcriptases.** RNA was diluted in denaturing buffer (90% foramide, 5 mM EDTA), heated to 80 ℃ for 2 minutes and snap-cooled on ice. DMSO or NAI-N$_3$ was added and mixed well. The solution was heated at 80 ℃ for 5 minutes and cooled on ice. RNA was recovered using Qiagen RNeasy kit. RT reaction buffer: 50 mM Tris-HCl (pH 7.4), 75 mM KCl, 10 mM DTT, 3 mM $MnCl_2$. Manual suggested optimal reaction temperature of each enzyme was used (42 ℃ incubation 1 h for M-MLV, ProtoScript II, and Marathon; 50 ℃ incubation 1 h for SSIV and Maxima H minus). For TGIRT, reaction buffer: 50 mM Tris-HCl (pH 8.3), 75 mM KCl, 10 mM DTT, 1.5 mM $MgCl_2$. 50 ℃ incubation 1 h.

Reference sequence for **a** (378 nt): 5'-
aggtttataccttcccaggtaacAAACCAACCAACTTTCGATCTCTTGTAGATCTGTTCTCTAAACGAACTTTAAAATCTGTGT
GGCTGTCACTCGGCTGCGTGCTTAGTGCACTCACGCAGTATAATTAATAACTAATTACTGTCGTTGACAGGACAC
GAGTAACTCGTCTATCTTCTGCAGGCTGCTTACGGTTTCGTCCGTGTTGCAGCCGATCATCAGCACATCTAGGTT
TCGTCCGGGTGTGACCGAAAGGTAAGATGGAGAGCCTTGTCCCTGGTTTCAACGAGGGAGTCAAAGTTCTGTTT
GCCCTGATCTGCATCGCTGTGGCCGAGGCCAAGCCCACCGAGAACAACGAagacttcaacatcgtggccg.

Reference sequence for **b** (827 nt): 5'-

aacttcctttattttccttacagGGTTTTAGACAAAATCAAAAAGAAGGAAGGTGCTCACATTCCTTAAATTAAGGAGTAAGTCT
GCCAGCATTATGAAAGTGAATCTTACTTTTGTAAAACTTTATGGTTTGTGGAAAACAAATGTTTTTGAACATTTAAA
AAGTTCAGATGTTAGAAAGTTGAAAGGTTAATGTAAAACAATCAATATTAAAGAATTTTGATGCCAAAACTATTAGA
TAAAAGGTTAATCTACATCCCTACTAGAATTCTCATACTTAACTGGTTGGTTGTGTGGAAGAAACATACTTTCACA
ATAAAGAGCTTTAGGATATGATGCCATTTTATATCACTAGTAGGCAGACCAGCAGACTTTTTTTTATTGTGATATG
GGATAACCTAGGCATACTGCACTGTACACTCTGACATATGAAGTGCTCTAGTCAAGTTTAACTGGTGTCCACAGA
GGACATGGTTTAACTGGAATTCGTCAAGCCTCTGGTTCTAATTTCTCATTTGCAGGAAATGCTGGCATAGAGCAG
CACTAAATGACACCACTAAAGAAACGATCAGACAGATCTGGAATGTGAAGCGTTATAGACGATAACTGGCCTCAT
TTCTTCAAAATATCAAGTGTTGGGAAAGAAAAAAGGAAGTGGAATGGGTAACTCTTCTTGATTAAAGGTTATGTAA
TAACCAAATGCAATGTGAAATATTTTACTGGACTCTATTTTGAAAAACCATCTGTAAAAGACTGAGGTGGGGGTG
GGAGGCCAGCACGGTGGTGAGGCAGTTGAgaaaaagtaagtacttgacatgataagatac

(lowercase = primer binding sequences)

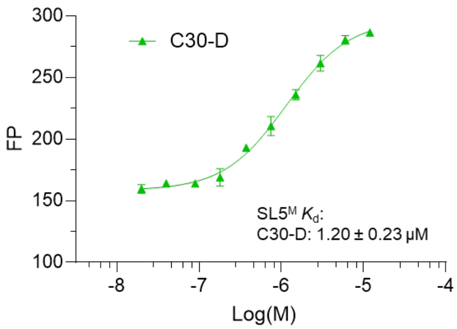


**Fig. S3 |** Binding affinity of C30-D with SL5$^{M}$ RNA.

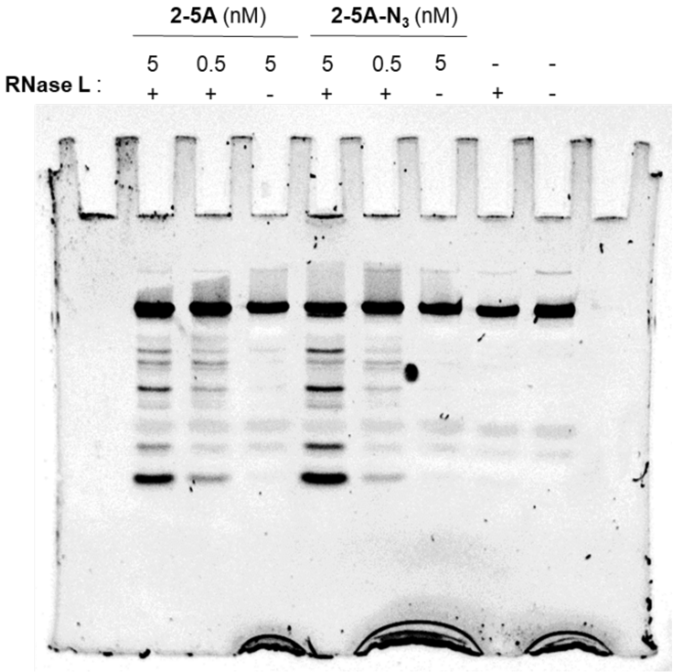


**Fig. S4 |** Full image for Extended Data Fig. 6a.

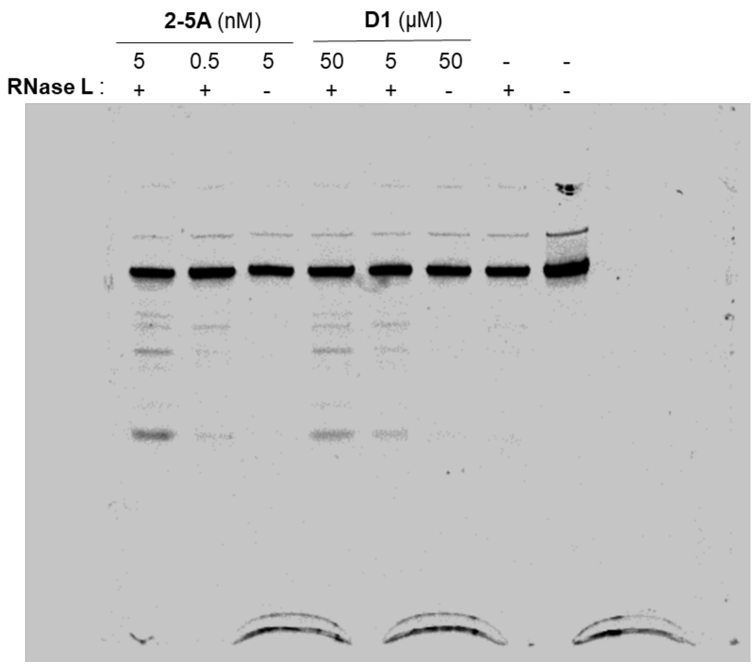


**Fig. S5 |** Full image for Extended Data Fig. 6b.

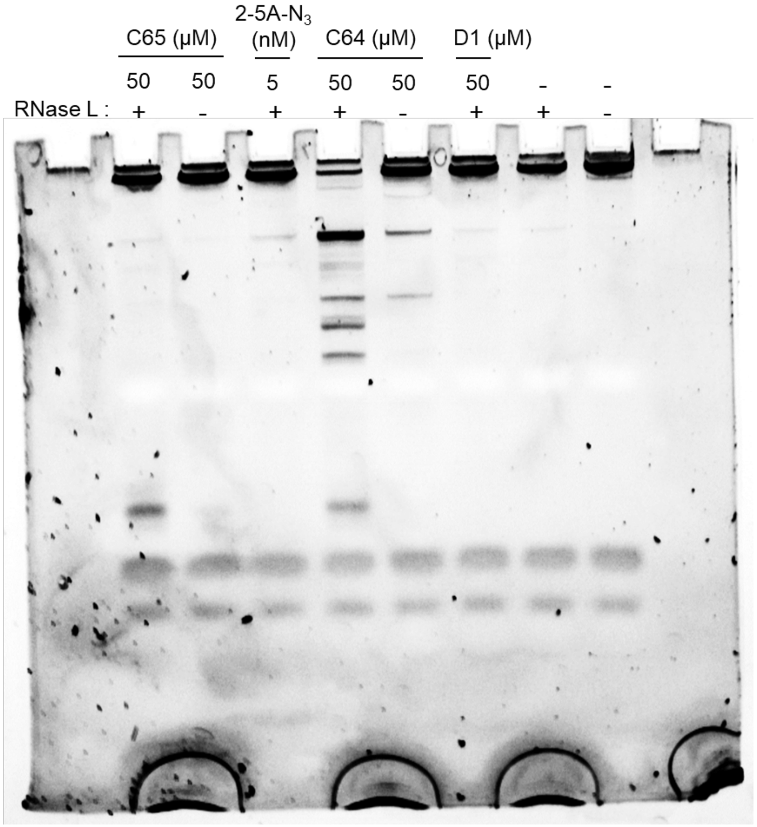


**Fig. S6 |** Full image for Fig. 4b.

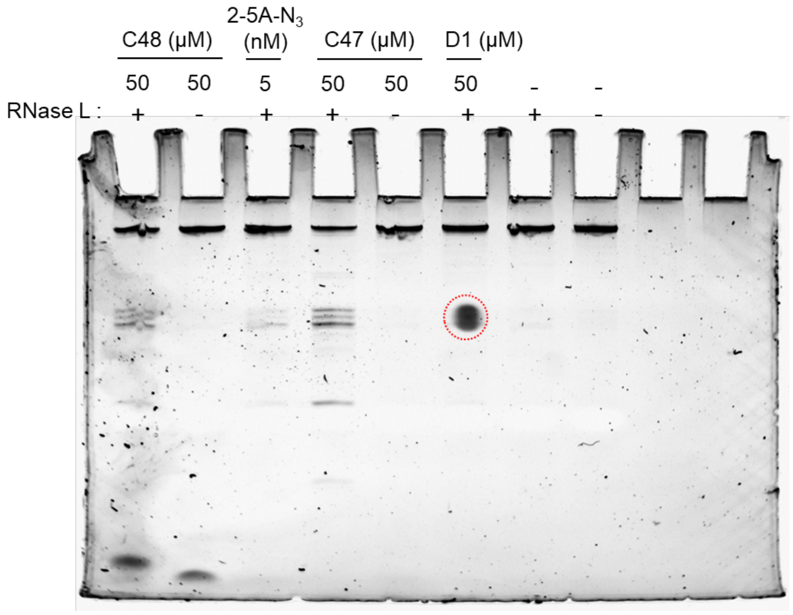


**Fig. S7 |** Full image for Extended Data Fig. 7a. (red circle is a staining artifact).

#### Methods

##### In vitro RNA transcription

In vitro RNA transcription was performed using a T7 RNA Polymerase (New England Biolabs, M0251L) following the manual's protocol. Usually, 100 µL transcription mixture containing 2 µg DNA template, 10 µL 10× transcription buffer, 0.5 mM NTPs, 2 µL RNase inhibitor (ApexBio, Cat. K1046) and 10 µL T7 polymerase was incubated at 37 °C for 12 h. 2 µL DNase I (10 U/µL, Roche) was added into the reaction mixture and incubated for 15 min at 37 °C. RNA was purified by precipitation with isopropanol or gel electrophoresis on a denaturing polyacrylamide gel. Finally, RNA was dissolved in molecular biology water and quantified by NanoDrop (ND-1000, Thermo, Waltham, MA, USA).

RNA sequences from in vitro transcription:

| | sequences |
|---|---|
| SL5 | UCGUUGACAGGACACGAGUAACUCGUCUAUCUUCUGCAGGCUGCUUACGGUUUCGUCCGUGUUG<br>CAGCCGAUCAUCAGCACAUCUAGGUUUCGUCCGGGUGUGACCGAAAGGUAAGAUGGAGAGCCUU<br>GUCCCUGGUUUCAACGA |
| $SL5^M$ | ACUCGUCUAUCUUCUGCAGGCUGCUUGCAGCCGAUCAUCAGCACAUCUACGGGUGUGACCGAAA<br>GGUAAGAUGGAGAGC |
| 1 | ACUCAUCUAUCUUCUGCAGGCUGCUUGCAGCCGAUCAUCAGCACAUCUACGGGUGUGACCGAAA<br>GGUAAGAUGGAGAGC |
| 2 | ACUCCUCUAUCUUCUGCAGGCUGCUUGCAGCCGAUCAUCAGCACAUCUACGGGUGUGACCGAAA<br>GGUAAGAUGGAGAGC |
| 3 | ACUCUUCUAUCUUCUGCAGGCUGCUUGCAGCCGAUCAUCAGCACAUCUACGGGUGUGACCGAAA<br>GGUAAGAUGGAGAGC |
| 4 | ACUCGGUCUAUCUUCUGCAGGCUGCUUGCAGCCGAUCAUCAGCACAUCUACGGGUGUGACCGAA<br>AGGUAAGAUGGAGAGC |
| 5 | ACUCGUCUAUCUUCUGCAGGCUGCUUGCAGCCGAUCAUCAGCACAUCUACGGGUGUGACCGAAA<br>GGUAAGAUGGAGGAGC |
| 6 | ACUCGUCUAUCUUCUGCAGGCUGCUUGCAGCCGAUCAUCAGCACAUCUACGGGUGUGACCGAAA<br>GGUAAGAUGGAAGAGC |
| 7 | ACUCGUCUAUCUUCUGCAGGCUGCUUGCAGCCGAUCAUCAGCACAUCUACGGGUGUGACCGAAA<br>GGUAAGAUGGACGAGC |
| 8 | ACUUGUCUAUCUUCUGCAGGCUGCUUGCAGCCGAUCAUCAGCACAUCUACGGGUGUGACCGAAA<br>GGUAAGAUGGAAAGC |
| 9 | ACUCGCCUAUCUUCUGCAGGCUGCUUGCAGCCGAUCAUCAGCACAUCUACGGGUGUGACCGAAA<br>GGUAAGAUGGGGAGC |
| 10 | ACUCGUCUAUCUUCUGCAGGCUGCUUGCAGCCGAUCAUCAGCACAUCUACGGGUGUGACCGAAA<br>GGUAAGAUAAAGAGC |
| 11 | ACUCGUCUAUCUUCUGCAGGCUGAAACACAGCCGAUCAUCAGCACAUCUACGGGUGUGACCGAAA<br>GGUAAGAUGGAGAGC |
| 12 | ACUCGUCUAUCUUCUGCAGGCUGCUUGCAGCCGAUCAUCAGCACAUGAAACACGUGUGACCGAA<br>AGGUAAGAUGGAGAGC |
| 13 | ACUCGUCUAUCUUCUGCAGGCUGCUUGCAGCCGAUCAUCAGCACAUCUACGGGUGUGACGAAAC<br>ACGUAAGAUGGAGAGC |

(Mutations in $SL5^M$ are coloured in purple)

##### Fluorescence Polarization Binding Assay

Synthetic RNA oligomers were purchased from GenScript or Integrated DNA Technologies (IDT) and reconstituted in nuclease-free water (Invitrogen #AM9932). Compounds (1 mM in DMSO) were diluted in 2× assay buffer (40 mM MES, 200 mM NaCl, 0.0125% TritonX, pH 6.5) to 0.16 µM.

A 1:2 dilution series (8 points) of each RNA was prepared in 20–30 µL water to desired concentrations (i.e., 0.09 – 12 µM). 20–30 µL 2× working solution containing the 2× assay buffer and the small-molecule ligand was added to each RNA sample in 1:1 (v/v) and mixed by pipetting.

For fluorescence polarization measurement, 20 µL of the above 1× working solution was transferred into a 384-well, black, flat-bottom microplates (Greiner Bio-One) in duplicates or triplicates. The plate was equilibrated at room temperature for 5 min before being read using a microplate reader (SYNERGY H1, BioTek; Excitation/Emission = 360/460 nm) at 25 °C. The experimental data were analysed using Prism 8.0 software (Graphpad Software, San Diego, CA, USA). The dissociation constant ($K_d$) was calculated with 95 % confidence interval after nonlinear curve fitting (Sigmoidal, 4 parameters).

RNA sequences from IDT (first two) or GenScript (last three):

| | sequences |
|---|---|
| SL5A | GGCUGCUUACGGUUUCGUCCGUGUUGCAGCC |
| SL5B | CACAUCUAGGUUUCGUCCGGGUGUG |
| 14 | ACUCUCUAUCUUCUGCAGGCUGCUUGCAGCCGAUCAUCAGCACAUCUACGGGUGUGACCGAAAG GUAAGAUGGAGAGC |
| 15 | ACUCGUCUAUCUUCUGCAGGCUGCUUGCAGCCGAUCAUCACCACAUCUACGGGUGUCUCCGAAA GGUUUGAUGGAGAGC |
| 16 | ACUCGUCUAUCUUCUGCAGGCUGCUUGCAGCCCUUCUUCUCCACAUCUACGGGUGUGACCGAAA GGUAAGAUGGAGAGC |

(Mutations in $SL5^M$ are coloured in purple)

#### Reactivity of FAI-$N_3$ with four nucleotides in a denatured RNA

400 ng model RNA (in vitro transcribed, 2.5 µL in water) was added 16.5 µL denaturing buffer (90% formamide, 5 mM EDTA) and the mixture was heated at 80 °C for 5 min. 1 µL FAI-$N_3$ (2 M in DMSO) or DMSO (blank control) was added to the hot RNA solution and mixed by pipetting. The solution was incubated at 80 °C for 5 min and quenched by adding Qiagen buffer RLT. The RNA was then extracted using a Qiagen RNeasy kit following the user's manual. 25 ng RNA was used for RT-PCR following the same protocol for cgSHAPE-seq. Mutational profiling was performed using ShapeMapper2 and the result was used to generate Extended Data Fig. 2[1].

Reference sequence for ShapeMapper2 is:
5'aacttcctttattttccttacagGGTTTTAGACAAAATCAAAAAGAAGGAAGGTGCTCACATTCCTTAAA
TTAAGGAGTAAGTCTGCCAGCATTATGAAAGTGAATCTTACTTTTGTAAAACTTTATGGTTTG
TGGAAAACAAATGTTTTTGAACATTTAAAAAGTTCAGATGTTAGAAAGTTGAAAGGTTAATGT
AAAACAATCAATATTAAAGAATTTTGATGCCAAAACTATTAGATAAAAGGTTAATCTACATCC
CTACTAGAATTCTCATACTTAACTGGTTGGTTGTGTGGAAGAaacatactttcacaataaagagc
lowercase = forward and reverse primers

#### Surface Plasmon Resonance (SPR) Assay

SPR experiments were performed on a Biacore T200 (GE Healthcare) instrument at 25 °C using streptavidin pre-coated SA sensor chips (GE Healthcare). The running buffer composed of 10 mM HEPES, 100 mM NaCl, 0.05% Tween 20 (w/v), 5 mM EDTA, 0.1% (v/v) DMSO at pH 6.8 was prepared freshly, filtered through the 0.22 µm PVDF membrane prior to use. 3'-biotinylated RNA SL5 was prepared by 3' ligation according to the literature protocol[2]. For immobilization of the biotinylated RNA, the sensor chip was firstly conditioned with 3 consecutive 1 min injections of high salt solution (50 mM NaOH, 1M NaCl) at a flow rate of 10 µL/min. Next, the biotinylated RNA was diluted 100× in running buffer (100 nM) and applied over the streptavidin sensor chip surface at a flow rate of 10 µL/min to achieve immobilization level of about 500 RU. Finally, alkyne-PEG-biotin (50 µM in running buffer) was injected (1 min, 10 µL/min) to block remaining streptavidin surface binding sites. The kinetics analysis was performed following the BiaControl Software Wizard Kinetics protocol. Compound **C30** (HCl salt form, 10 mM in water) was diluted in the running buffer to the following concentrations (0, 0.075, 0.15, 0.3, 0.6 µM) and titrated over the immobilized RNA SL5 (contact time: 1 min, flow rate: 30 µL/min). The data analysis and plotting were performed using the instrument BiaEvaluation Software. All monitored resonance signals were subtracted with signals from a non-binding reference channel. Kinetic values ($K_d$, $k_a$, $k_d$) were calculated using the BiaEvaluation Software Binding Affinity protocol with 1:1 fitting. 3'-biotinylated RNA SL5 sequence: 5'-UCGUUGACAGGACACGAGUAACUCGUCUAUCUUCUGC AGGCUGCUUACGGUUUCGUCCGUGUUGCAGCCGAUCAUCAGCACAUCUAGGUUUCGUC CGGGUGUGACCGAAAGGUAAGAUGGAGAGCCUUGUCCCUGGUUUCAACGA-biotin

**Mutational profiling for compounds C30-D, C30-BD and C30-NM**

Protocols were similar to cgSHAPE-seq except RNA modification step:

For **C30-D** and **C30-BD**, total RNA was added water and 5× folding/reaction buffer (500 mM HEPES pH 7.4, 500 mM KCl, 30 mM $MgCl_2$) to make a 47.5 µL solution. The solution was incubated at 37 °C for 30 min to refold. 2.5 µL **C30-D or C30-BD** (2 mM in DMSO) or DMSO (blank control) was added to the total RNA and mixed well by pipetting. The mixture was then irradiated under 365 nM UV light in a photocrosslinker (Fisher Cat. #13-245-221) for 15 min. The RNA was then recovered using a RNeasy kit (Qiagen).

For **C30-NM**, total RNA was added water and 5× folding/reaction buffer (500 mM HEPES pH 7.4, 500 mM KCl, 30 mM $MgCl_2$) to make a 47.5 µL solution. The solution was incubated at 37 °C for 30 min to refold. 2.5 µL **C30-NM** (2 mM in DMSO) or DMSO (blank control) was added to the total RNA and mixed well by pipetting. The mixture was then incubated at 37 °C for 12 h. The RNA was recovered using a RNeasy kit (Qiagen).

500 ng total RNA was used for reverse transcription and then PCR as described in cgSHAPE-seq protocol. All reactions were performed in triplicates.

**Cytotoxicity Assay**

The ACE-2 expressing A549 cells were seeded in a 384-well plate at 10,000 cells per well in 30 µl of growth media[3]. The cells were then treated with various concentrations of **C64** (2.44 nM–20 µM at 1:2 serial dilution) and a DMSO control (0.1%) in triplicates. After incubation for 48 h in the presence of **C64**, the cell viability was measured using CellTitre-Glo (Promega, G9242) and a luminescence plate reader (BioTek, Cytation 5). The relative viability was normalized to the DMSO control.

**RNA-seq**

The Stranded mRNA-Seq was performed using the Illumina NovaSeq 6000 Sequencing System at the University of Kansas Medical Center Genomics Core (Kansas City, KS). Quality control on RNA submissions was completed using the Agilent Bioanalyzer 2100 using the RNA 6000 Nano Assay kit (Agilent Technologies, 5067-1511). Total RNA (750 ng) was used to initiate the library preparation protocol. The total RNA fraction was processed by oligo dT bead capture of mRNA, fragmentation, reverse transcription into cDNA, end repair of cDNA, ligation with the appropriate Unique Dual Index (UDI) adaptors, strand selection and library amplification by PCR using the Universal Plus mRNA-seq with UDI preparation kit (NuGEN Technologies, 0508-02).

Library validation was performed using the DNA 1000 Assay kit (Agilent Technologies, 5067-1504) on the Agilent Bioanalyzer 2100. Concentration of each library was determined by qPCR using the with the Roche Lightcycler96 using FastStart Essential DNA Green Master (Roche 06402712001) and KAPA Library Quant (Illumina) DNA Standards 1-6 (KAPA Biosystems KK4903). Libraries were pooled based on equal molar amounts to 1.9 nM for multiplexed sequencing.

Pooled libraries were denatured with 0.2N NaOH (0.04N final concentration) and neutralized with 400 mM Tris-HCl pH 8.0. A dilution of the pooled libraries to 380 pM was performed in the sample tube, on instrument, followed by onboard clonal clustering of the patterned flow cell using the NovaSeq 6000 S1 Reagent Kit v1.5 (200 cycle) (Illumina 20028318). A 2x101 cycle sequencing profile with dual index reads was completed using the following sequence profile: Read 1 – 101 cycles x Index Read 1 – 8 cycles x Index Read 2 – 8 cycles x Read 2 – 101 cycles. Following collection, sequence data was converted from .bcl file format to. fastq file format using bcl2fastq software and de-multiplexed into individual sequences for data distribution. We followed the differential gene expression (DGE) analysis pipeline developed by Maude and Cebola (https://github.com/CebolaLab/RNA-seq). The result was used to generate Extended Data Fig. 7 and Table S1.

**Table S1. Significantly changed RNA transcripts in C64-treated SARS-CoV2-5' UTR reporter cells determined by RNA-seq analysis (| $log_2$FoldChange | > 2 and adjusted p value (padj) < 0.05).**

| Gene | HGNC_symbol | baseMean | $log_2$FoldChange | lfcSE | pvalue | padj |
|---|---|---|---|---|---|---|
| ENSG00000117600 | PLPPR4 | 22.3 | -3.0 | 0.84 | 1.58E-05 | 7.60E-04 |
| ENSG00000111344 | RASAL1 | 71.8 | -2.8 | 0.39 | 3.54E-14 | 3.89E-11 |
| ENSG00000260448 | LCMT1-AS1 | 35.1 | -2.6 | 0.54 | 3.90E-08 | 6.07E-06 |
| ENSG00000105392 | CRX | 15.5 | -2.6 | 0.86 | 7.58E-05 | 2.51E-03 |
| ENSG00000157601 | MX1 | 14.2 | -2.6 | 1.05 | 4.42E-04 | 9.29E-03 |
| ENSG00000259030 | FPGT-TNNI3K | 62.9 | -2.6 | 0.53 | 4.37E-08 | 6.49E-06 |
| ENSG00000104320 | NBN | 27.3 | -2.4 | 0.68 | 8.41E-06 | 4.64E-04 |
| ENSG00000204116 | CHIC1 | 43.2 | -2.4 | 0.84 | 1.65E-04 | 4.49E-03 |
| ENSG00000157087 | ATP2B2 | 14.6 | -2.3 | 1.01 | 6.96E-04 | 1.27E-02 |
| ENSG00000050767 | COL23A1 | 81.1 | -2.1 | 0.59 | 1.17E-05 | 5.96E-04 |
| ENSG00000220008 | LINGO3 | 26.3 | -2.1 | 0.65 | 5.38E-05 | 1.93E-03 |
| ENSG00000243708 | PLA2G4B | 31.8 | -2.1 | 0.62 | 3.17E-05 | 1.29E-03 |
| ENSG00000287908 | | 25.0 | -2.0 | 0.91 | 6.33E-04 | 1.19E-02 |
| ENSG00000156983 | BRPF1 | 357.2 | 2.0 | 1.49 | 2.55E-03 | 3.08E-02 |
| ENSG00000271781 | | 24.6 | 2.0 | 0.61 | 3.17E-05 | 1.29E-03 |
| ENSG00000275215 | RNA5-8SN3 | 55.9 | 2.1 | 0.44 | 8.66E-08 | 1.12E-05 |
| ENSG00000259660 | DNM1P47 | 52.3 | 2.2 | 0.93 | 3.81E-04 | 8.37E-03 |
| ENSG00000151702 | FLI1 | 1074.6 | 2.4 | 0.90 | 2.03E-04 | 5.27E-03 |
| ENSG00000277209 | RPPH1 | 32.2 | 2.5 | 0.62 | 2.47E-06 | 1.75E-04 |
| ENSG00000085998 | POMGNT1 | 373.0 | 2.6 | 1.26 | 8.47E-04 | 1.45E-02 |
| ENSG00000204970 | PCDHA1 | 17.3 | 2.6 | 1.69 | 1.48E-03 | 2.10E-02 |

| | | | | | | |
|---|---|---|---|---|---|---|
| ENSG00000177646 | ACAD9 | 363.3 | 2.6 | 1.13 | 4.85E-04 | 9.87E-03 |
| ENSG00000254873 | | 29.8 | 2.7 | 1.21 | 5.99E-04 | 1.15E-02 |
| ENSG00000136271 | DDX56 | 1346.0 | 2.8 | 1.20 | 4.88E-04 | 9.92E-03 |
| ENSG00000170160 | CCDC144A | 15.5 | 2.9 | 1.76 | 1.61E-03 | 2.23E-02 |
| ENSG00000288534 | | 193.6 | 3.0 | 1.25 | 3.55E-04 | 8.00E-03 |
| ENSG00000115155 | OTOF | 24.3 | 3.5 | 0.62 | 7.42E-10 | 2.27E-07 |
| ENSG00000251357 | | 109.3 | 3.6 | 2.18 | 1.17E-03 | 1.83E-02 |
| ENSG00000227082 | LINC02798 | 18.7 | 3.6 | 1.61 | 4.10E-04 | 8.83E-03 |
| ENSG00000107669 | ATE1 | 395.9 | 3.6 | 2.22 | 1.13E-03 | 1.79E-02 |
| ENSG00000082438 | COBLL1 | 434.3 | 3.8 | 1.13 | 2.07E-05 | 9.35E-04 |
| ENSG00000070831 | CDC42 | 1913.2 | 4.3 | 1.60 | 1.65E-04 | 4.49E-03 |
| ENSG00000285976 | | 1863.1 | 4.7 | 1.50 | 3.99E-05 | 1.54E-03 |
| ENSG00000174405 | LIG4 | 133.6 | 5.3 | 1.73 | 4.44E-05 | 1.65E-03 |
| ENSG00000126561 | STAT5A | 128.2 | 6.1 | 2.11 | 5.94E-05 | 2.07E-03 |
| ENSG00000196305 | IARS1 | 1519.8 | 6.2 | 1.62 | 3.08E-06 | 2.07E-04 |

#### Chemistry

**General Methods.** Reagents and solvents were purchased from commercial sources (Fisher, Sigma-Aldrich and Combi-Blocks) and used as received. Reactions were tracked by TLC (Silica gel 60 $F_{254}$, Merck) and Waters ACQUITY UPLC-MS system (ACQUITY UPLC H Class Plus in tandem with Qda Mass Detector). Intermediates and products were purified by a Teledyne ISCO Combi-Flash system using prepacked $SiO_2$ cartridges. NMR spectra were acquired on a Bruker AV400 instrument (400 MHz for $^1$H NMR, 100 MHz for $^{13}$C NMR) or Bruker AV500 instrument (500 MHz for $^1$H NMR, 126 MHz for $^{13}$C NMR). $^{13}$C shifts were obtained with $^1$H decoupling. MestReNova 14.0.1 developed by MESTRELAB RESEARCH was used for NMR data processing. MS-ESI spectra were recorded on Waters Qda Mass Detector. HPLC was performed on Waters ACQUITY UPLC H Class Plus system using Waters BEH C18 (2.1 mm × 50 mm, 1.7 μm) column and peak detection at 254nm with UV.

Compounds **C2**, **C4**, **C6** and **C29** were synthesized following procedures reported in literature and verified by NMR and Mass spectra[2,4].

General procedures for compounds **C30**, **C31**, **C32**, **C36 and C34.**

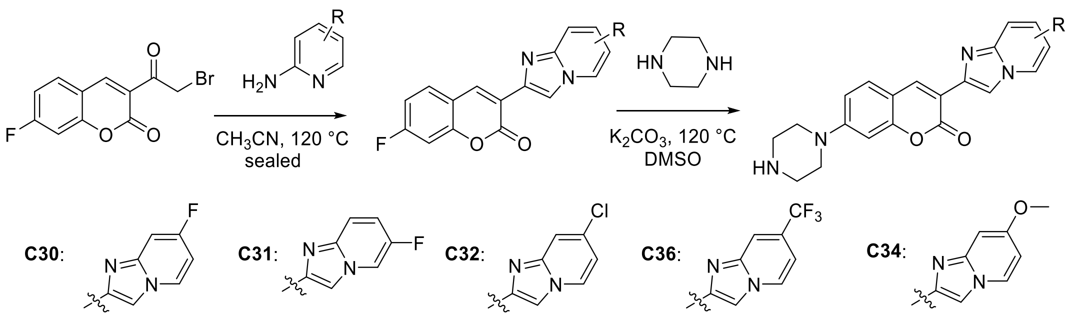

Step 1: Synthesis of Coumarin intermediate

A mixture of 3-(2-bromoacetyl)-7-fluoro-2*H*-chromen-2-one (1 eq) and substituted aminopyridine (1.2 eq) in acetonitrile was heated to 120 °C for 20 min in a Biotage Initiator+ microwave reactor. TLC and LC-MS showed completion of reaction. The reaction mixture was filtered, and the solid was washed with acetonitrile to afford a yellow solid which was used in next step without further purification.

Step 2: Synthesis of final product

To a solution of Coumarin intermediate (1 eq) and piperazine (1.5 eq) in DMSO was added $K_2CO_3$ (3 eq). The reaction mixture was heated to 120 °C and stirred for 2 h. TLC and LC-MS showed completion of reaction. The mixture was cooled to room temperature, poured into ice water and extracted with ethyl acetate. Organic layer was washed with brine, dried with anhydrous sodium sulphate and removed under vacuum. The residue was purified by silica gel column chromatography (0 – 10% $CH_3OH$ in $CH_2Cl_2$) to afford product as yellow solid.

**3-(7-fluoroimidazo[1,2-*a*]pyridin-2-yl)-7-(piperazin-1-yl)-2*H*-chromen-2-one (C30)**

Following General procedures, from 3-(2-bromoacetyl)-7-fluoro-2*H*-chromen-2-one (50 mg, 0.17 mmol) and 4-fluoro-2-aminopyridine (22 mg, 0.2 mmol), 21 mg of compound **C30** was obtained as a yellow solid (two steps yield, 33%). MS-ESI (*m*/*z*) $[M+H]^+$ 365.12.

$^1$H NMR (400 MHz, DMSO-$d_6$) δ 8.73 – 8.69 (m, 2H), 8.53 (s, 1H), 7.73 (d, *J* = 8.9 Hz, 1H), 7.39 (dd, *J* = 10.1, 2.6 Hz, 1H), 7.05 (dd, *J* = 8.9, 2.4 Hz, 1H), 6.99 (td, *J* = 7.6, 2.6 Hz, 1H), 6.93 (d, *J* = 2.4 Hz, 1H), 3.48 (t, *J* = 5.2 Hz, 4H), 3.05 (t, *J* = 5.2 Hz, 4H).

$^{13}$C NMR (126 MHz, DMSO-$d_6$) δ 160.2 (d, *J* = 248.0 Hz), 159.4, 154.8, 153.1, 144.3 (d, *J* = 12.5 Hz), 139.3, 138.5, 129.7, 129.3 (d, *J* = 11.8 Hz), 114.9, 112.1 (d, *J* = 28.1 Hz), 110.6, 104.2 (d, *J* = 29.8 Hz), 99.8, 99.6, 99.5, 46.0, 43.8.

**3-(6-fluoroimidazo[1,2-*a*]pyridin-2-yl)-7-(piperazin-1-yl)-2*H*-chromen-2-one (C31)**

Following General Procedures, from 3-(2-bromoacetyl)-7-fluoro-2*H*-chromen-2-one (50 mg, 0.17 mmol) and 5-fluoro-2-aminopyridine (22 mg, 0.2 mmol), 23 mg of compound **C31** was obtained as a yellow solid (two steps yield, 36%). MS-ESI (*m*/*z*) $[M+1]^+$ 365.18.

$^1$H NMR (400 MHz, DMSO-$d_6$) δ 8.84 (dd, *J* = 4.6, 2.4 Hz, 1H), 8.71 (s, 1H), 8.54 (s, 1H), 7.70 (d, *J* = 8.8 Hz, 1H), 7.61 (dd, *J* = 9.9, 5.2 Hz, 1H), 7.40 – 7.35 (m, 1H), 7.02 (dd, *J* = 8.9, 2.4 Hz, 1H), 6.87 (d, *J* = 2.3 Hz, 1H), 3.34 (t, *J* = 5.2 Hz, 4H), 2.88 (t, *J* = 5.2 Hz, 4H).

$^{13}$C NMR (126 MHz, DMSO-$d_6$) δ 159.6, 155.0, 153.7, 152.3 (d, *J* = 232.0 Hz), 142.1, 139.7, 138.7, 129.6, 117.3 (d, *J* = 26.2 Hz), 116.8 (d, *J* = 9.9 Hz), 114.4, 114.0, 113.6, 111.7, 110.1, 99.3, 47.5, 45.0.

**3-(7-chloroimidazo[1,2-*a*]pyridin-2-yl)-7-(piperazin-1-yl)-2*H*-chromen-2-one** (**C32**)

Following General Procedures, from 3-(2-bromoacetyl)-7-fluoro-2*H*-chromen-2-one (50 mg, 0.17 mmol) and 4-chloro-2-aminopyridine (26 mg, 0.2 mmol), 31 mg of compound **C32** was obtained as a yellow solid (two steps yield, 46%). MS-ESI (*m*/*z*) $[M+1]^+$ 381.13.

$^1$H NMR (400 MHz, DMSO-$d_6$) δ 8.70 (s, 1H), 8.67 (d, *J* = 7.3 Hz, 1H), 8.56 (s, 1H), 7.71 (d, *J* = 8.0 Hz, 1H), 7.70 (s, 1H), 7.04 – 6.98 (m, 2H), 6.88 (d, *J* = 2.3 Hz, 1H), 3.35 (t, *J* = 5.2 Hz, 4H), 2.89 (t, *J* = 5.1 Hz, 4H).

$^{13}$C NMR (126 MHz, DMSO-$d_6$) δ 159.5, 155.0, 153.7, 143.9, 139.4, 138.9, 130.6, 129.7, 128.2, 114.7, 114.2, 113.1, 112.6, 111.7, 110.2, 99.3, 47.4, 44.9.

**7-(piperazin-1-yl)-3-(7-(trifluoromethyl)imidazo[1,2-*a*]pyridin-2-yl)-2*H*-chromen-2-one** (**C36**)

Following General Procedures, from 3-(2-bromoacetyl)-7-fluoro-2*H*-chromen-2-one (50 mg, 0.17 mmol) and 4-trifluoromethyl-2-aminopyridine (32 mg, 0.2 mmol), 35 mg of compound **C36** was obtained as a yellow solid (two steps yield, 48%).MS-ESI (*m*/*z*) $[M+1]^+$ 415.19.

$^1$H NMR (500 MHz, DMSO-$d_6$) δ 9.33 (s, 1H), 8.82 (d, *J* = 7.1 Hz, 1H), 8.73 (s, 1H), 8.70 (s, 1H), 7.96 (s, 1H), 7.71 (d, *J* = 8.8 Hz, 1H), 7.19 (dd, *J* = 7.1, 1.9 Hz, 1H), 7.02 (dd, *J* = 8.9, 2.5 Hz, 1H), 6.94 (d, *J* = 2.4 Hz, 1H), 3.60 (t, *J* = 5.3 Hz, 4H), 3.16 (brs, 4H).

$^{13}$C NMR (126 MHz, DMSO-$d_6$) δ 159.3, 155.0, 152.7, 141.8, 139.6, 139.5, 130.0, 129.0, 124.5, 122.4, 114.1, 113.6, 112.2, 110.8, 108.0, 100.3, 44.0, 42.2.

**3-(7-methoxyimidazo[1,2-*a*]pyridin-2-yl)-7-(piperazin-1-yl)-2*H*-chromen-2-one** (**C34**)

Following General Procedures, from 3-(2-bromoacetyl)-7-fluoro-2*H*-chromen-2-one (50 mg, 0.17 mmol) and 4-methoxypyridin-2-amine (25 mg, 0.2 mmol), 33 mg of compound **C34** was obtained as a yellow solid (two steps yield, 51%). MS-ESI (*m*/*z*) $[M+1]^+$ 377.18.

$^1$H NMR (400 MHz, DMSO-$d_6$) δ 8.61 (s, 1H), 8.45 (d, *J* = 7.4 Hz, 1H), 8.34 (s, 1H), 7.66 (d, *J* = 8.8 Hz, 1H), 7.00 (dd, *J* = 8.9, 2.4 Hz, 1H), 6.88 (d, *J* = 2.5 Hz, 1H), 6.85 (d, *J* = 2.4 Hz, 1H), 6.61 (dd, *J* = 7.4, 2.5 Hz, 1H), 3.85 (s, 3H), 3.30 (t, *J* = 5.1 Hz, 4H), 2.85 (t, *J* = 5.1 Hz, 4H).

$^{13}$C NMR (126 MHz, DMSO-$d_6$) δ 160.0, 158.4, 155.2, 154.0, 146.2, 138.5, 138.2, 129.8, 128.2, 115.4, 112.1, 111.6, 110.7, 107.1, 99.8, 94.1, 56.0, 48.2, 45.6.

Synthesis of **C30** conjugates (**C30-alkyne, C47 and C48**)

**7-(4-(3,6,9,12-tetraoxapentadec-14-yn-1-yl)piperazin-1-yl)-3-(7-fluoroimidazo[1,2-a]pyridin-2-yl)-2H-chromen-2-one** (**C30-alkyne**)

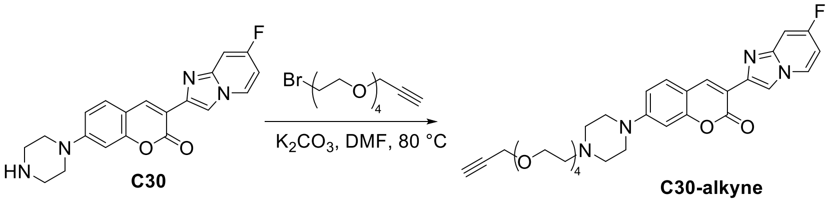


Compound **C30** (50 mg, 0.14 mmol) in DMF (1 mL) was added $K_2CO_3$ (37 mg, 0.27 mmol) and Propargyl-PEG$_4$-Br (80 mg, 0.27 mmol) and the reaction mixture was heated at 80 °C for overnight. The reaction mixture was added ice-water and filtered. The precipitate was washed with water and dried. The crude product was purified by silica gel column chromatography (0 – 5% $CH_3OH$ in $CH_2Cl_2$) to afford **C30-alkyne** as a yellow solid (47 mg, 72%). MS-ESI (*m/z*) $[M+1]^+$ 579.22.

$^1$H NMR (500 MHz, DMSO-$d_6$) δ 8.71 – 8.68 (m, 2H), 8.52 (s, 1H), 7.69 (d, *J* = 8.8 Hz, 1H), 7.38 (dd, *J* = 10.2, 2.6 Hz, 1H), 7.02 (dd, *J* = 8.9, 2.4 Hz, 1H), 6.97 (td, *J* = 7.6, 2.6 Hz, 1H), 6.87 (d, *J* = 2.3 Hz, 1H), 4.15 (d, *J* = 2.4 Hz, 2H), 3.58 – 3.51 (m, 14H), 3.42 (t, *J* = 2.5 Hz, 1H), 3.37 (t, *J* = 5.1 Hz, 4H), 2.58 – 2.53 (m, 6H).

$^{13}$C NMR (126 MHz, DMSO-$d_6$) δ 160.2 (d, *J* = 249.5 Hz), 159.5, 154.9, 153.4, 144.3 (d, *J* = 14.9 Hz), 139.4, 138.6, 129.6, 129.2 (d, *J* = 11.9 Hz), 114.5, 112.1, 111.7, 110.2, 104.2 (d, *J* = 29.5 Hz), 99.5 (d, *J* = 23.5 Hz), 99.4, 80.3, 77.1, 69.8, 69.7, 69.5, 68.5, 68.4, 57.5, 57.1, 52.8, 46.8.

**Ethyl (*Z*)-5-(4-(2-(2-(4-(13-(4-(3-(7-fluoroimidazo[1,2-a]pyridin-2-yl)-2-oxo-2H-chromen-7-yl)piperazin-1-yl)-2,5,8,11-tetraoxatridecyl)-1H-1,2,3-triazol-1-yl)ethoxy)ethoxy)-3-hydroxybenzylidene)-4-oxo-2-(phenylamino)-4,5-dihydrothiophene-3-carboxylate** (**C47**)

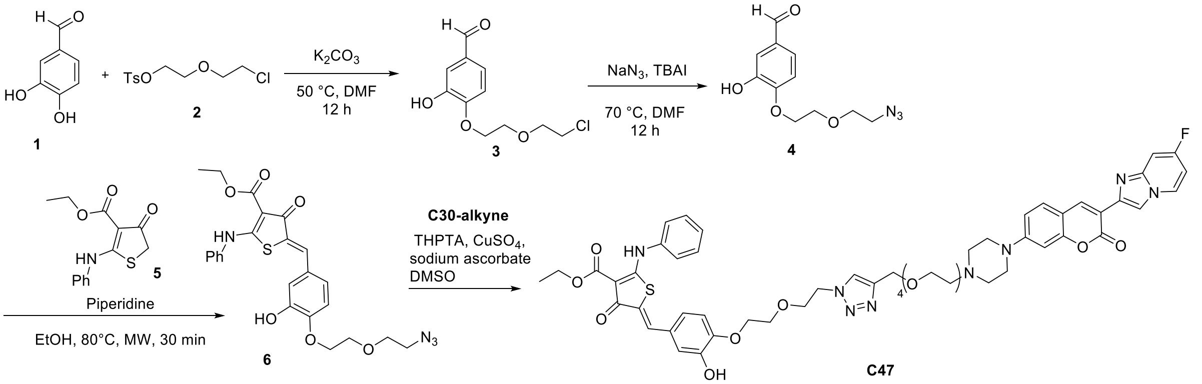


3,4-dihydroxybenzaldehyde **1** (0.1 g, 0.72 mmol) in DMF (2 mL) was added 2-(2-chloroethoxy)ethyl 4-methylbenzenesulfonate **2** (0.19 g, 0.72 mmol) and potassium carbonate (0.1 g, 0.72 mmol). The reaction mixture was stirred at 50 °C for overnight. Added water to the reaction mixture, extracted with diethyl ether. The organic phase was washed with brine solution, dried ($Na_2SO_4$) and concentrated in vacuo to afford compound **3** as a colorless oil (0.15 g, 85%). MS-ESI (*m/z*) $[M+1]^+$ 245.00, 247.01.

Compound **3** (0.15 g, 0.56 mmol) in DMF (2 mL) was added sodium azide (0.06 g, 0.83 mmol) and tetrabutyl ammonium iodide (0.23 g, 0.11 mmol). The reaction mixture was stirred at 70 °C for overnight. Added water to the reaction mixture, extracted with diethyl ether. The organic phase was washed with brine solution, dried ($Na_2SO_4$) and concentrated in vacuo. The crude product was purified by silica gel column chromatography (0 – 18% EtOAc in hexanes) to afford compound **4** as a yellow oil (0.08 g, 57%). MS-ESI (*m/z*) $[M+1]^+$ 252.08.

$^1$H NMR (400 MHz, Chloroform-*d*) δ 9.88 (s, 1H), 7.48 (d, *J* = 2.0 Hz, 1H), 7.44 (dd, *J* = 8.2, 2.0 Hz, 1H), 7.03 (d, *J* = 8.2 Hz, 1H), 4.37 – 4.29 (m, 2H), 3.98 – 3.88 (m, 2H), 3.83 – 3.75 (m, 2H), 3.50 – 3.42 (m, 2H).

Compound **4** (0.05 g, 0.199 mmol) in ethanol was added ethyl 4-oxo-2-(phenylamino)-4,5-dihydro thiophene-3-carboxylate **5** (0.057 g, 0.218 mmol) and piperidine (3 μL, 0.029 mmol). The reaction mixture was heated under microwave irradiation at 100 °C for 45 min. Evaporated the solvent and added ethanol, heated to 50 °C and filtered and washed with cold ethanol to get compound **6** as a yellow solid, (0.055 g, 57%). MS-ESI (*m/z*) $[M+1]^+$ 497.08.

$^{1}H$ NMR (500 MHz, $CDCl_3$) δ 11.52 (s, 1H), 7.76 (s, 1H), 7.55 – 7.48 (m, 2H), 7.41 (d, *J* = 7.7 Hz, 3H), 7.16 (d, *J* = 2.1 Hz, 1H), 7.06 (dd, *J* = 8.3, 2.2 Hz, 1H), 6.93 (d, *J* = 8.3 Hz, 1H), 4.44 (q, *J* = 7.1 Hz, 2H), 4.25 (dd, *J* = 5.3, 3.7 Hz, 2H), 3.91 – 3.84 (m, 2H), 3.77 (t, *J* = 4.8 Hz, 2H), 3.45 (t, *J* = 4.8 Hz, 2H), 1.47 (t, *J* = 7.0 Hz, 3H).

$^{13}C$ NMR (126 MHz, DMSO-$d_6$) δ 177.4, 171.5, 162.3, 142.3, 142.1, 132.4, 126.7, 125.2, 123.8, 123.1, 121.2, 119.4, 114.8, 111.2, 108.9, 65.5, 64.6, 64.3, 55.9, 45.8, 9.7.

Compound **6** (9 mg, 0.018 mmol) in DMSO (0.5 mL) was added compound **C30-alkyne** (10 mg, 0.017 mmol), sodium ascorbate (40 µL, 100 mM in water, 0.004 mmol), THPTA (40 µL, 100 mM in water, 0.004 mmol) and $CuSO_4$ (40 µL, 100 mM in water, 0.004 mmol). The reaction vial was sealed, evacuated, and refilled with $N_2$ three times and stirred at room temperature for overnight. DMSO was removed under vacuum and the residue was purified by silica gel column chromatography (0 – 10% $CH_3OH$ in $CH_2Cl_2$) to afford **C47** as a yellow solid (7 mg, 38%). MS-ESI (*m*/*z*) $[M/2+1]^+$ 538.35.

$^{1}H$ NMR (500 MHz, DMSO-$d_6$) δ 11.24 (s, 1H), 9.44 (s, 1H), 8.71 – 8.68 (m, 2H), 8.52 (s, 1H), 8.05 (s, 1H), 7.67 (d, *J* = 8.8 Hz, 1H), 7.56 – 7.43 (m, 6H), 7.38 (d, *J* = 7.8 Hz, 1H), 7.01 – 6.95 (m, 5H), 6.86 (s, 1H), 4.54 (t, *J* = 5.2 Hz, 2H), 4.47 (s, 2H), 4.27 (q, *J* = 7.1 Hz, 2H), 4.12 – 4.07 (m, 3H), 3.88 (t, *J* = 5.2 Hz, 2H), 3.75 (t, *J* = 5.2 Hz, 2H), 3.56 – 3.47 (m, 14H), 3.18 (d, *J* = 5.1 Hz, 3H), 2.57-2.55 (m, 4H), 1.29 (t, *J* = 7.1 Hz, 3H).

$^{13}C$ NMR (126 MHz, DMSO-$d_6$) δ 180.9, 175.3, 164.9, 160.2 (d, *J* = 252.0 Hz), 159.5, 154.9, 153.3, 148.6, 147.0, 143.8, 139.5, 138.6, 137.5, 130.0, 129.6, 129.5, 129.2 (d, *J* = 11.9 Hz), 128.1, 126.4, 125.5, 124.7, 124.3, 123.1, 115.9, 114.5, 113.7, 112.1, 111.7, 110.2, 104.2 (d, *J* = 30.2 Hz), 99.6, 99.4, 96.9, 69.8, 69.7, 69.6, 68.9, 68.8, 68.6, 67.8, 63.4, 59.5, 52.7, 49.3, 48.6, 14.4.

**Compound C48**

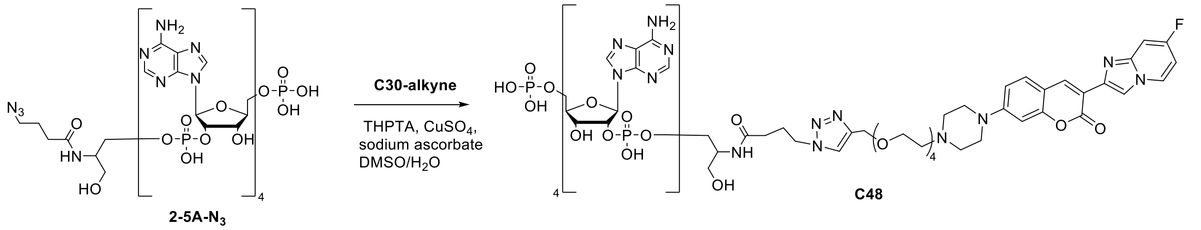


Compound **C30-alkyne** (2 mg, 0.003 mmol) and **2-5A-N$_3$** (WuXi AppTec, 5 mg, 0.003 mmol) in DMSO/$H_2O$ (0.4 mL/0.4 mL) was added sodium ascorbate (15 µL, 100 mM in water, 0.0015 mmol), THPTA (15 µL, 100 mM in water, 0.0015 mmol) and $CuSO_4$ (15 µL, 100 mM in water, 0.0015 mmol). The reaction vial was sealed, evacuated, and refilled with $N_2$ three times and stirred at room temperature for overnight. Solvent was removed by freeze-dry and the residue was purified by reverse-phase column chromatography (0 – 98% acetonitrile in 0.1% ammonium formate water solution) to afford **C48** as a yellow solid (2 mg, 26%). MS-ESI (*m*/*z*) $[(M+n)/n]^+$ 726.74 (n=3), 545.33 (n=4), 435.25 (n=5). HPLC purity: 96.3%.

**Ethyl (*Z*)-5-(3-hydroxy-4-(2-(2-(4-(13-((2-(2-oxo-7-(piperazin-1-yl)-2H-chromen-3-yl)imidazo[1,2-a]pyridin-7-yl)oxy)-2,5,8,11-tetraoxatridecyl)-1H-1,2,3-triazol-1-yl)ethoxy)ethoxy)benzylidene)-4-oxo-2-(phenylamino)-4,5-dihydrothiophene-3-carboxylate (C64)**

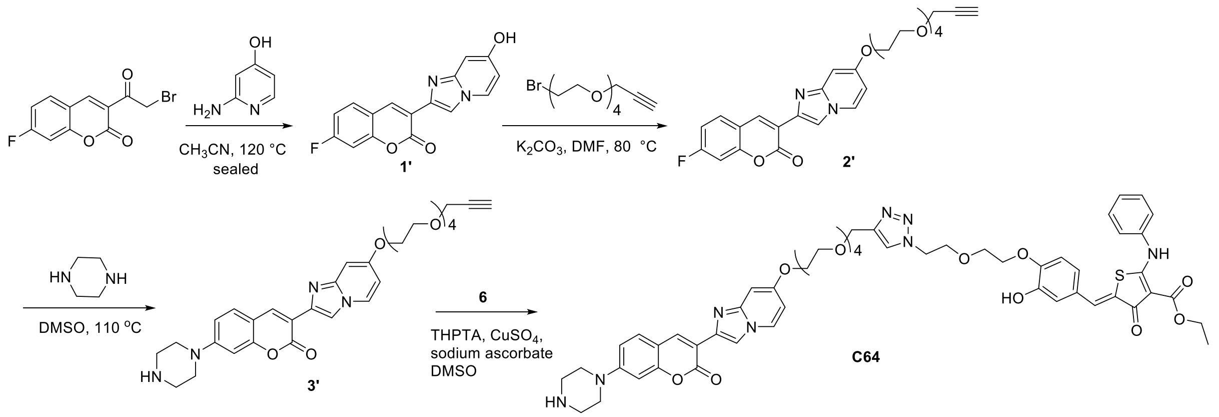

Following general procedures step 1, from 3-(2-bromoacetyl)-7-fluoro-2H-chromen-2-one (200 mg, 0.7 mmol) and 2-aminopyridin-4-ol (77 mg, 0.7 mmol), 50 mg of compound **1'** was obtained as a yellow solid (yield, 24%). MS-ESI (*m*/*z*) $[M+1]^+$ 297.03.

Compound **1'** (41 mg, 0.14 mmol) in DMF (1 mL) was added $K_2CO_3$ (37 mg, 0.27 mmol) and Propargyl-PEG$_4$-Br (60 mg, 0.21 mmol). The reaction mixture was heated at 80 °C for 6 h. The reaction mixture was added ice-water and extracted with EtOAc. The organic phase was washed with brine solution, dried ($Na_2SO_4$) and concentrated in vacuo. The crude product was purified by silica gel column chromatography (0 – 5% $CH_3OH$ in $CH_2Cl_2$) to afford compound **2'** as a yellow solid (21 mg, 30%). MS-ESI (*m*/*z*) $[M+1]^+$ 511.10.

Following general procedures step 2, from compound **2'** (21 mg, 0.7 mmol), 14 mg compound **3'** was obtained as a yellow solid (yield, 59%). MS-ESI (*m*/*z*) $[M+1]^+$ 577.22.

$^1$H NMR (500 MHz, DMSO-*d*$_6$) δ 8.62 (s, 1H), 8.46 (d, *J* = 7.4 Hz, 1H), 8.35 (s, 1H), 7.67 (d, *J* = 9.0 Hz, 1H), 7.01 (d, *J* = 8.9 Hz, 1H), 6.90 (s, 1H), 6.86 (s, 1H), 6.63 (d, *J* = 7.5 Hz, 1H), 4.19 (t, *J* = 5.0 Hz, 2H), 4.14 (d, *J* = 2.4 Hz, 2H), 3.80 (t, *J* = 5.0 Hz, 2H), 3.62 – 3.60 (m, 2H), 3.57 – 3.52 (m, 10H), 3.42 (t, *J* = 2.5 Hz, 1H), 2.31 (t, *J* = 5.1 Hz, 4H), 2.86 (t, *J* = 5.1 Hz, 4H).

$^{13}$C NMR (126 MHz, DMSO-*d*$_6$) δ 159.6, 157.0, 154.7, 153.6, 145.7, 138.1, 137.8, 129.4, 127.8, 115.0, 111.7, 111.2, 110.3, 106.7, 99.3, 94.3, 80.4, 77.1, 69.9, 69.8, 69.7, 69.5, 68.6, 68.5, 67.6, 57.5, 47.8, 45.2.

Compound **3'** (4 mg, 0.007 mmol) in DMSO (0.5 mL) was added compound **6** (3.5 mg, 0.007 mmol), sodium ascorbate (20 μL, 100 mM in water, 0.002 mmol), THPTA (20 μL, 100 mM in water, 0.002 mmol) and $CuSO_4$ (20 μL, 100 mM in water, 0.002 mmol). The reaction vial was sealed, evacuated, and refilled with $N_2$ three times and stirred at room temperature for overnight. DMSO was removed under vacuum and the residue was purified by silica gel column chromatography (0 – 10% $CH_3OH$ in $CH_2Cl_2$) to afford **C64** as a yellow solid (3 mg, 41%). MS-ESI (*m*/*z*) $[M/2+1]^+$ 537.40.

$^1$H NMR (500 MHz, DMSO-*d*$_6$) δ 9.42 (s, 1H), 8.62 (s, 1H), 8.45 (d, *J* = 7.4 Hz, 1H), 8.34 (s, 1H), 8.04 (s, 1H), 7.68 (d, *J* = 8.8 Hz, 1H), 7.50 – 7.47 (m, 2H), 7.39 – 7.33 (m, 4H), 7.03 – 6.88 (m, 6H), 6.61 (dd, *J* = 7.4, 2.5 Hz, 1H), 4.53 (t, *J* = 5.2 Hz, 2H), 4.47 (s, 2H), 4.24 (q, *J* = 7.1 Hz, 2H), 4.16 (t, *J* = 5.2 Hz, 2H), 4.06 (t, *J* = 5.2 Hz, 2H), 3.87 (t, *J* = 5.2 Hz, 2H), 3.78 – 3.73 (m, 4H), 3.59 – 3.47 (m, 16H), 1.28 (t, *J* = 7.1 Hz, 3H).

$^{13}$C NMR (126 MHz, DMSO-*d*$_6$) δ 181.4, 165.2, 159.5, 157.0, 154.6, 148.2, 147.0, 145.7, 143.8, 138.0, 137.7, 129.4, 127.8, 127.0, 124.6, 124.4, 122.7, 115.9, 115.4, 113.8, 111.9, 111.3, 110.6, 106.8, 99.8, 96.5, 94.3, 69.9, 69.8, 69.8, 69.7, 68.9, 68.8, 68.6, 67.8, 67.6, 63.4, 59.2, 49.3, 14.5.

**Compound C65**

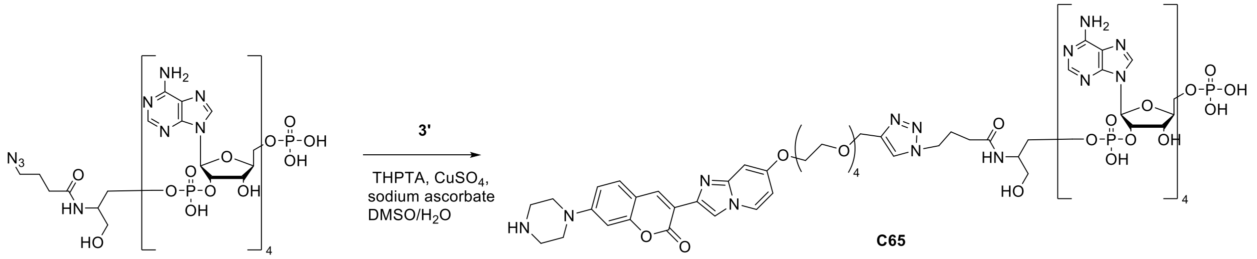


Compound **3'** (3 mg, 0.005 mmol) and **2-5A-N₃** (WuXi AppTec, 8 mg, 0.005 mmol) in DMSO/$H_2O$ (0.5 mL/0.5 mL) was added sodium ascorbate (25 μL, 100 mM in water, 0.0025 mmol), THPTA (25 μL, 100 mM in water, 0.0025 mmol) and $CuSO_4$ (25 μL, 100 mM in water, 0.0025 mmol). The reaction vial was sealed, evacuated, and refilled with $N_2$ three times and stirred at room temperature for overnight. Solvent was removed by freeze-dry and the residue was purified by reverse-phase column chromatography (0 – 98% acetonitrile in 0.1% ammonium formate water solution) to afford **C65** as a yellow solid (3 mg, 26%). MS-ESI (*m*/*z*) $[(M+n)/n]^+$ 726.29 (n=3), 545.00 (n=4). HPLC purity: 91.2%.

**7-(4-(2-(3-(but-3-yn-1-yl)-3*H*-diazirin-3-yl)ethyl)piperazin-1-yl)-3-(7-fluoroimidazo[1,2-*a*]pyridin-2-yl)-2*H*-chromen-2-one (C30-D)**

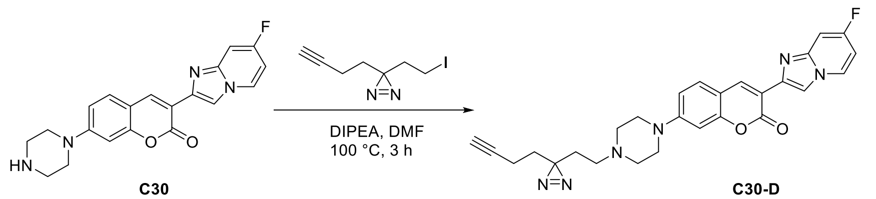


Compound **C30** (10 mg, 0.027 mmol) in DMF (0.5 mL) was added 3-(but-3-yn-1-yl)-3-(2-iodoethyl)-3*H*-diazirine (7 mg, 0.028 mmol) and DIPEA (14 μL, 0.081 mmol). The mixture was stirred at 100 °C for 3 h. The mixture was diluted with DCM and water. The organic phase was washed with brine solution, dried ($Na_2SO_4$) and concentrated in vacuo. The crude product was purified by silica gel column chromatography (0 – 10% $CH_3OH$ in $CH_2Cl_2$) to afford compound **C30-D** as a yellow solid (7 mg, 52%). MS-ESI (m/z) $[M+1]^+$ 485.19. HPLC purity: >98%.

$^1$H NMR (500 MHz, DMSO-*d*$_6$) δ 8.63 – 8.59 (m, 2H), 8.44 (s, 1H), 7.60 (d, *J* = 8.8 Hz, 1H), 7.30 (dd, *J* = 10.1, 2.6 Hz, 1H), 6.94 (dd, *J* = 8.9, 2.4 Hz, 1H), 6.89 (td, *J* = 7.6, 2.6 Hz, 1H), 6.79 (d, *J* = 2.4 Hz, 1H), 2.77 (t, *J* = 2.6 Hz, 1H), 2.40 (t, *J* = 5.1 Hz, 4H), 2.12 (t, *J* = 7.4 Hz, 2H), 1.96 (td, *J* = 7.4, 2.7 Hz, 2H), 1.55 – 1.51 (m, 4H).

$^{13}$C NMR (126 MHz, DMSO-*d*$_6$) δ 160.6 (d, *J* = 239.4 Hz), 159.9, 155.3, 153.8, 144.8 (d, *J* = 14.6 Hz), 139.8, 139.0, 130.0, 129.7 (d, *J* = 11.6 Hz), 115.0, 112.4 (d, *J* = 42.5 Hz), 110.7, 104.6 (d, *J* = 29.8 Hz), 100.1, 99.9, 83.8, 72.2, 52.6, 52.5, 47.3, 32.2, 29.9, 28.1, 13.2.

***N*-((*S*)-1-((2-(4-(3-(7-fluoroimidazo[1,2-*a*]pyridin-2-yl)-2-oxo-2*H*-chromen-7-yl)piperazin-1-yl)ethyl)amino)-3-(3-methyl-3*H*-diazirin-3-yl)-1-oxopropan-2-yl)-1-(5-((3a*S*,4*S*,6a*R*)-2-oxohexahydro-1*H*-thieno[3,4-*d*]imidazol-4-yl)pentanamido)-3,6,9,12-tetraoxapentadecan-15-amide (C30-BD)**

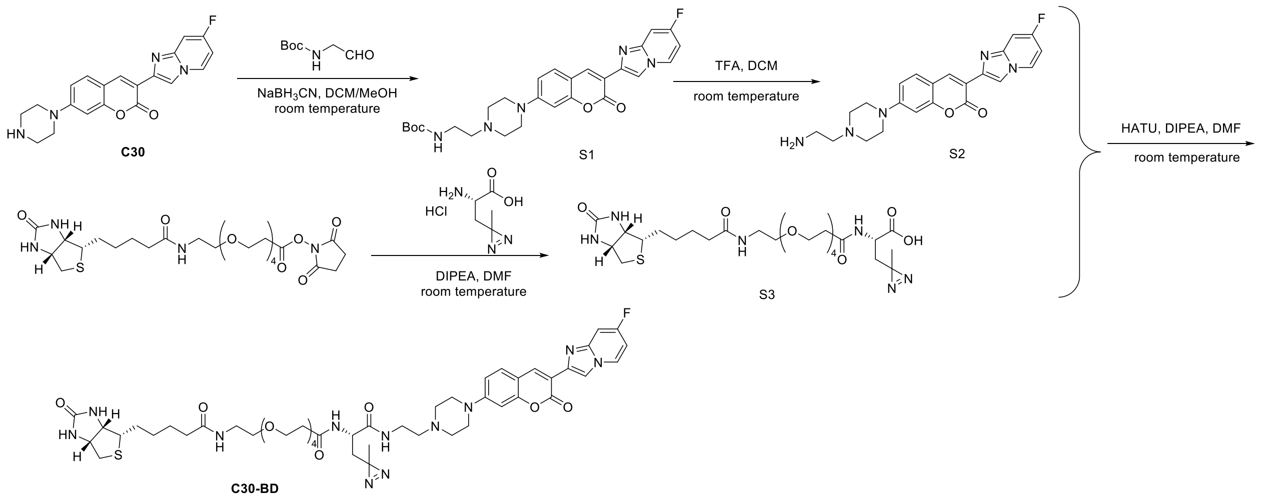

Compound **C30** (40 mg, 0.11 mmol) in DCM (0.5 mL) and MeOH (0.5 mL) was cooled with ice. Acetaldehyde (32 μL, 0.55 mmol) was added, followed by sodium cyanoborohydride (35 mg, 0.55 mmol). The mixture was stirred at room temperature for 2 h. The mixture was diluted with DCM and water was added dropwise to quench the reaction. The organic phase was washed with brine solution, dried ($Na_2SO_4$) and concentrated in vacuo. The crude product was purified by silica gel column chromatography (0 – 5% $CH_3OH$ in $CH_2Cl_2$) to afford compound **S1** as a yellow solid (40 mg, 72%). MS-ESI (m/z) $[M+1]^+$ 508.13.

Compound **S1** was dissolved in DCM (1 mL), TFA (0.5 mL) was added. The mixture was stirred at room temperature for 1 h. DCM and TFA were removed under vacuum. The residue was dissolved in DMF (1 mL) and used directly in the amide coupling reaction without purification.

Biotin-NHS (30 mg, 0.051 mmol) was dissolved in DMF (1 mL), (*S*)-2-amino-3-(3-methyl-3H-diazirin-3-yl) propanoic acid hydrochloride (10 mg, 0.055 mmol) and DIPEA (44 μL, 0.25 mmol) were added. The mixture was stirred at room temperature overnight. The mixture was cooled with ice, HATU (28 mg, 0.075 mmol) was added, and stirred at room temperature for 15 min. Compound **S2** in DMF was added and the mixture was stirred at room temperature overnight. The mixture was diluted with ethyl acetate and water. The organic phase was washed with brine solution, dried ($Na_2SO_4$) and concentrated in vacuo. The crude product was purified by silica gel column chromatography (0 – 10% $CH_3OH$ in $CH_2Cl_2$) to afford compound **C30-BD** as a yellow solid (10 mg, 20%). MS-ESI (m/z) $[(M+2)/2]^+$ 503.86. HPLC purity: >98%.

$^1H$ NMR (500 MHz, DMSO-$d_6$) δ 8.71 – 8.68 (m, 2H), 8.52 (s, 1H), 8.14 (d, *J* = 8.4 Hz, 1H), 7.96 (t, *J* = 5.6 Hz, 1H), 7.83 (t, *J* = 5.7 Hz, 1H), 7.70 (d, *J* = 9.0 Hz, 1H), 7.39 (dd, *J* = 10.0, 2.6 Hz, 1H), 7.03 (dd, *J* = 9.0, 2.4 Hz, 1H), 6.98 (td, *J* = 7.6, 2.6 Hz, 1H), 6.88 (d, *J* = 2.4 Hz, 1H), 6.42 (s, 1H), 6.36 (s, 1H), 4.32 – 4.29 (m, 1H), 4.27 – 4.22 (m, 1H), 4.14 – 4.11 (m, 1H), 3.63 (td, *J* = 6.6, 1.4 Hz, 2H), 3.50 – 3.49 (m, 12H), 3.40 – 3.38 (m, 5H), 3.26 – 3.16 (m, 5H), 3.11 – 3.07 (m, 1H), 2.81 (dd, *J* = 12.4, 5.1 Hz, 1H), 2.58 (d, *J* = 12.5 Hz, 1H), 2.53 (t, *J* = 5.2 Hz, 4H), 2.42 (t, *J* = 6.0 Hz, 4H), 2.06 (t, *J* = 7.5 Hz, 2H), 1.75 (dd, *J* = 14.7, 5.0 Hz, 1H), 1.63 – 1.58 (m, 1H), 1.53 – 1.46 (m, 4H), 1.33 – 1.27 (m, 2H), 1.01 (s, 3H).

$^{13}C$ NMR (126 MHz, DMSO-$d_6$) δ 172.6, 171.0, 170.4, 163.2, 160.7 (d, *J* = 252 Hz), 160.0, 155.4, 153.8, 144.8 (d, *J* = 14.8 Hz), 139.8, 139.1, 130.1, 129.7 (d, *J* = 11.6 Hz), 114.9, 112.4 (d, *J* = 42.2 Hz), 110.7, 104.6 (d, *J* = 29.7 Hz), 100.1, 99.9, 70.2, 70.1, 70.0, 69.6, 67.2, 61.5, 59.7, 57.0, 55.9, 52.7, 49.1, 47.3, 35.6, 28.7, 28.5, 25.7, 24.8, 20.0.

**(*S*)-2-(4-(4-(bis(2-chloroethyl)amino)phenyl)butanamido)-6-(3-(2-(2-(2-(4-(3-(7-fluoroimidazo[1,2-*a*]pyridin-2-yl)-2-oxo-2*H*-chromen-7-yl)piperazin-1-yl)ethoxy)ethoxy)ethoxy)propanamido)-*N*-(2-(5-((3a*S*,4*S*,6a*R*)-2-oxohexahydro-1*H*-thieno[3,4-*d*]imidazol-4-yl)pentanamido)ethyl)hexanamide (C30-NM)**

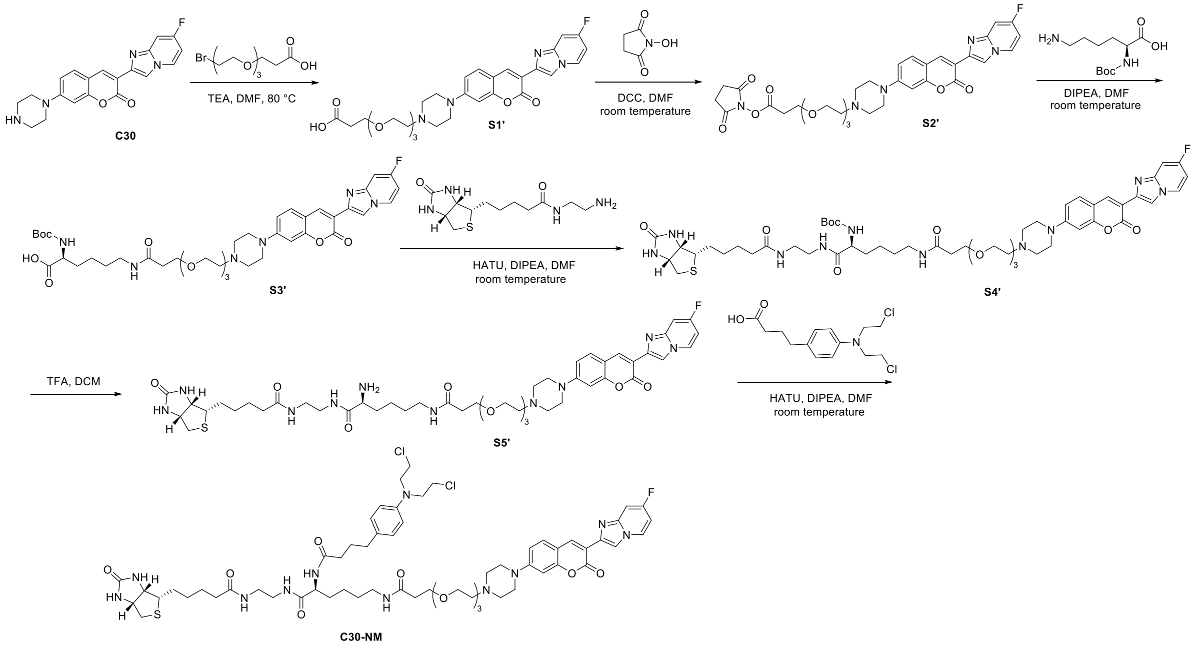


Compound **C30** (50 mg, 0.137 mmol) in DMF (1 mL) was added 3-(2-(2-(2 bromoethoxy)ethoxy)ethoxy) propanoic acid (58 mg, 0.2 mmol) and triethylamine (55 µL, 0.4 mmol). The mixture was stirred at 80 °C overnight. DMF and TEA were removed under vacuum. The crude product was purified by silica gel column chromatography (0 – 15% $CH_3OH$ in $CH_2Cl_2$) to afford compound **S1'** as a yellow solid (55 mg, 70%). MS-ESI (m/z) $[M+1]^+$ 569.21.

Compound **S1'** (55 mg, 0.096 mmol) was dissolved in DMF (1 mL), N-hydroxysuccinimide (17 mg, 0.15 mmol) and DCC (31 mg, 0.15 mmol) were added. The mixture was stirred at room temperature overnight. Precipitate was filtered and DMF was removed under vacuum. The crude product was purified by silica gel column chromatography (0 – 10% $CH_3OH$ in $CH_2Cl_2$) to afford compound **S2'** as a yellow solid (50 mg, 77%). MS-ESI (m/z) $[M+1]^+$ 666.27.

Compound **S2'** (50 mg, 0.075 mmol) was dissolved in DMF (1 mL), Boc-Lys-OH (27 mg, 0.11 mmol) and DIPEA (19 µL, 0.11 mmol) were added. The mixture was stirred at room temperature overnight. DMF and DIPEA were removed under vacuum. The crude product was purified by silica gel column chromatography (0 – 15% $CH_3OH$ in $CH_2Cl_2$) to afford compound **S3'** as a yellow solid (45 mg, 75%). MS-ESI (m/z) $[M+1]^+$ 797.26.

Compound **S3'** (45 mg, 0.056 mmol) was dissolved in DMF (1 mL) and cooled with ice, HATU (27 mg, 0.07 mmol) and DIPEA (24 µL, 0.14 mmol) were added. The mixture was stirred at room temperature for 15 min. Biotin-amine (20 mg, 0.07 mmol) was added and the mixture was stirred at room temperature overnight. DMF and DIPEA were removed under vacuum. The crude product was purified by silica gel column chromatography (0 – 15% $CH_3OH$ in $CH_2Cl_2$) to afford compound **S4'** as a yellow solid (40 mg, 67%). MS-ESI (m/z) $[(M+2)/2]^+$ 533.38.

Compound **S4'** (40 mg, 0.037 mmol) was dissolved in DCM (1 mL) and TFA (0.5 mL). The mixture was stirred at room temperature for 1 h. DCM and TFA were removed under vacuum. The residue was dissolved in DMF (0.5 mL) and used directly in the amide coupling reaction without purification.

Chlorambucil (12 mg, 0.04 mmol) was dissolved in DMF (0.5 mL), HATU (19 mg, 0.05 mmol) and DIPEA (34 µL, 0.2 mmol) were added. The mixture was stirred at room temperature for 15 min. Compound **S5'** in DMF was added and the mixture was stirred at room temperature overnight. DMF and DIPEA were removed under vacuum. The crude product was purified by silica gel column chromatography (0 – 10% $CH_3OH$ in $CH_2Cl_2$) to afford compound **C30-NM** as a yellow solid (20 mg, 43%). MS-ESI (m/z) $[(M+2)/2]^+$ 626.74. HPLC purity: >98%.

$^1$H NMR (500 MHz, DMSO-$d_6$) δ 8.71 – 8.68 (m, 2H), 8.52 (s, 1H), 7.91 – 7.88 (m, 2H), 7.81 (t, *J* = 5.6 Hz, 1H), 7.76 (t, *J* = 5.3 Hz, 1H), 7.69 (d, *J* = 8.9 Hz, 1H), 7.39 (dd, *J* = 10.1, 2.6 Hz, 1H), 7.04 – 6.96 (m, 4H), 6.88 (d, *J* = 2.4 Hz, 1H), 6.66 (s, 1H), 6.65 (s, 1H), 6.43 (s, 1H), 6.36 (s, 1H), 4.30 (dd, *J* = 7.7, 5.1 Hz, 1H), 4.15 – 4.10 (m, 3H), 3.72– 3.66 (m, 8H), 3.59 (t, *J* = 6.5 Hz, 2H), 3.55 (t, *J* = 5.9 Hz, 2H), 3.51 – 3.47 (m, 8H), 3.17 (d, *J* = 5.2 Hz, 1H), 3.10 – 3.07 (m, 5H), 3.03 – 2.98 (m, 2H), 2.81 (dd, *J* = 12.4, 5.1 Hz, 1H), 2.59 – 2.53 (m, 7H), 2.42 (t, *J* = 8.0 Hz, 2H), 2.29 (t, *J* = 6.5 Hz, 2H), 2.14 (t, *J* = 7.5 Hz, 2H), 2.04 (t, *J* = 7.5 Hz, 2H), 1.75 – 1.69 (m, 2H), 1.64 – 1.57 (m, 3H), 1.52 – 1.44 (m, 4H), 1.39 – 1.24 (m, 7H).

$^{13}$C NMR (126 MHz, DMSO-$d_6$) δ 172.8, 172.6, 172.5, 170.3, 163.2, 160.0, 155.4, 153.9, 144.8, 144.7, 139.9, 139.1, 130.5, 130.0, 129.8, 129.6, 114.9, 112.5, 112.3, 112.2, 110.7, 100.1, 99.9, 70.2, 70.0, 68.8, 67.3, 61.5, 59.7, 57.6, 55.8, 53.2, 52.7, 47.3, 35.7, 35.2, 34.1, 32.0, 29.3, 28.6, 28.5, 27.8, 25.6, 23.4.

### NMR and HPLC Spectra
# C30

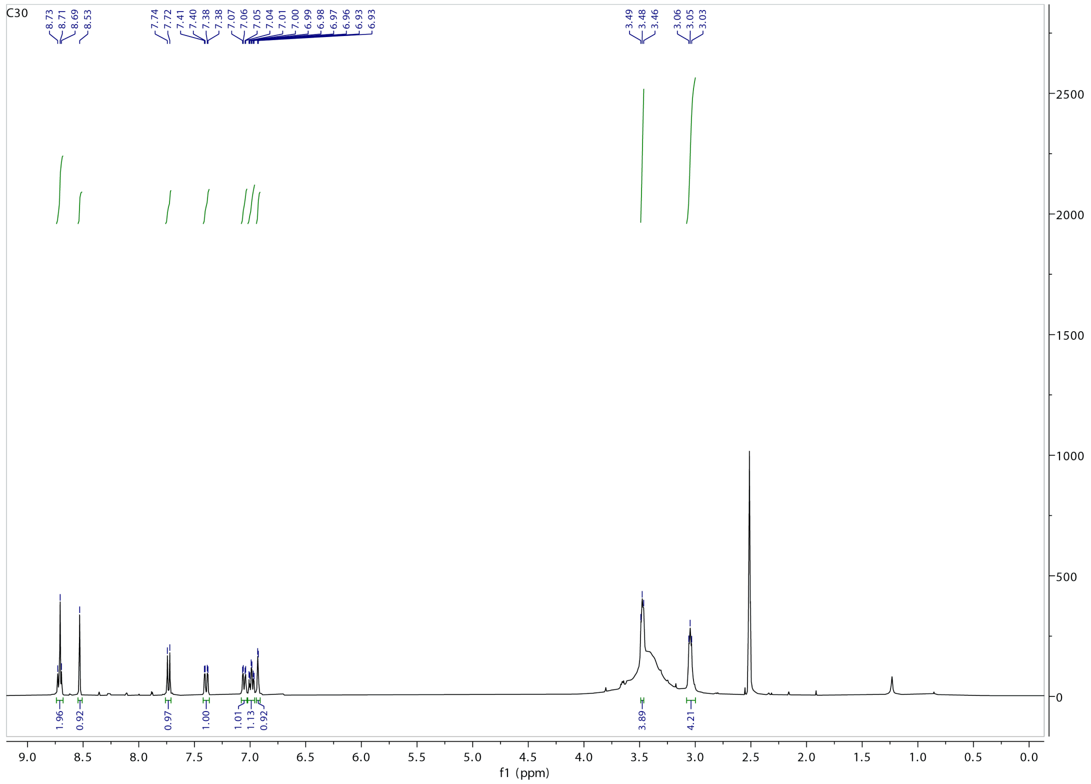

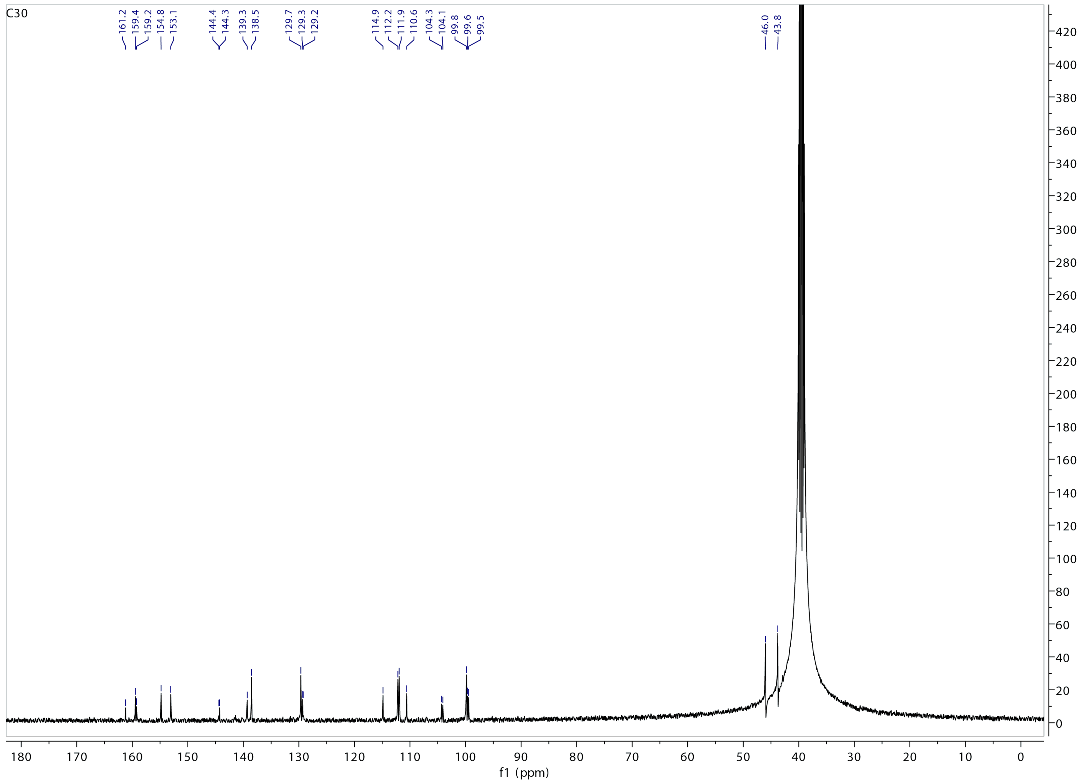

# C31

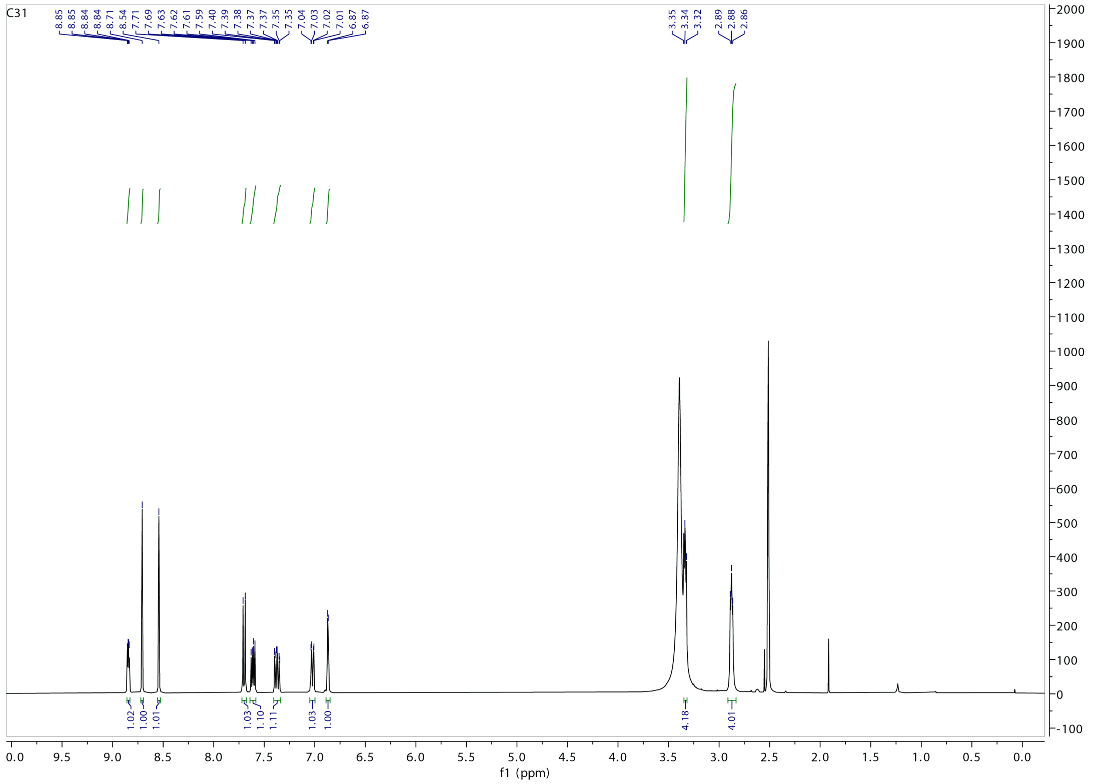
C31
8.85
8.85
8.84
8.84
8.71
8.54
7.71
7.69
7.63
7.62
7.61
7.59
7.40
7.39
7.38
7.37
7.37
7.35
7.04
7.03
7.02
7.01
6.87
6.87
3.35
3.34
3.32
2.89
2.88
2.86
1.02
1.00
1.01
1.03
1.10
1.11
1.03
1.00
4.18
4.01
f1 (ppm)

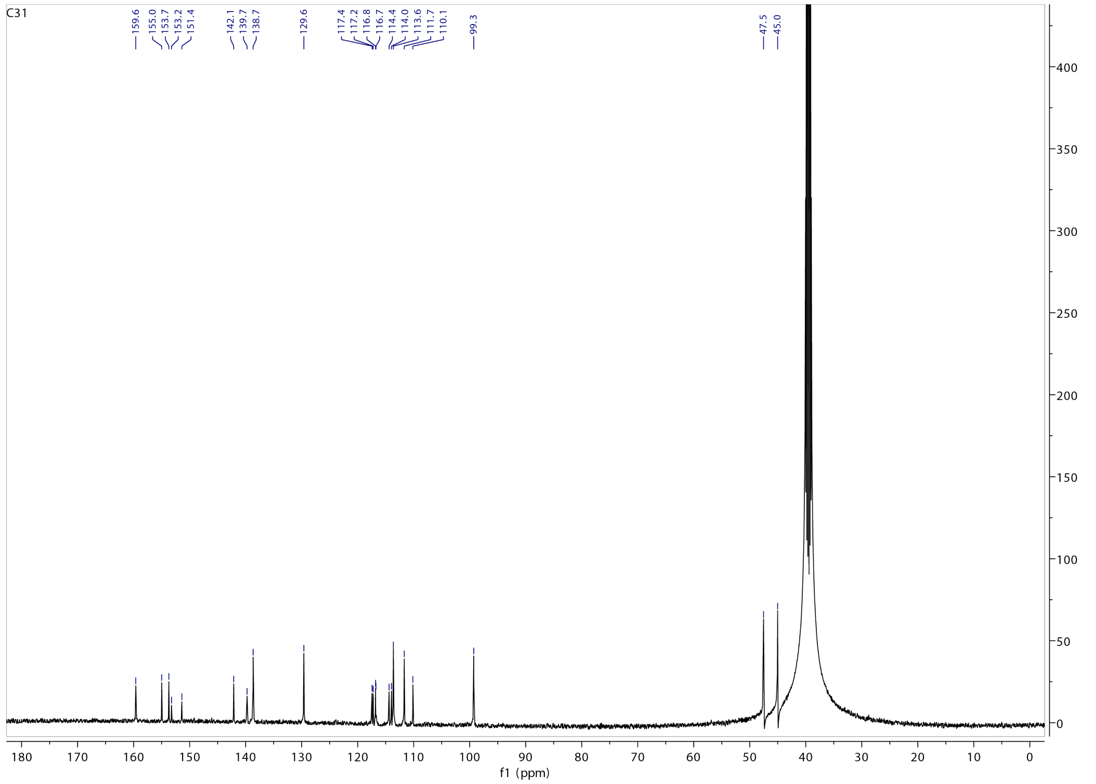
C31
159.6
155.0
153.7
153.2
151.4
142.1
139.7
138.7
129.6
117.4
117.2
116.8
116.7
114.4
114.0
113.6
111.7
110.1
99.3
47.5
45.0
f1 (ppm)

**C32**

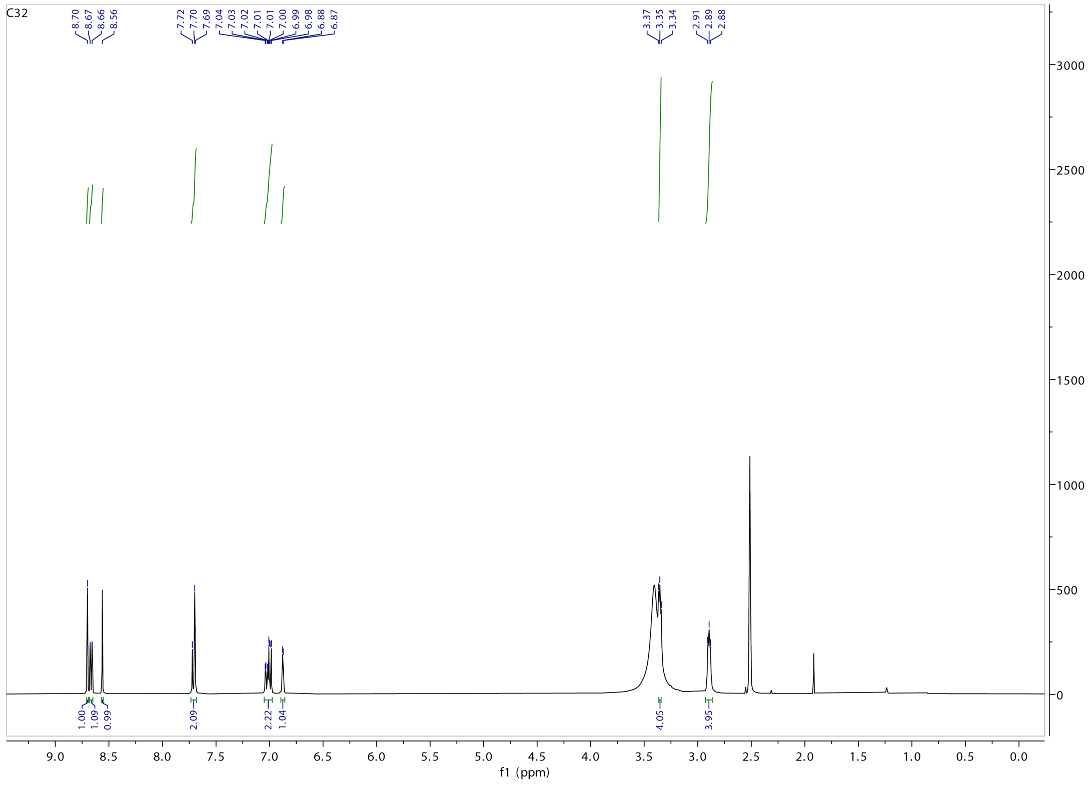
C32
8.70
8.67
8.66
8.56
7.72
7.70
7.69
7.04
7.03
7.02
7.01
7.01
7.00
6.99
6.98
6.88
6.87
3.37
3.35
3.34
2.91
2.89
2.88
1.00
1.09
0.99
2.09
2.22
1.04
4.05
3.95
f1 (ppm)

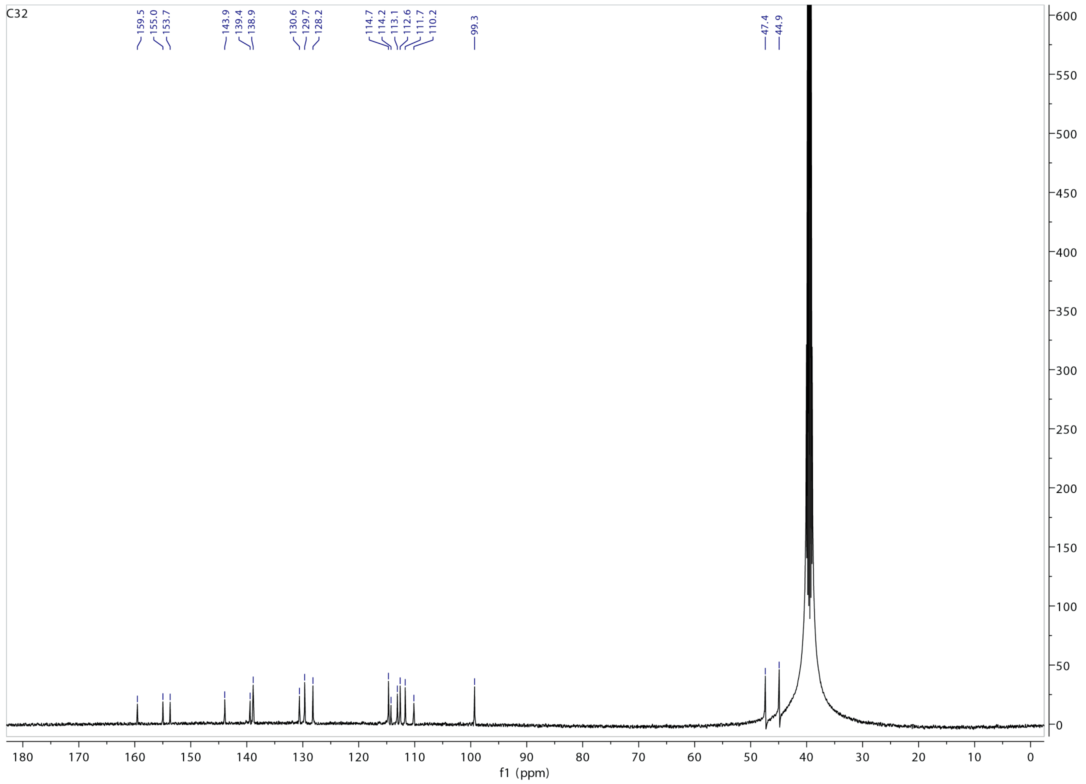
C32
159.5
155.0
153.7
143.9
139.4
138.9
130.6
129.7
128.2
114.7
114.2
113.1
112.6
111.7
110.2
99.3
47.4
44.9
f1 (ppm)

# C36

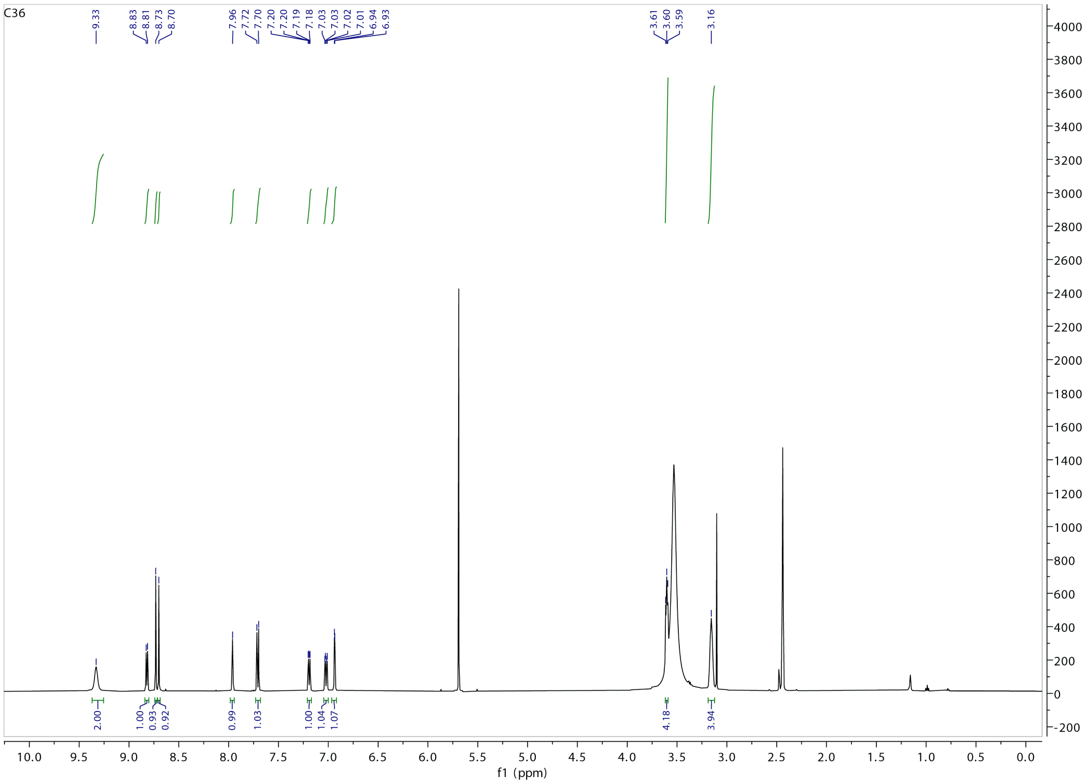
C36
9.33
8.83
8.81
8.73
8.70
7.96
7.72
7.70
7.20
7.20
7.19
7.18
7.03
7.03
7.02
7.01
6.94
6.93
3.61
3.60
3.59
3.16
2.00
1.00
0.93
0.92
0.99
1.03
1.00
1.04
1.07
4.18
3.94
f1 (ppm)

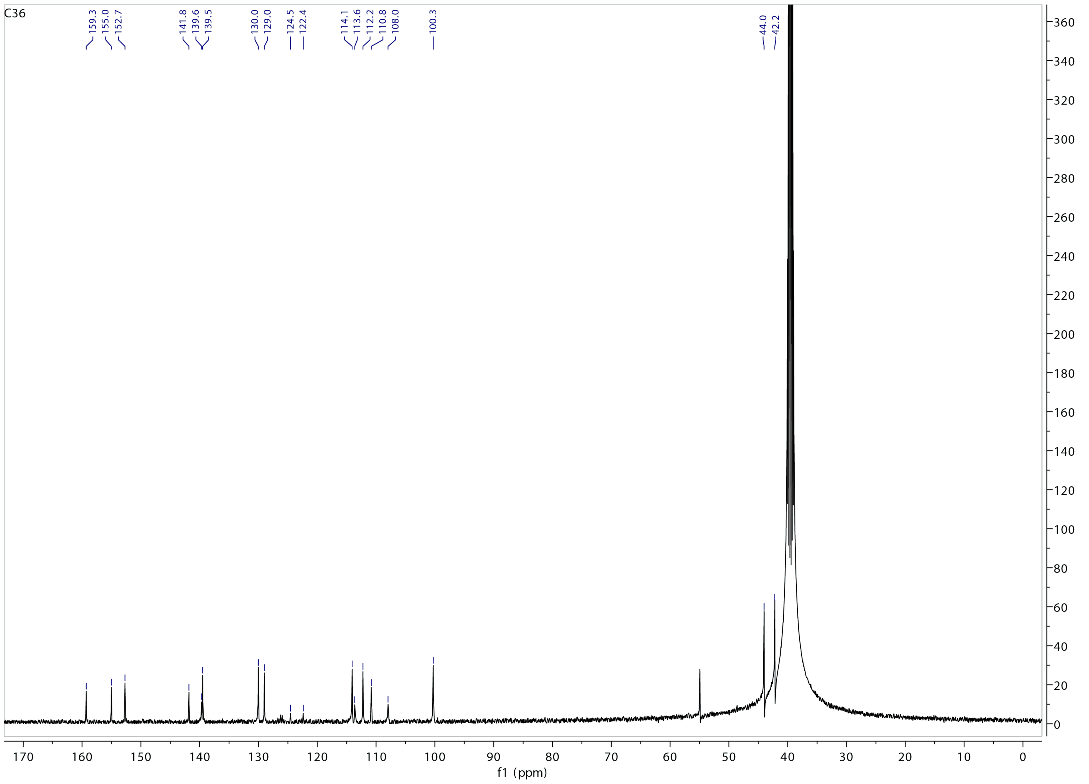
C36
159.3
155.0
152.7
141.8
139.6
139.5
130.0
129.0
124.5
122.4
114.1
113.6
112.2
110.8
108.0
100.3
44.0
42.2
f1 (ppm)

# C34

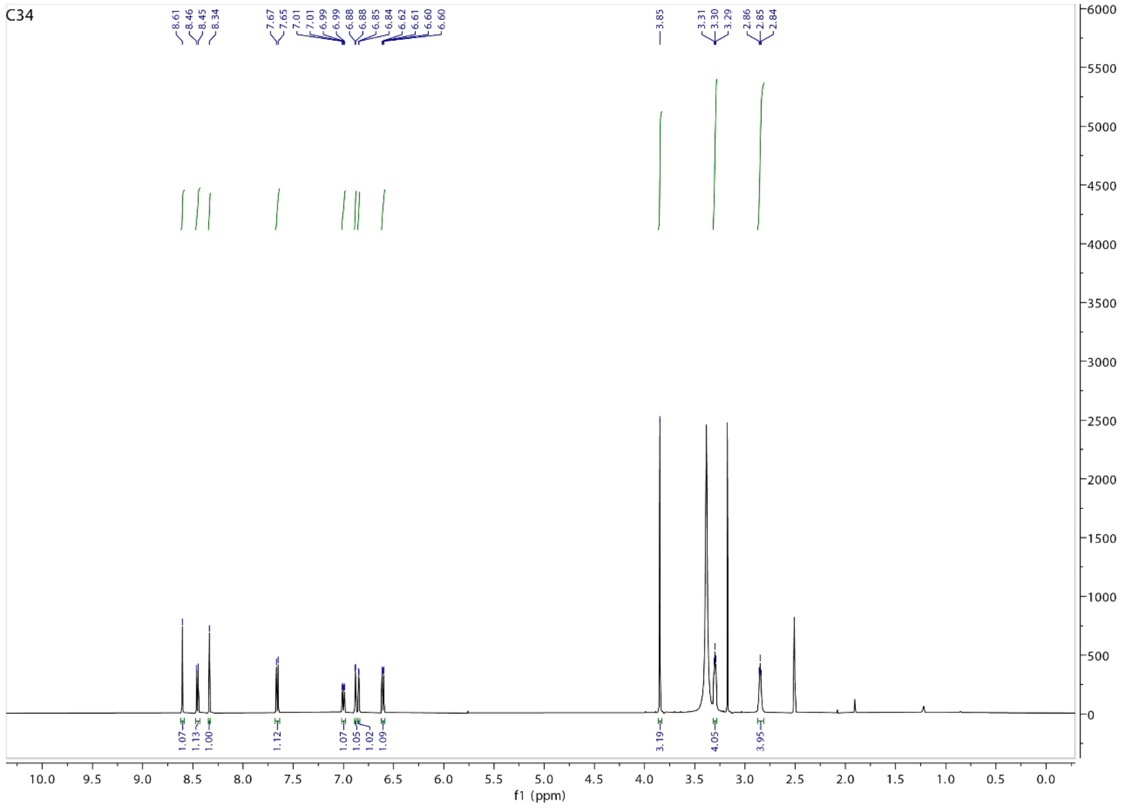

C34
8.61
8.46
8.45
8.34
7.67
7.65
7.01
7.01
6.99
6.99
6.88
6.88
6.85
6.84
6.62
6.61
6.60
6.60
3.85
3.31
3.30
3.29
2.86
2.85
2.84
1.07
1.13
1.00
1.12
1.07
1.05
1.02
1.09
3.19
4.05
3.95
f1 (ppm)


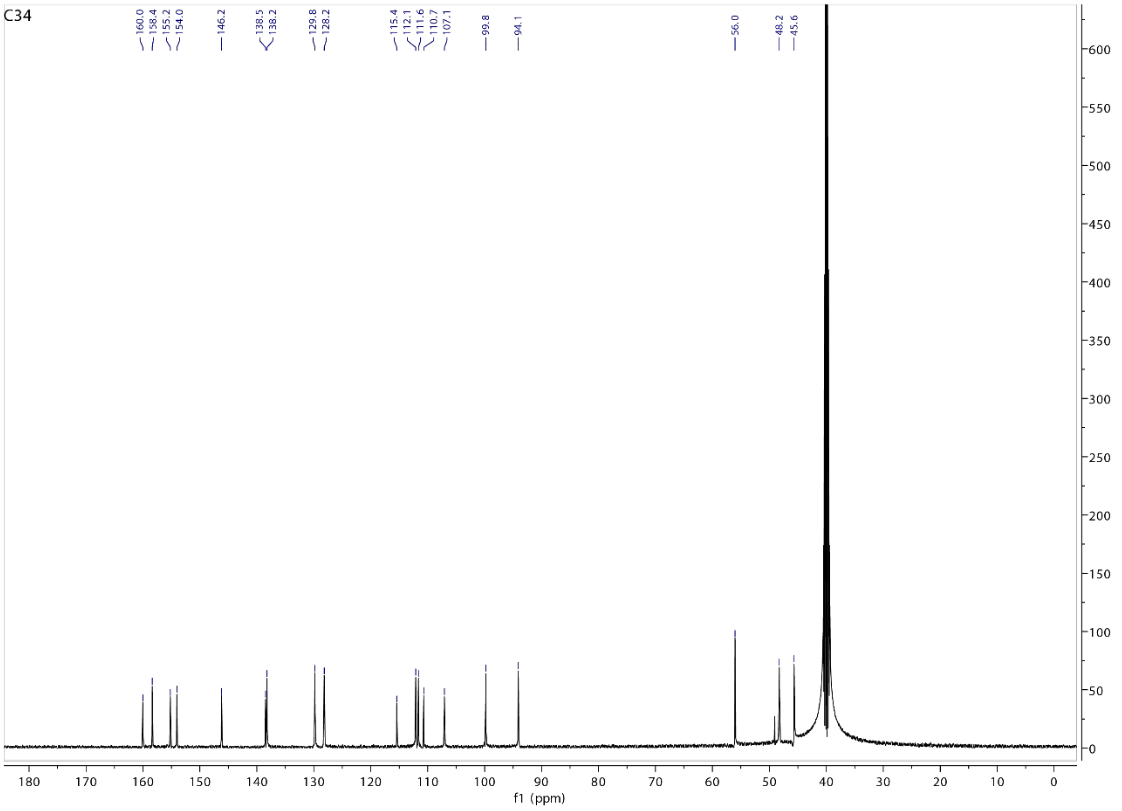

C34
160.0
158.4
155.2
154.0
146.2
138.5
138.2
129.8
128.2
115.4
112.1
111.6
110.7
107.1
99.8
94.1
56.0
48.2
45.6
f1 (ppm)

#### C30-alkyne

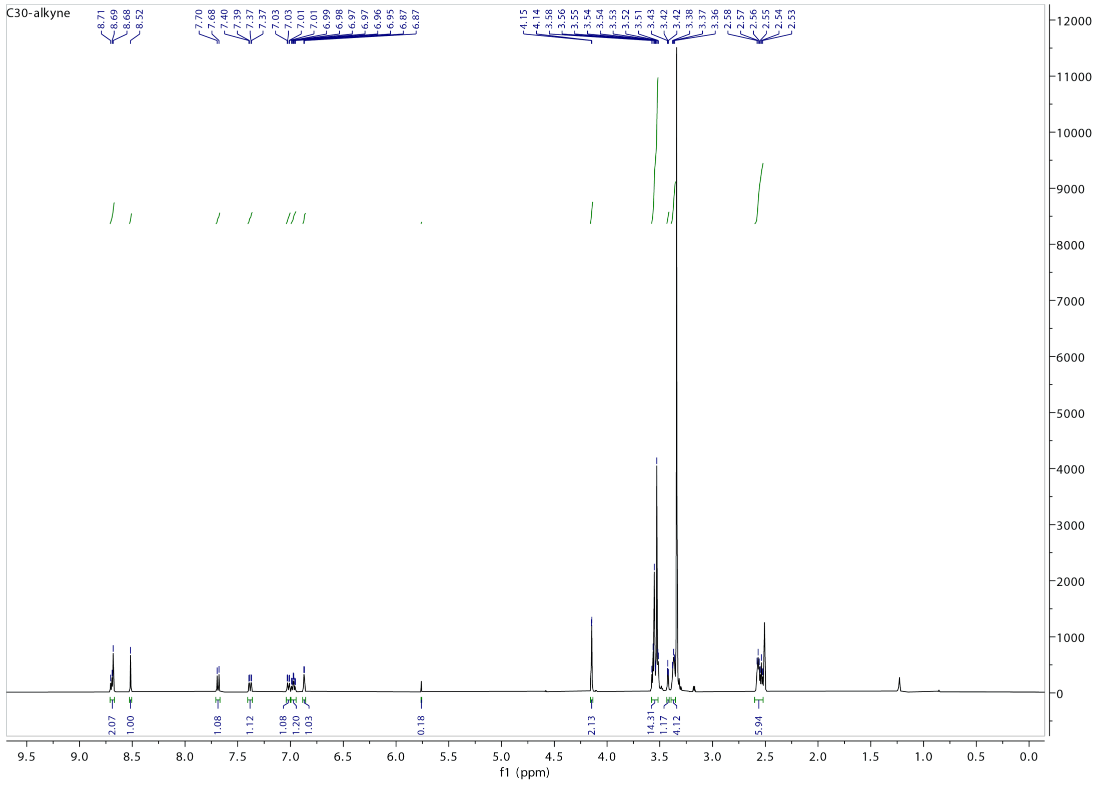

C30-alkyne
8.71 8.69 8.68 8.52 7.70 7.68 7.40 7.39 7.37 7.37 7.03 7.03 7.01 6.99 6.98 6.97 6.97 6.96 6.95 6.87 6.87 4.15 4.14 3.58 3.56 3.55 3.54 3.54 3.53 3.52 3.51 3.43 3.42 3.42 3.38 3.37 3.36 2.58 2.57 2.56 2.55 2.54 2.53
2.07 1.00 1.08 1.12 1.08 1.20 1.03 0.18 2.13 14.31 1.17 4.12 5.94
f1 (ppm)


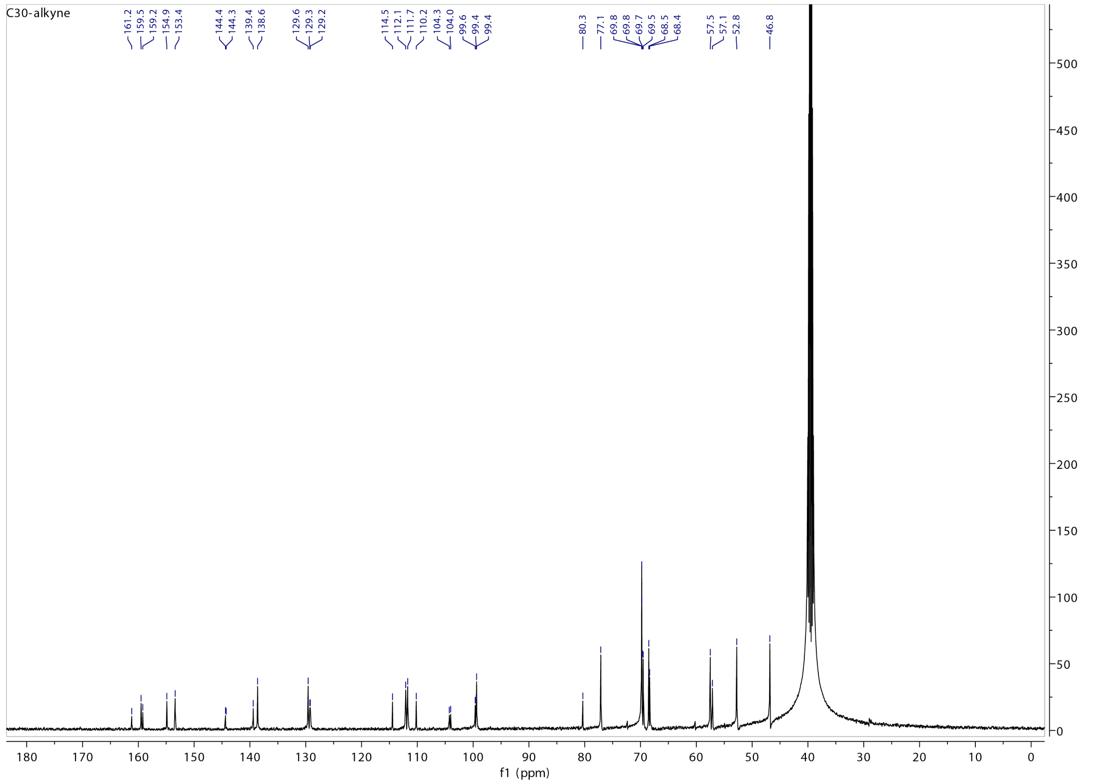

C30-alkyne
161.2 159.5 159.2 154.9 153.4 144.4 144.3 139.4 138.6 129.6 129.3 129.2 114.5 112.1 111.7 110.2 104.3 104.0 99.6 99.4 99.4 80.3 77.1 69.8 69.8 69.7 69.5 68.5 68.4 57.5 57.1 52.8 46.8
f1 (ppm)

### C30-FCA

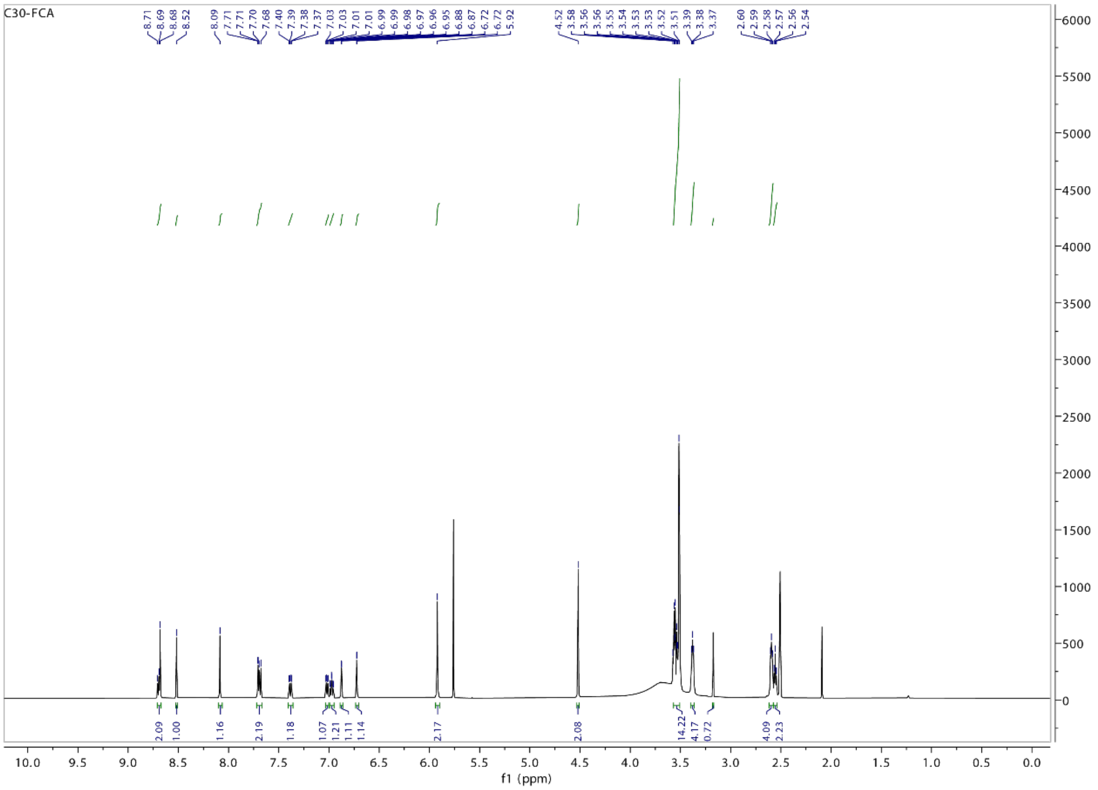


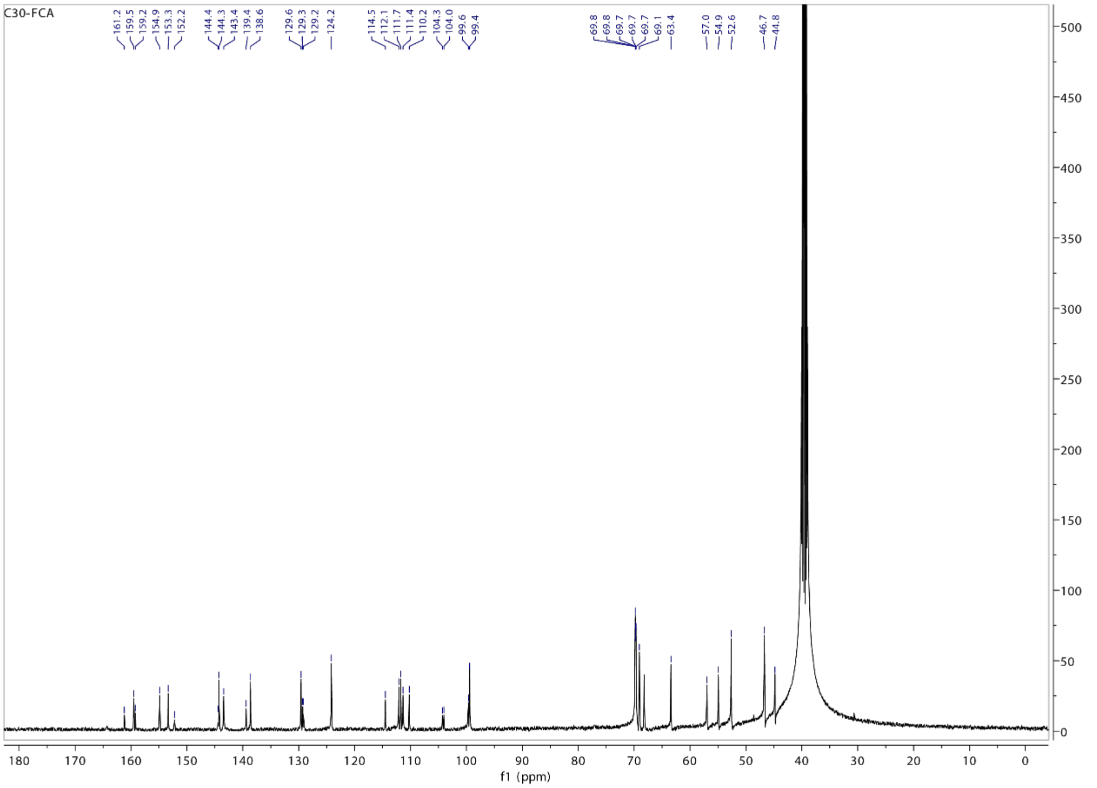

### Compound 6

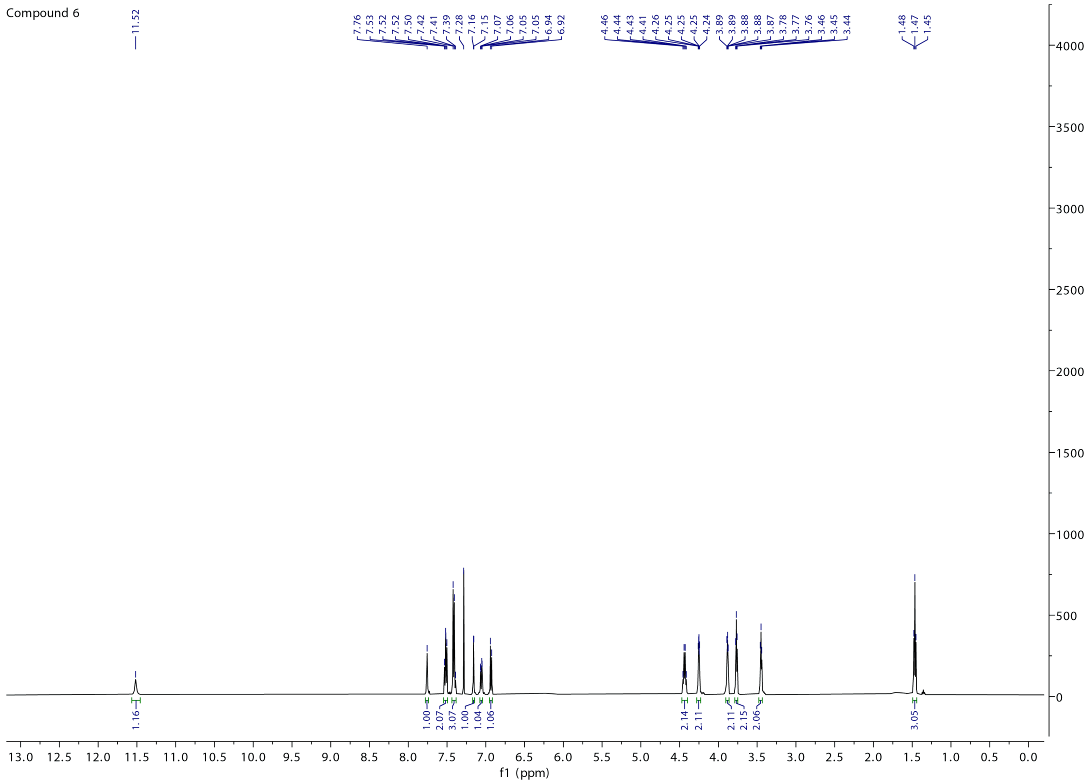
Compound 6
11.52
7.76
7.53
7.52
7.52
7.50
7.42
7.41
7.39
7.28
7.16
7.15
7.07
7.06
7.05
7.05
6.94
6.92
4.46
4.44
4.43
4.41
4.26
4.25
4.25
4.25
4.24
3.89
3.89
3.88
3.88
3.87
3.78
3.77
3.76
3.46
3.45
3.44
1.48
1.47
1.45
1.16
1.00
2.07
3.07
1.00
1.04
1.06
2.14
2.11
2.11
2.15
2.06
3.05
f1 (ppm)

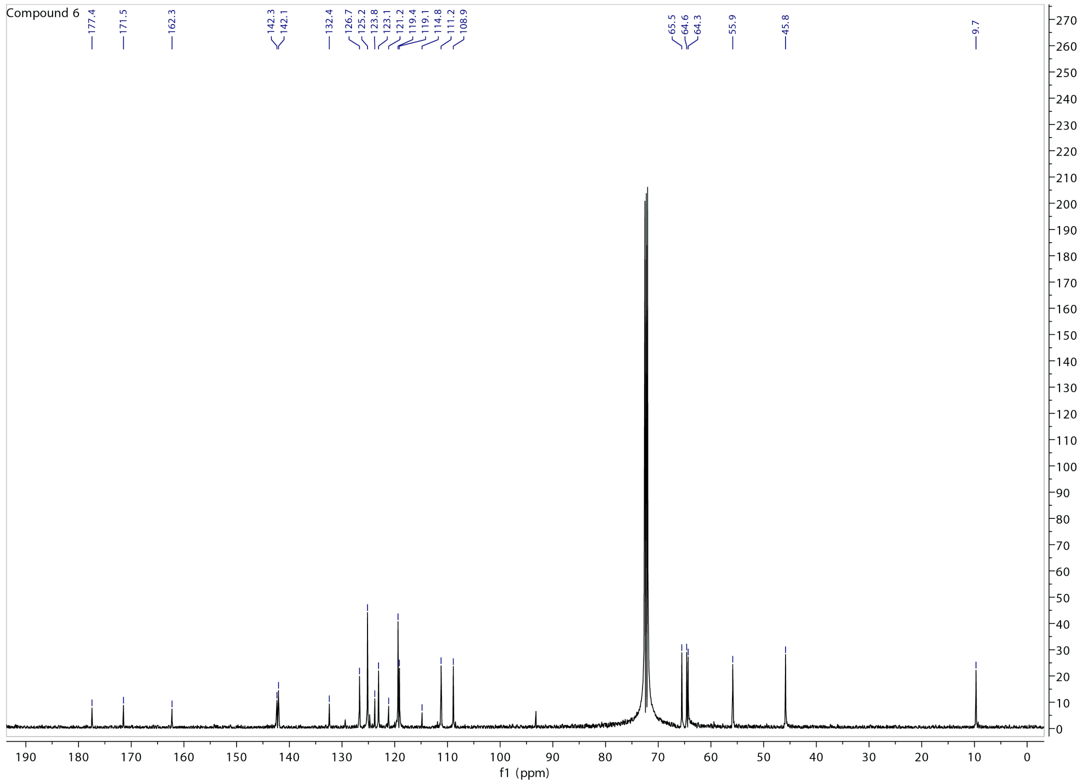
Compound 6
177.4
171.5
162.3
142.3
142.1
132.4
126.7
125.2
123.8
123.1
121.2
119.4
119.1
114.8
111.2
108.9
65.5
64.6
64.3
55.9
45.8
9.7
f1 (ppm)

# C47

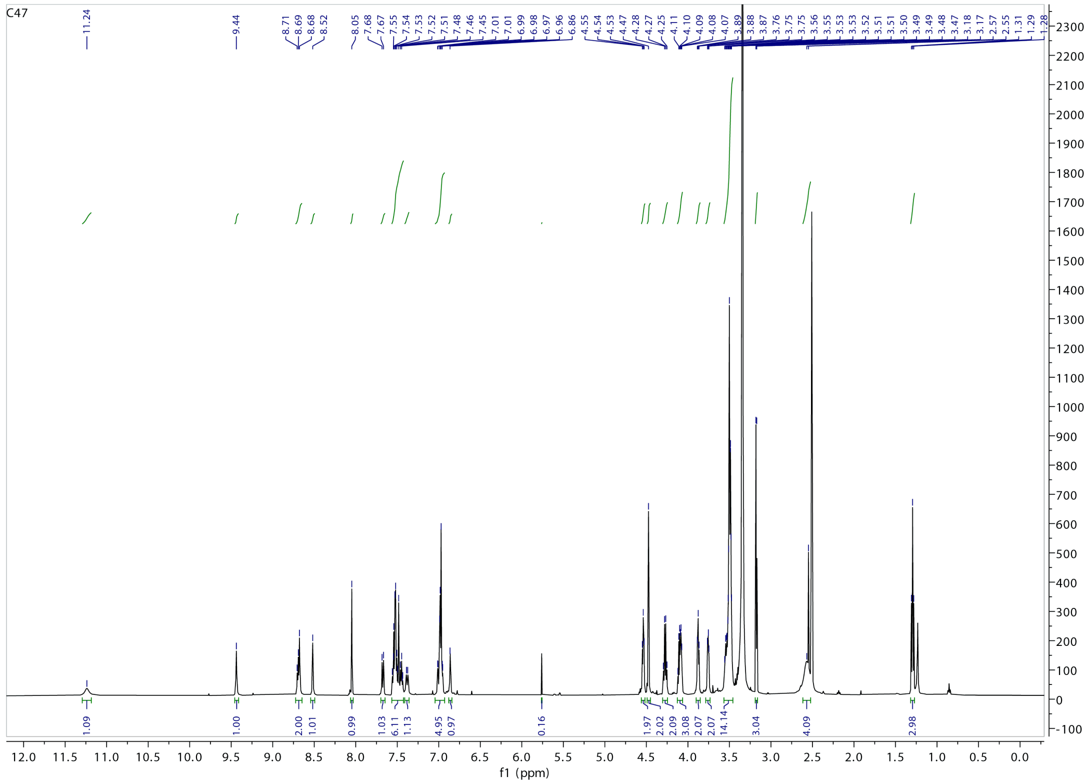

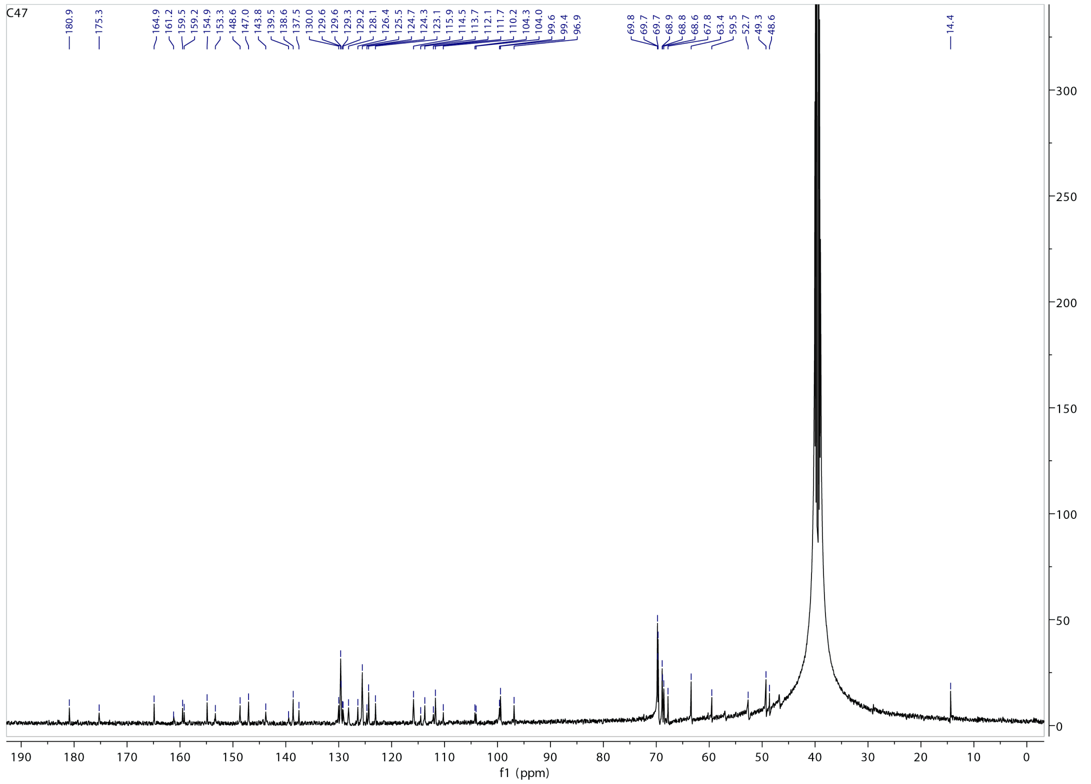

### Compound 3’

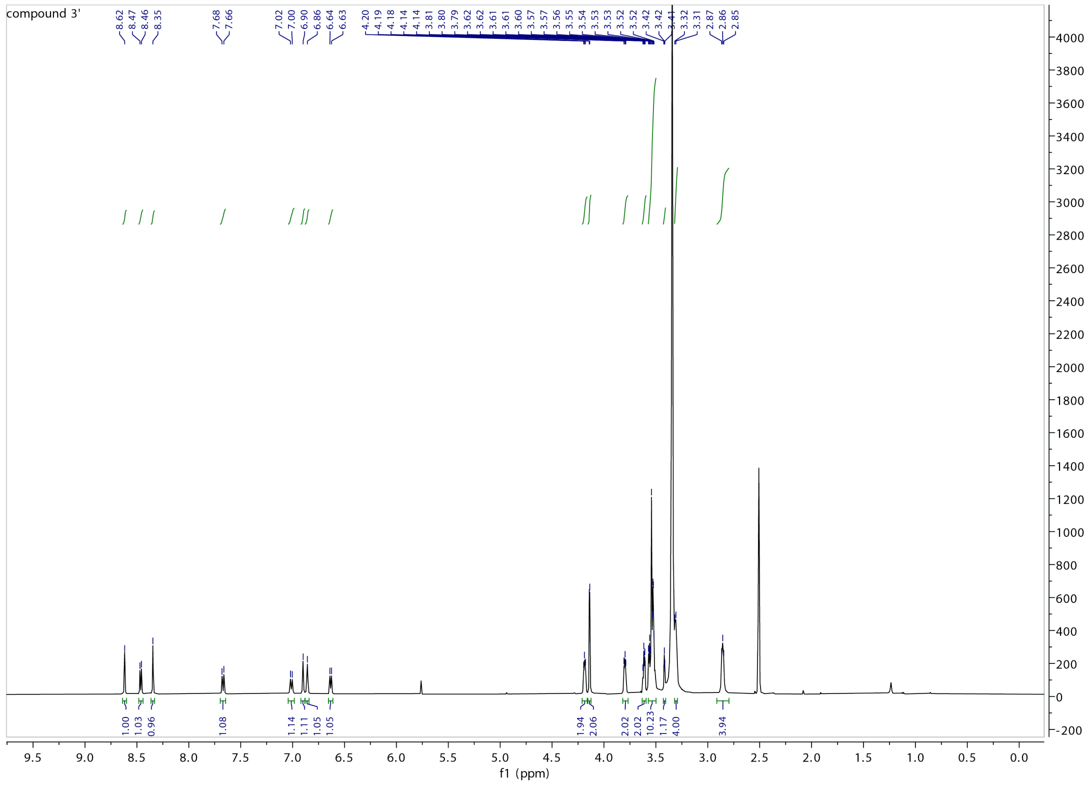


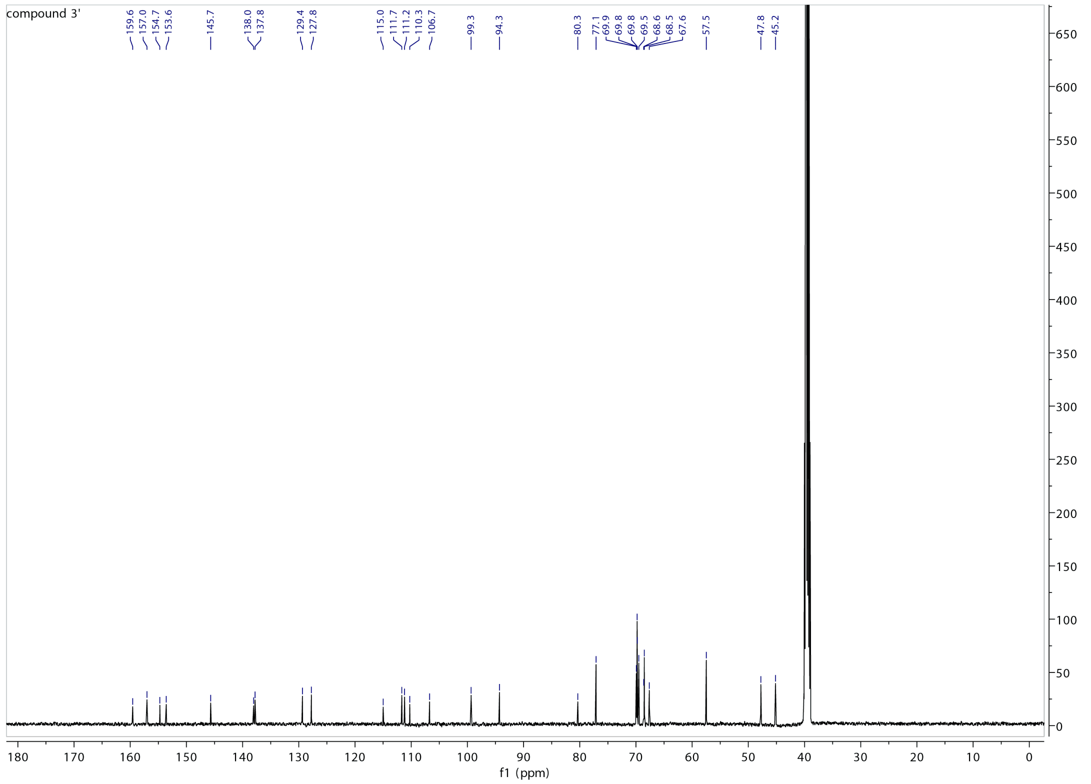

# C64

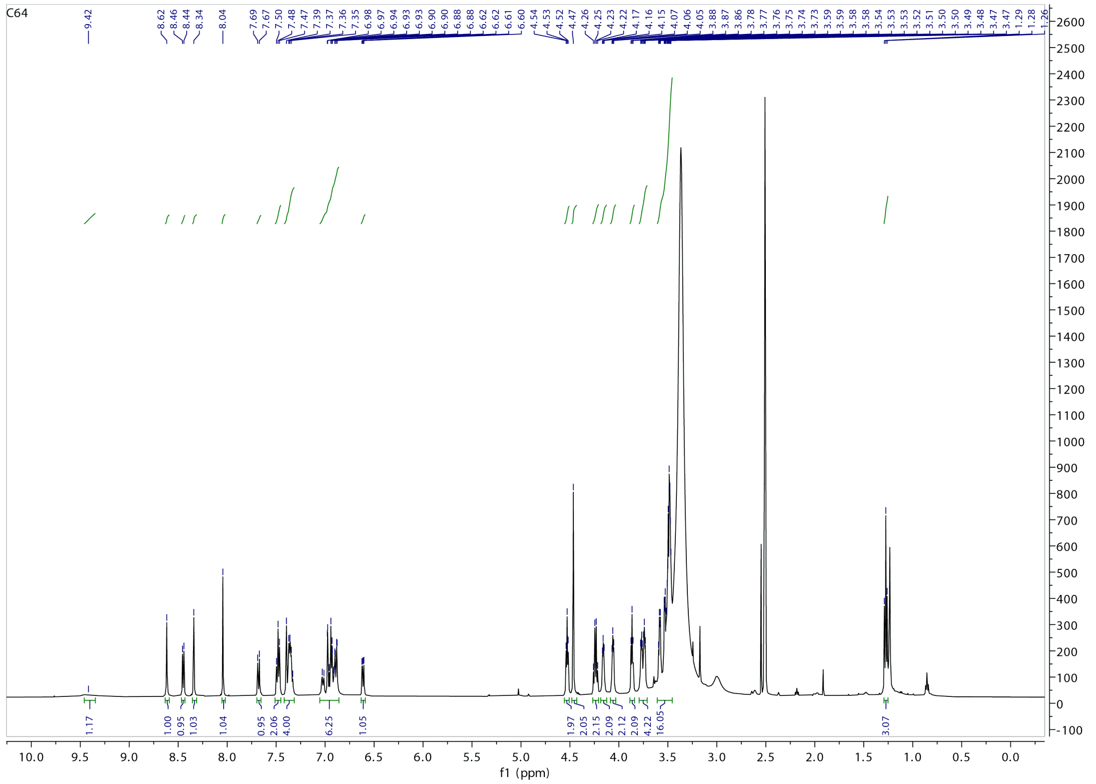
C64
9.42
8.62
8.46
8.44
8.34
8.04
7.69
7.67
7.50
7.48
7.47
7.39
7.37
7.36
7.35
6.98
6.97
6.94
6.93
6.93
6.90
6.90
6.88
6.88
6.62
6.62
6.61
6.60
4.54
4.53
4.52
4.47
4.26
4.25
4.23
4.22
4.17
4.16
4.15
4.07
4.06
4.05
3.88
3.87
3.86
3.78
3.77
3.76
3.75
3.74
3.73
3.59
3.59
3.58
3.58
3.54
3.53
3.53
3.52
3.51
3.50
3.50
3.49
3.48
3.47
3.47
1.29
1.28
1.26
1.17
1.00
0.95
1.03
1.04
0.95
2.06
4.00
6.25
1.05
1.97
2.05
2.15
2.09
2.12
2.09
4.22
16.05
3.07
f1 (ppm)

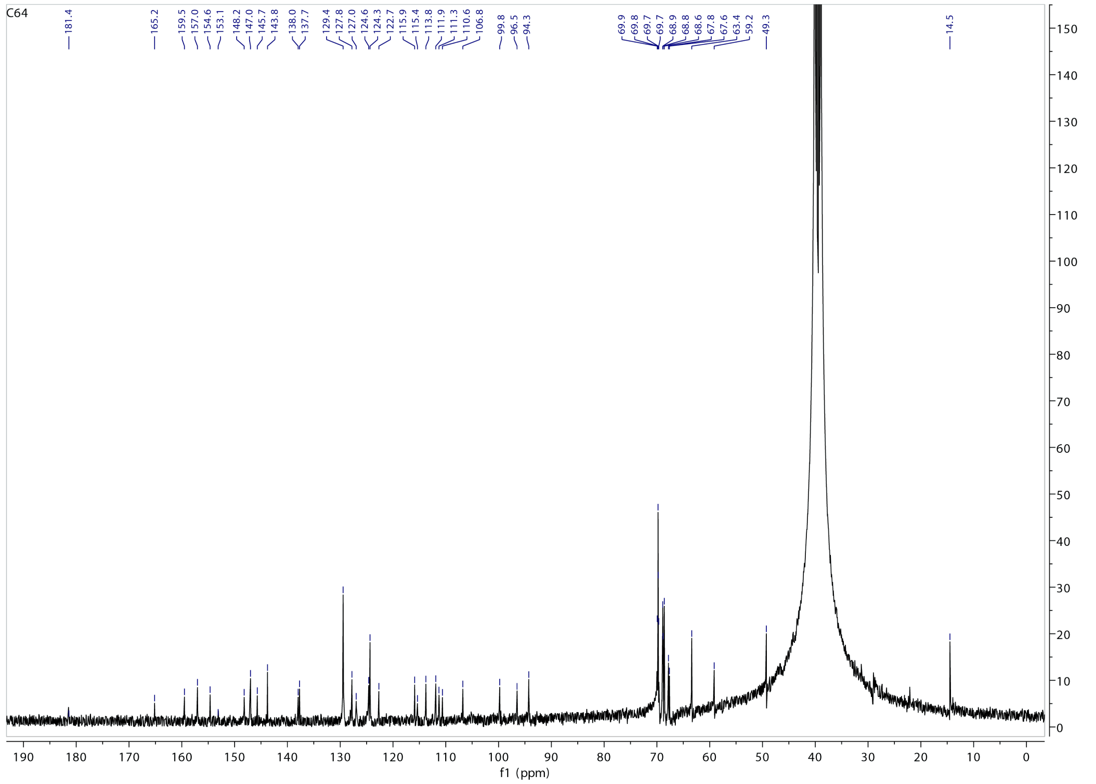
C64
181.4
165.2
159.5
157.0
154.6
153.1
148.2
147.0
145.7
143.8
138.0
137.7
129.4
127.8
127.0
124.6
124.3
122.7
115.9
115.4
113.8
111.9
111.3
110.6
106.8
99.8
96.5
94.3
69.9
69.8
69.7
68.9
68.8
68.6
67.8
67.6
63.4
59.2
49.3
14.5
f1 (ppm)

# C30-D

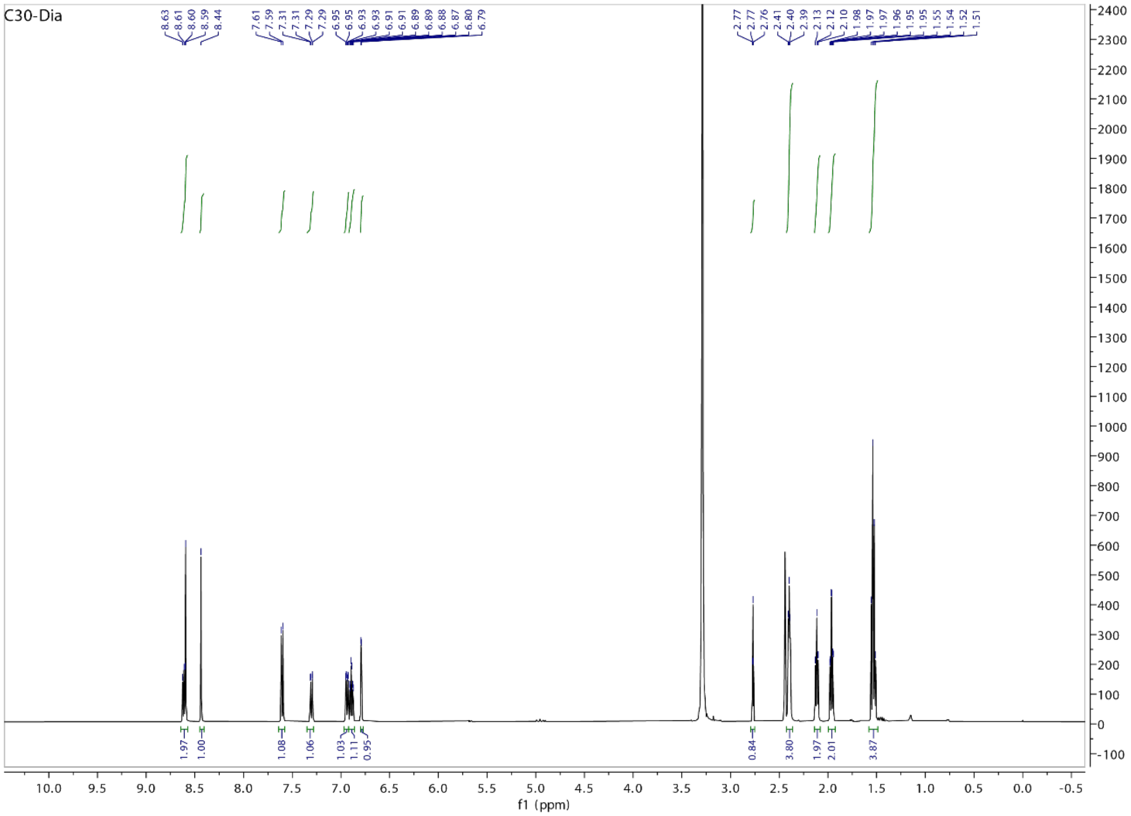
C30-Dia
8.63 8.61 8.60 8.59 8.44
7.61 7.59 7.31 7.31 7.29 7.29 6.95 6.95 6.93 6.93 6.91 6.91 6.89 6.89 6.88 6.87 6.80 6.79
2.77 2.77 2.76 2.41 2.40 2.39 2.13 2.12 2.10 1.98 1.97 1.97 1.96 1.95 1.95 1.55 1.54 1.52 1.51
1.97 1.00 1.08 1.06 1.03 1.11 0.95 0.84 3.80 1.97 2.01 3.87
f1 (ppm)

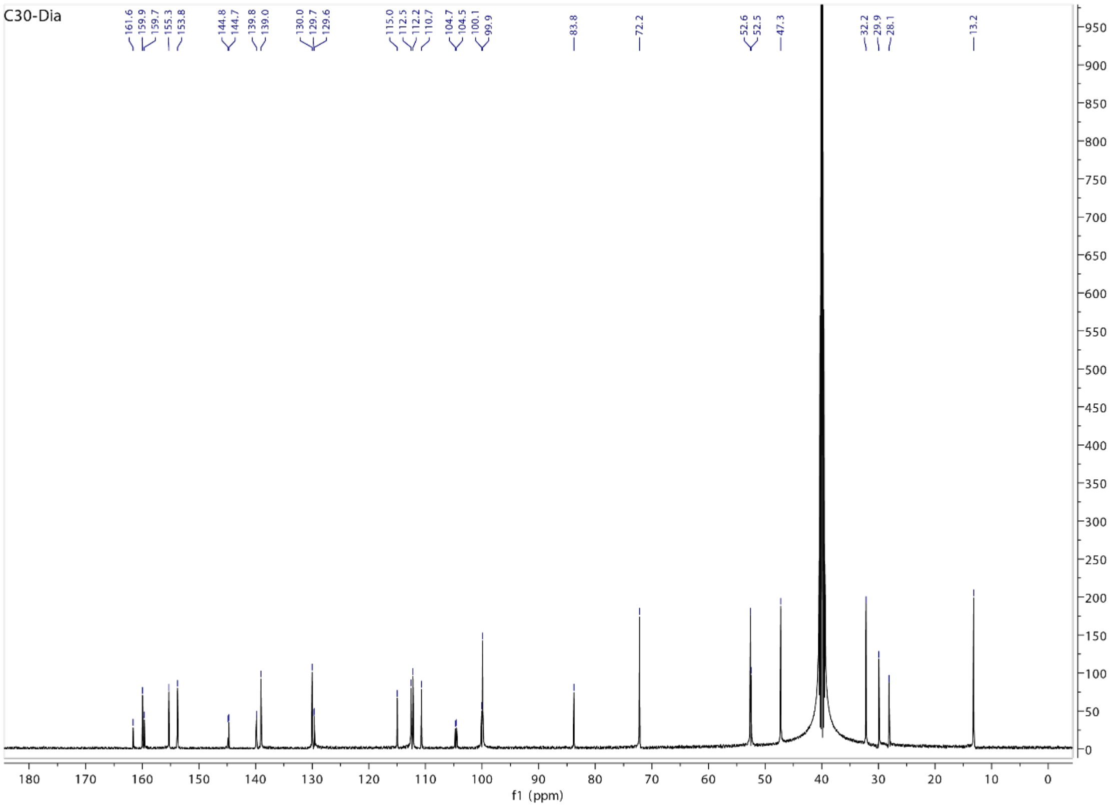
C30-Dia
161.6 159.9 159.7 155.3 153.8 144.8 144.7 139.8 139.0 130.0 129.7 129.6 115.0 112.5 112.2 110.7 104.7 104.5 100.1 99.9 83.8 72.2 52.6 52.5 47.3 32.2 29.9 28.1 13.2
f1 (ppm)

## C30-BD

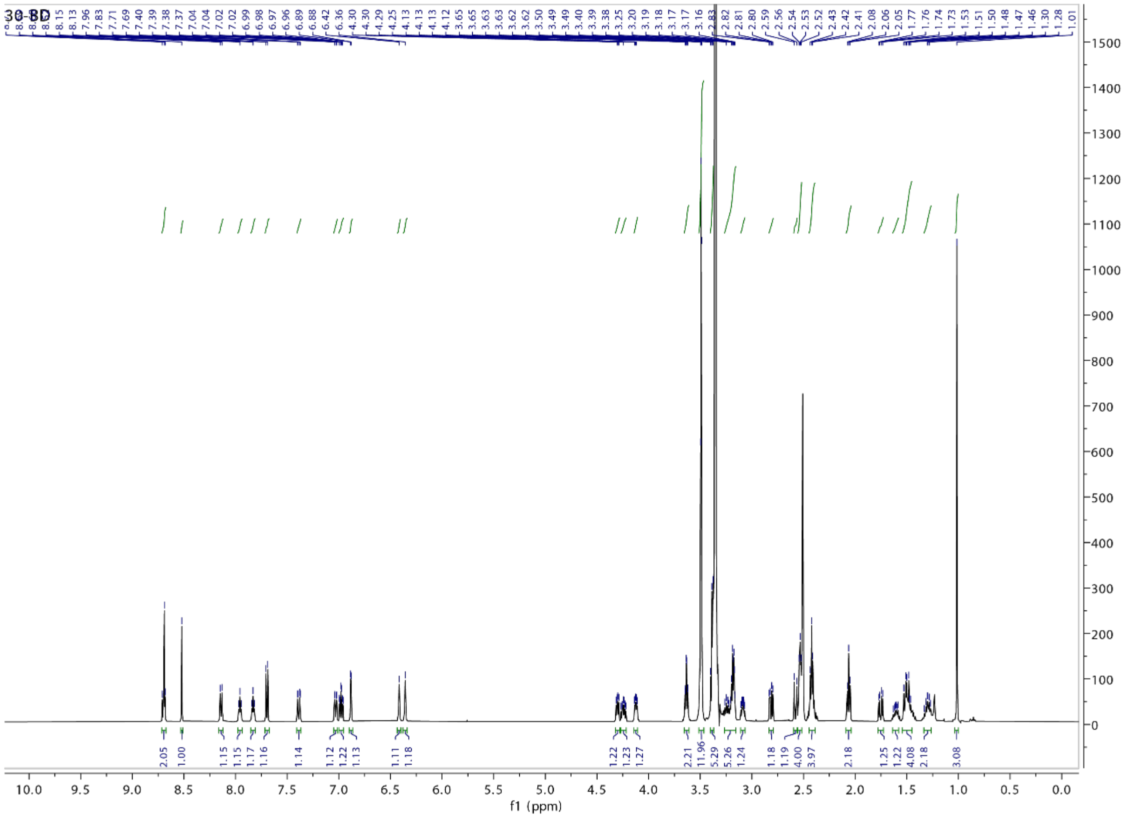

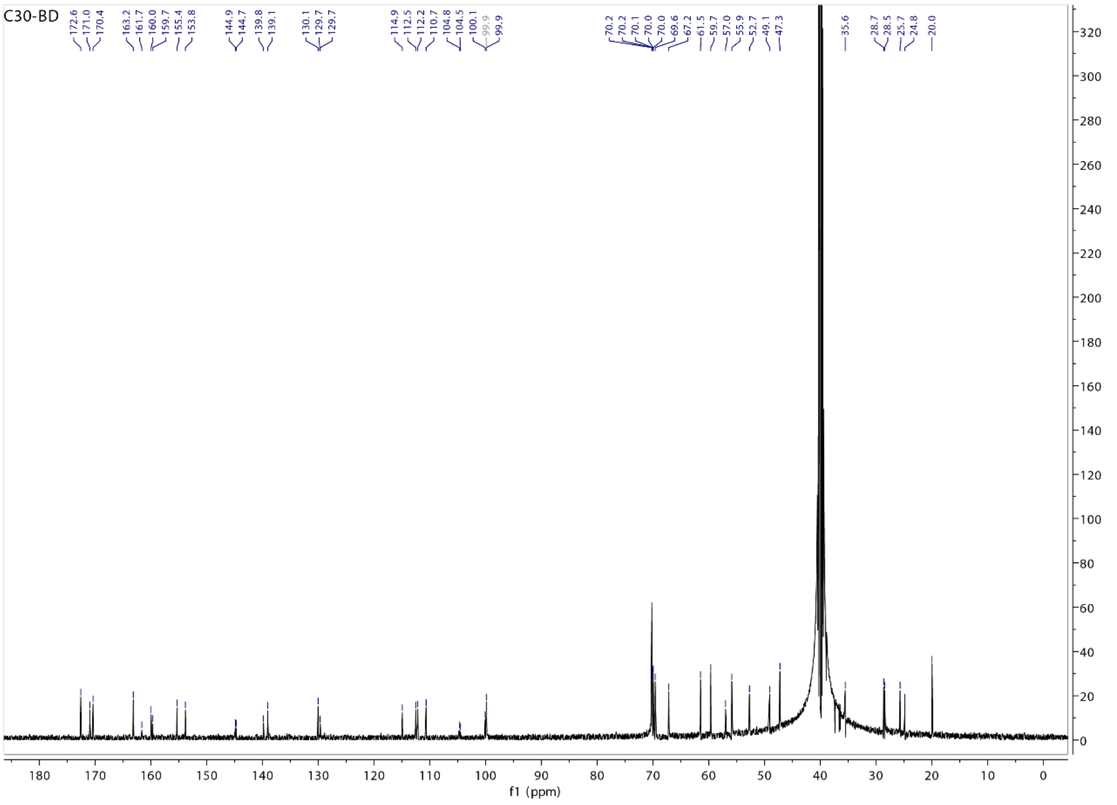

## C30-NM

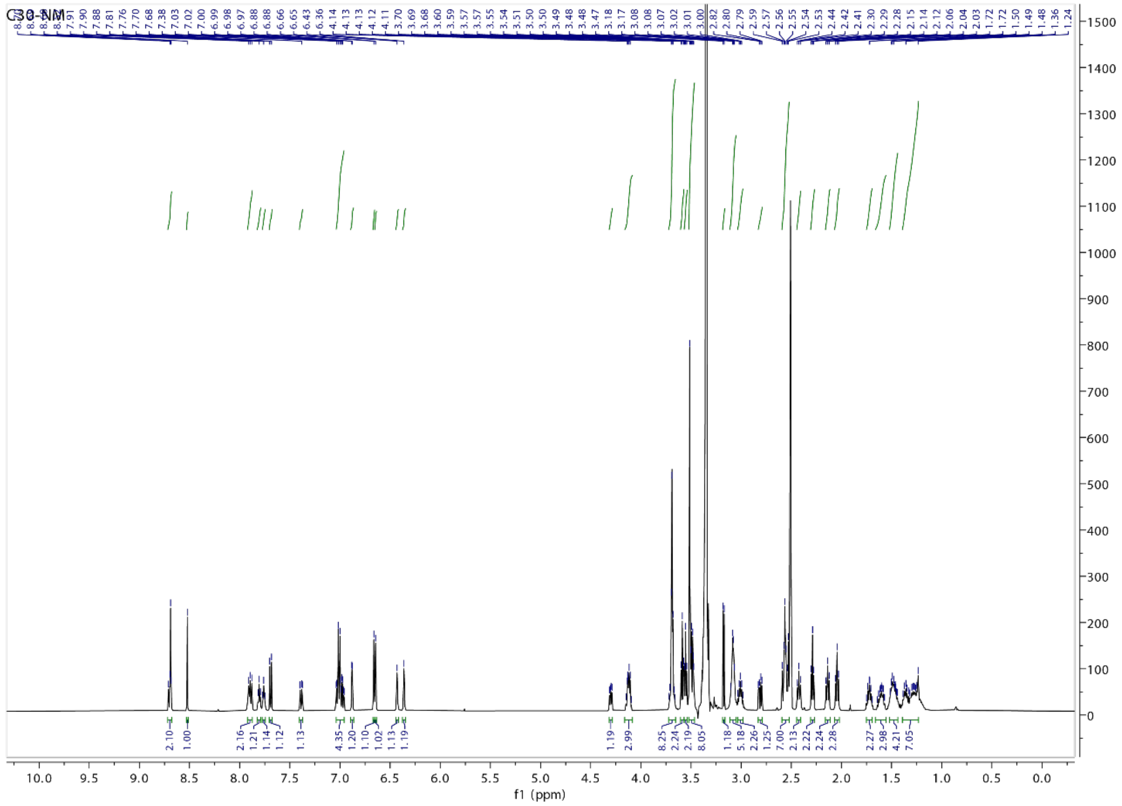


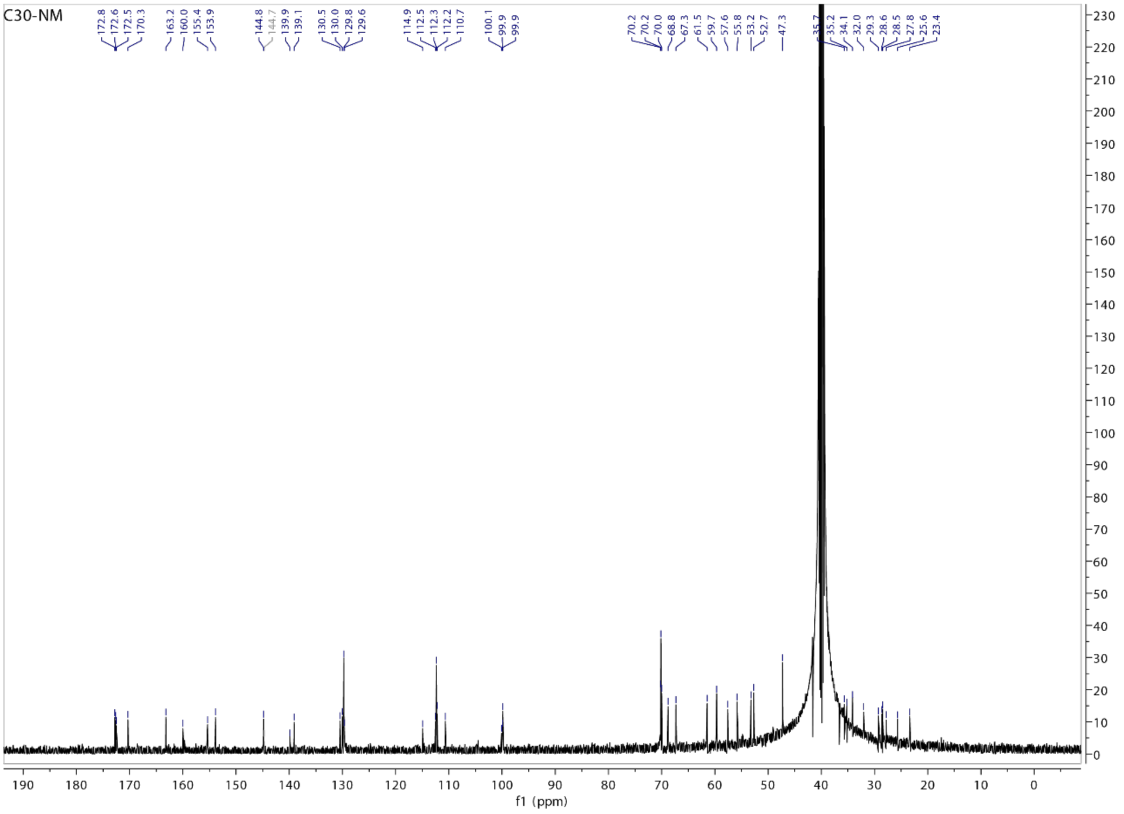

#### HPLC analysis of C30-FAI

The purity of **C30-FAI** was determined by titration with β-mercaptoethanol and analyzed by UPLC-MS. Briefly, 5 µl of the reaction solution (0.5 µmol of **C30-FAI**) was added to a 95 µl DMSO solution containing 0.4 µl β-mercaptoethanol (5.7 µmol). The mixture was incubated at room temperature for 4 h to form an adduct from **C30-FAI**. The mixture was then diluted in methanol and analyzed by UPLC-MS. Analysis shows the mixture contains ~75% **C30-FAI-β-mercaptoethanol** adduct (MS-ESI $[M+H]^+$ 806.27) and ~25% **C30-FCA** (MS-ESI $[M+H]^+$ 746.28). The stock solution was used as a **75 mM** solution.

Organic: acetonitrile; aqueous: 0.1% formic acid in water. Gradient: 0-17.5 min, 2 – 70% acetonitrile; 17.5-20 min, 70 – 98% acetonitrile.

Empower™ 3 SOFTWARE

##### SAMPLE INFORMATION

| | | | |
|---|---|---|---|
| Sample Name: | zt-4ffai-shoh | Acquired By: | System |
| Sample Type: | Unknown | Sample Set Name: | ztsm |
| Vial: | 1:F,7 | Acq. Method Set: | L3ABT2_20min |
| Injection #: | 1 | Processing Method: | 4ffai |
| Injection Volume: | 4.00 ul | Channel Name: | PDA Ch1 254nm@4.8nm |
| Run Time: | 20.0 Minutes | Proc. Chnl. Descr.: | PDA Ch1 254nm@4.8nm |

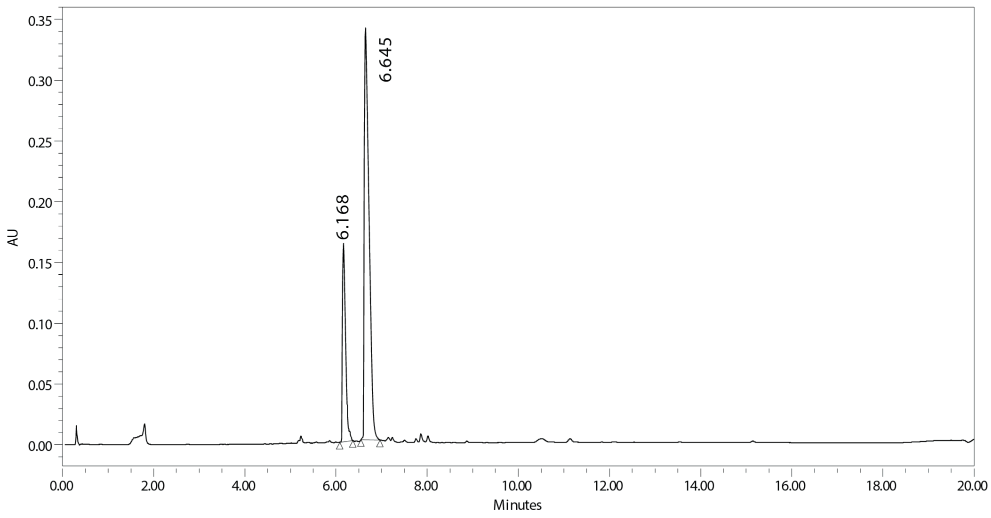


| | RT | Area | % Area | Height |
|---|---|---|---|---|
| 1 | 6.168 | 790914 | 24.19 | 163107 |
| 2 | 6.645 | 2479299 | 75.81 | 338681 |

#### HPLC analysis of C48

Organic: acetonitrile; aqueous: 0.1% ammonium formate in water. Gradient: 0-15 min, 2 – 50% acetonitrile; 15-20 min, 50 – 98% acetonitrile.

Empower™ 3 SOFTWARE

##### SAMPLE INFORMATION

| | | | |
|---|---|---|---|
| Sample Name: | c48 | Acquired By: | System |
| Sample Type: | Unknown | Sample Set Name: | ztsm |
| Vial: | 1:F,1 | Acq. Method Set: | L4ADT2_25A |
| Injection #: | 1 | Processing Method: | c48 |
| Injection Volume: | 2.00 ul | Channel Name: | PDA Ch1 254nm@4.8nm |
| Run Time: | 20.0 Minutes | Proc. Chnl. Descr.: | PDA Ch1 254nm@4.8nm |

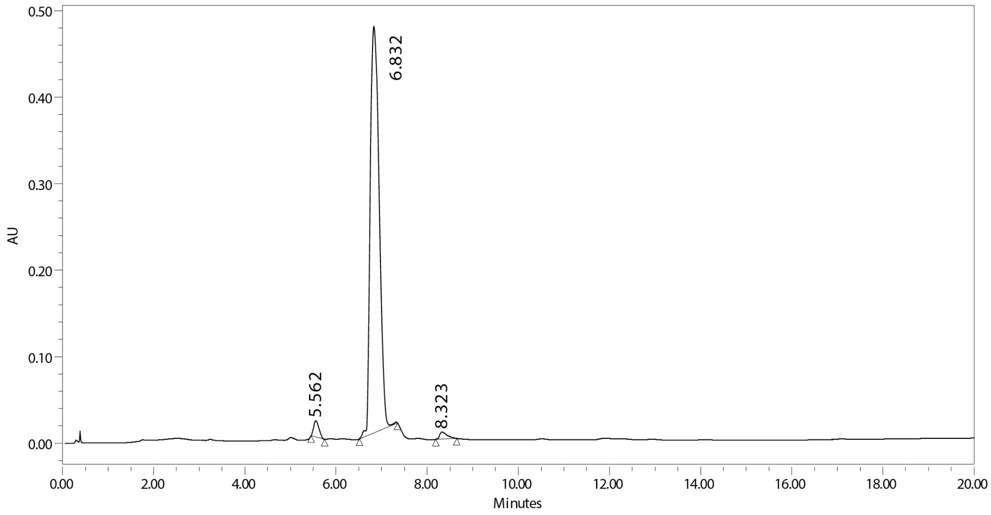


| | RT | Area | % Area | Height |
|---|---|---|---|---|
| 1 | 5.562 | 152161 | 2.27 | 18560 |
| 2 | 6.832 | 6471626 | 96.36 | 469726 |
| 3 | 8.323 | 92033 | 1.37 | 8088 |

#### HPLC analysis of C65

Organic: acetonitrile; aqueous: 0.1% ammonium formate in water. Gradient: 0-15 min, 2 – 50% acetonitrile; 15-20 min, 50 – 98% acetonitrile.

Empower™ 3 SOFTWARE

##### SAMPLE INFORMATION

| | | | |
|---|---|---|---|
| Sample Name: | c65 | Acquired By: | System |
| Sample Type: | Unknown | Sample Set Name: | ztsm |
| Vial: | 1:F,2 | Acq. Method Set: | L4ADT2_25A |
| Injection #: | 1 | Processing Method: | c65 |
| Injection Volume: | 4.00 ul | Channel Name: | PDA Ch1 254nm@4.8nm |
| Run Time: | 20.0 Minutes | Proc. Chnl. Descr.: | PDA Ch1 254nm@4.8nm |

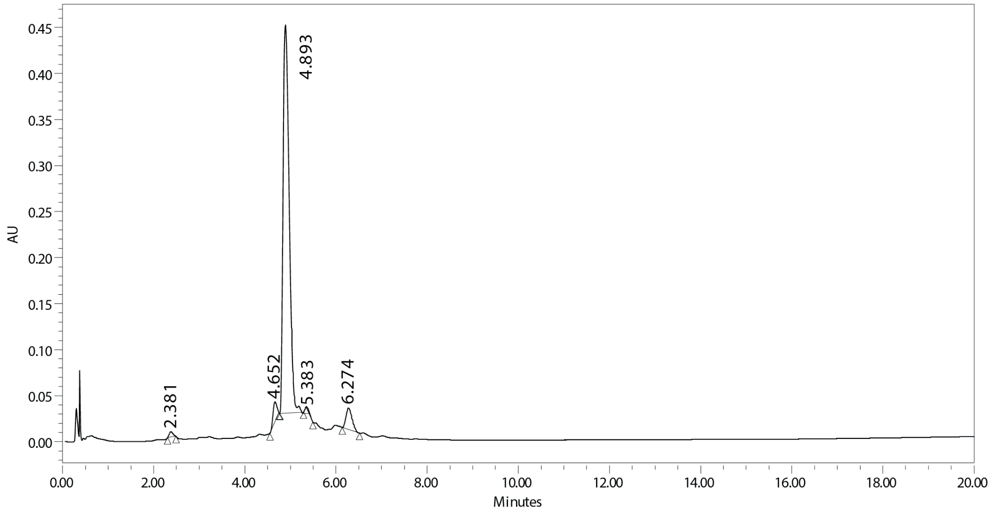


| | RT | Area | % Area | Height |
|---|---|---|---|---|
| 1 | 2.381 | 33091 | 0.76 | 5632 |
| 2 | 4.652 | 126876 | 2.93 | 22179 |
| 3 | 4.893 | 3946438 | 91.21 | 421167 |
| 4 | 5.383 | 9686 | 0.22 | 2341 |
| 5 | 6.274 | 210735 | 4.87 | 23437 |

**HPLC analysis of C30-D**

Organic: acetonitrile; aqueous: 0.1% formic acid in water. Gradient: 0-5 min, 2 – 98% acetonitrile.

SAMPLE INFORMATION

| | | | |
|---|---|---|---|
| Sample Name: | zt-4f-dia | Acquired By: | System |
| Sample Type: | Unknown | Sample Set Name: | ztsm |
| Vial: | 1:F,4 | Acq. Method Set: | L4ABT1 |
| Injection #: | 1 | Processing Method: | DIA |
| Injection Volume: | 5.00 ul | Channel Name: | PDA Ch1 254nm@4.8nm |
| Run Time: | 5.0 Minutes | Proc. Chnl. Descr.: | PDA Ch1 254nm@4.8nm |

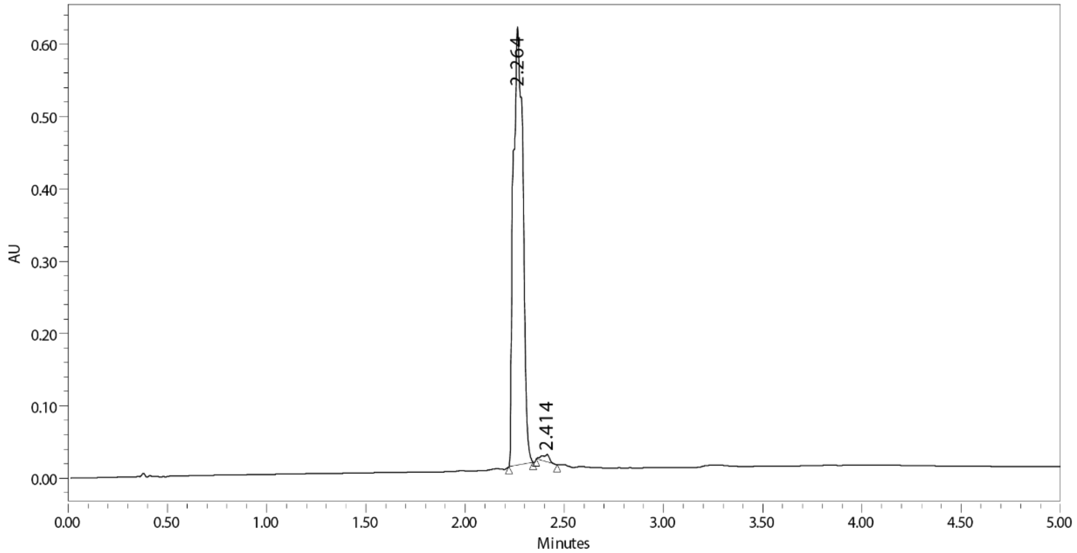


| | RT | Area | % Area | Height |
|---|---|---|---|---|
| 1 | 2.264 | 2027567 | 98.69 | 605695 |
| 2 | 2.414 | 26834 | 1.31 | 10139 |

**HPLC analysis of C30-BD**

Organic: acetonitrile; aqueous: 0.1% formic acid in water. Gradient: 0-5 min, 2 – 98% acetonitrile.

SAMPLE INFORMATION

| | | | |
|---|---|---|---|
| Sample Name: | ZT-C30-B/D | Acquired By: | System |
| Sample Type: | Unknown | Sample Set Name: | ZTSM |
| Vial: | 1:E,4 | Acq. Method Set: | L4ABT1 |
| Injection #: | 1 | Processing Method: | BD |
| Injection Volume: | 3.00 ul | Channel Name: | PDA Ch1 254nm@4.8nm |
| Run Time: | 5.0 Minutes | Proc. Chnl. Descr.: | PDA Ch1 254nm@4.8nm |

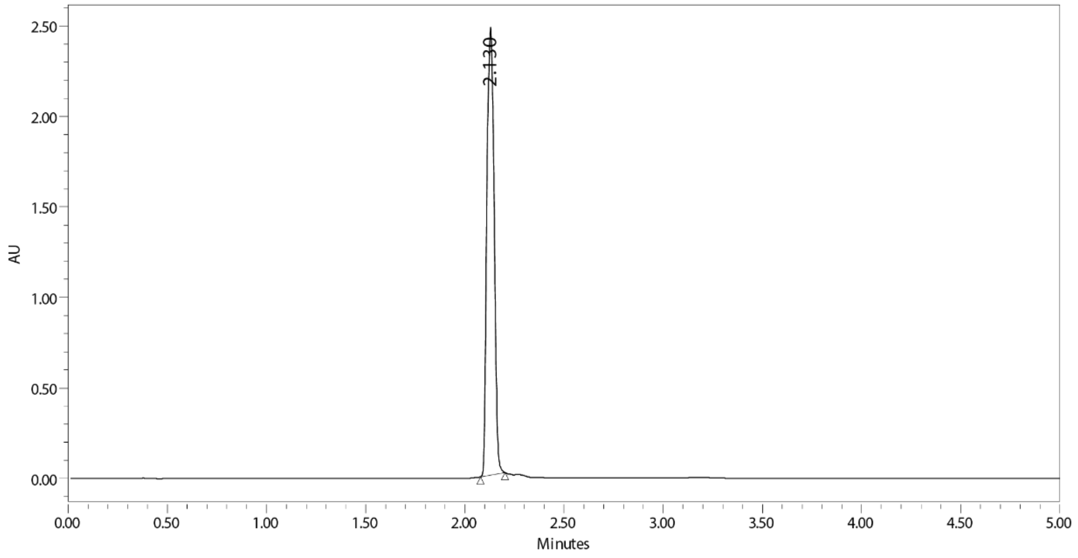


| | RT | Area | % Area | Height |
|---|---|---|---|---|
| 1 | 2.130 | 6341917 | 100.00 | 2471904 |

#### HPLC analysis of C30-NM

Organic: acetonitrile; aqueous: 0.1% formic acid in water. Gradient: 0-5 min, 2 – 98% acetonitrile.

SAMPLE INFORMATION

| | | | |
|---|---|---|---|
| Sample Name: | ZT-C30-NM | Acquired By: | System |
| Sample Type: | Unknown | Sample Set Name: | ZTSM |
| Vial: | 1:E,5 | Acq. Method Set: | L4ABT1 |
| Injection #: | 1 | Processing Method: | NM |
| Injection Volume: | 5.00 ul | Channel Name: | PDA Ch1 254nm@4.8nm |
| Run Time: | 5.0 Minutes | Proc. Chnl. Descr.: | PDA Ch1 254nm@4.8nm |

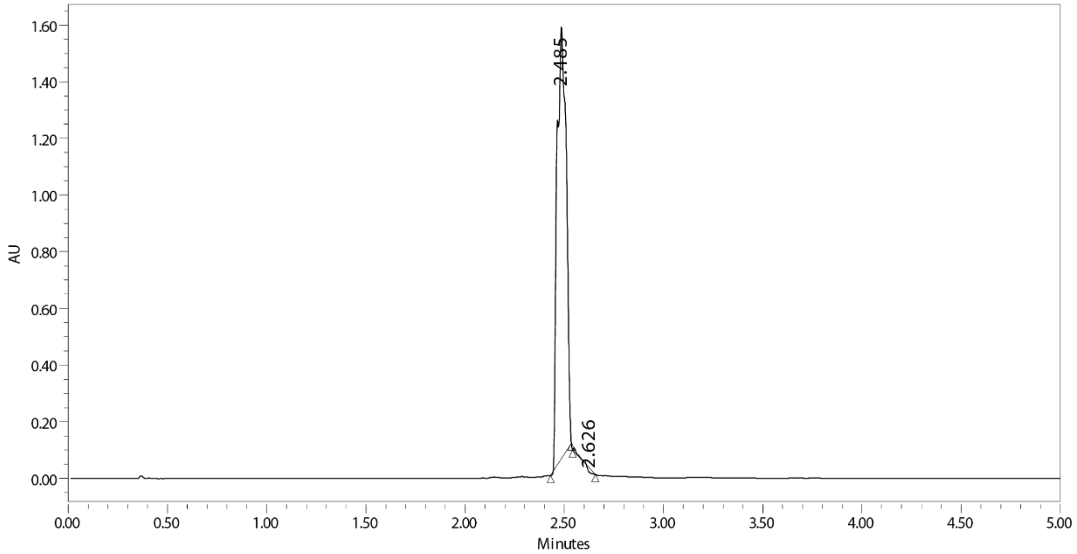


| | RT | Area | % Area | Height |
|---|---|---|---|---|
| 1 | 2.485 | 4948424 | 99.09 | 1523049 |
| 2 | 2.626 | 45478 | 0.91 | -14865 |
